## Supplementary File 1 for "Synaptic targets of photoreceptors specialized to detect color and skylight polarization in *Drosophila*"

### Supplementary File 1: Tables of all seed column R7, R8, R7-DRA and R8-DRA target cells by type

#### R7 and R8 outgoing Dm9

| name | skid | pR7a | pR7b | yR7a | yR7b | pR8a | pR8b | yR8a | yR8b | total | %R7 | %R8 | %p | %y |
|---|---|---|---|---|---|---|---|---|---|---|---|---|---|---|
| Putative Dm9 11452428 MF | 11452427 | 29 | 37 | 26 | 23 | 70 | 65 | 52 | 66 | 368 | 31.2 | 68.8 | 54.6 | 45.4 |
| Putative Dm9 11447062 CL | 11447061 | 16 | 0 | 11 | 0 | 0 | 0 | 12 | 0 | 39 | 69.2 | 30.8 | 41.0 | 59.0 |
| Putative Dm9 11450496 CL | 11450495 | 0 | 0 | 0 | 4 | 0 | 0 | 6 | 5 | 15 | 26.7 | 73.3 | 0.0 | 100.0 |
| Putative Dm9 11484680 MF | 11484679 | 8 | 0 | 0 | 0 | 0 | 0 | 0 | 0 | 8 | 100.0 | 0.0 | 100.0 | 0.0 |
| Putative Dm9 11444387 HL | 11444386 | 0 | 4 | 0 | 0 | 0 | 4 | 0 | 0 | 8 | 50.0 | 50.0 | 100.0 | 0.0 |
| Putative Dm9 11454715 CL | 11454714 | 0 | 0 | 0 | 2 | 0 | 3 | 0 | 2 | 7 | 28.6 | 71.4 | 42.9 | 57.1 |
| Total | | 53 | 41 | 37 | 29 | 70 | 72 | 70 | 73 | 445 | 36.0 | 64.0 | 53.0 | 47.0 |

#### R7 and R8 outgoing Dm8

| name | skid | pR7a | pR7b | yR7a | yR7b | pR8a | pR8b | yR8a | yR8b | total | %R7 | %R8 | %p | %y |
|---|---|---|---|---|---|---|---|---|---|---|---|---|---|---|
| Putative Dm8 11453022 CL | 11453021 | 4 | 4 | 19 | 35 | 0 | 0 | 0 | 0 | 62 | 100 | 0 | 12.9 | 87.1 |
| Putative Dm8 10411812 MF | 10411811 | 21 | 40 | 0 | 0 | 0 | 0 | 0 | 0 | 61 | 100 | 0 | 100.0 | 0.0 |
| Putative Dm8 10208776 MF | 10208775 | 15 | 3 | 35 | 6 | 0 | 0 | 0 | 0 | 59 | 100 | 0 | 30.5 | 69.5 |
| Putative Dm8 11453312 CL | 11453311 | 9 | 17 | 8 | 15 | 0 | 0 | 0 | 0 | 49 | 100 | 0 | 53.1 | 46.9 |
| Putative Dm8 10109587 MF | 10109586 | 39 | 4 | 0 | 0 | 0 | 0 | 0 | 0 | 43 | 100 | 0 | 100.0 | 0.0 |
| Putative Dm8 11500072 HL | 11500071 | 1 | 17 | 0 | 0 | 0 | 0 | 0 | 0 | 18 | 100 | 0 | 100.0 | 0.0 |
| Putative Dm8 11448877 HL | 11523807 | 0 | 17 | 0 | 0 | 0 | 0 | 0 | 0 | 17 | 100 | 0 | 100.0 | 0.0 |
| Putative Dm8 10419715 MF | 10419714 | 14 | 0 | 0 | 2 | 0 | 0 | 0 | 0 | 16 | 100 | 0 | 87.5 | 12.5 |
| Putative Dm8 11447521 MF | 11447520 | 0 | 0 | 9 | 5 | 0 | 0 | 0 | 0 | 14 | 100 | 0 | 0.0 | 100.0 |
| Putative Dm8 10410433 MF | 10410432 | 10 | 3 | 0 | 0 | 0 | 0 | 0 | 0 | 13 | 100 | 0 | 100.0 | 0.0 |
| Putative Dm8 10196190 MF | 10196189 | 10 | 0 | 3 | 0 | 0 | 0 | 0 | 0 | 13 | 100 | 0 | 76.9 | 23.1 |
| Putative Dm8 10245572 MF | 10245571 | 0 | 0 | 2 | 9 | 0 | 0 | 0 | 0 | 11 | 100 | 0 | 0.0 | 100.0 |
| Putative Dm8 10971699 MF | 10971698 | 0 | 0 | 0 | 10 | 0 | 0 | 0 | 0 | 10 | 100 | 0 | 0.0 | 100.0 |
| Putative Dm8 11445682 MF | 11445681 | 7 | 0 | 0 | 0 | 0 | 0 | 0 | 0 | 7 | 100 | 0 | 100.0 | 0.0 |
| Putative Dm8 10995249 MF | 10995248 | 0 | 0 | 0 | 3 | 0 | 0 | 0 | 0 | 3 | 100 | 0 | 0.0 | 100.0 |
| Total | | 130 | 105 | 76 | 85 | 0 | 0 | 0 | 0 | 396 | 100 | 0 | 59.3 | 40.7 |

#### R7 and R8 outgoing MeTu

| name | skid | pR7a | pR7b | yR7a | yR7b | pR8a | pR8b | yR8a | yR8b | total | %R7 | %R8 | %p | %y |
|---|---|---|---|---|---|---|---|---|---|---|---|---|---|---|
| Putative MeTu 10409864 MF | 10409863 | 22 | 13 | 17 | 12 | 0 | 0 | 0 | 0 | 64 | 100 | 0 | 54.7 | 45.3 |
| Putative MeTu 11455123 CL | 11455122 | 0 | 21 | 4 | 14 | 0 | 0 | 0 | 0 | 39 | 100 | 0 | 53.8 | 46.2 |
| Putative MeTu 11455157 CL | 11455156 | 6 | 9 | 9 | 12 | 0 | 0 | 0 | 0 | 36 | 100 | 0 | 41.7 | 58.3 |
| Putative MeTu 10409693 MF | 10409692 | 11 | 0 | 6 | 0 | 0 | 0 | 0 | 0 | 17 | 100 | 0 | 64.7 | 35.3 |
| Putative MeTu 11448396 MF | 11448395 | 0 | 0 | 8 | 4 | 0 | 0 | 0 | 0 | 12 | 100 | 0 | 0.0 | 100.0 |
| Putative MeTu 11455113 CL | 11455112 | 0 | 0 | 0 | 4 | 0 | 0 | 0 | 0 | 4 | 100 | 0 | 0.0 | 100.0 |
| Putative MeTu 11499694 CL | 11499693 | 0 | 0 | 3 | 0 | 0 | 0 | 0 | 0 | 3 | 100 | 0 | 0.0 | 100.0 |
| Total | | 39 | 43 | 47 | 46 | 0 | 0 | 0 | 0 | 175 | 100 | 0 | 46.9 | 53.1 |

#### R7 and R8 outgoing R7

| name | skid | pR7a | pR7b | yR7a | yR7b | pR8a | pR8b | yR8a | yR8b | total | %R7 | %R8 | %p | %y |
|---|---|---|---|---|---|---|---|---|---|---|---|---|---|---|
| Putative R7 col G1 10585941 MF | 10585940 | 0 | 0 | 0 | 0 | 0 | 0 | 46 | 0 | 46 | 0.0 | 100.0 | 0.0 | 100.0 |
| Putative R7 col A1 10082583 MF | 10082582 | 0 | 0 | 0 | 0 | 39 | 0 | 0 | 0 | 39 | 0.0 | 100.0 | 100.0 | 0.0 |
| Putative R7 col I1 10538511 MF | 10538510 | 0 | 0 | 0 | 0 | 0 | 36 | 0 | 0 | 36 | 0.0 | 100.0 | 100.0 | 0.0 |
| Putative R7 col H1 10653594 MF | 10653593 | 0 | 0 | 0 | 0 | 0 | 0 | 0 | 36 | 36 | 0.0 | 100.0 | 0.0 | 100.0 |
| Putative R7 10653781 MF | 10653780 | 0 | 0 | 0 | 3 | 0 | 0 | 0 | 0 | 3 | 100.0 | 0.0 | 0.0 | 100.0 |
| Total | | 0 | 0 | 0 | 3 | 39 | 36 | 46 | 36 | 160 | 1.9 | 98.1 | 46.9 | 53.1 |

#### R7 and R8 outgoing Tm5c

| name | skid | pR7a | pR7b | yR7a | yR7b | pR8a | pR8b | yR8a | yR8b | total | %R7 | %R8 | %p | %y |
|---|---|---|---|---|---|---|---|---|---|---|---|---|---|---|
| Putative Tm5c 11450506 CL | 11450505 | 0 | 0 | 0 | 2 | 0 | 0 | 21 | 22 | 45 | 4.4 | 95.6 | 0.0 | 100.0 |
| Putative Tm5c 11470103 MF | 11470102 | 0 | 0 | 1 | 0 | 0 | 0 | 23 | 0 | 24 | 4.2 | 95.8 | 0.0 | 100.0 |
| Putative Tm5c 11449561 MF | 11449560 | 2 | 5 | 0 | 0 | 6 | 4 | 3 | 4 | 24 | 29.2 | 70.8 | 70.8 | 29.2 |
| Putative Tm5c 11574444 MF | 11574443 | 0 | 0 | 2 | 0 | 0 | 0 | 16 | 0 | 18 | 11.1 | 88.9 | 0.0 | 100.0 |
| Putative Tm5c 11473479 CL | 11473478 | 0 | 0 | 0 | 4 | 0 | 0 | 0 | 14 | 18 | 22.2 | 77.8 | 0.0 | 100.0 |
| Putative Tm5c 11485095 MF | 11485094 | 1 | 0 | 0 | 0 | 10 | 0 | 0 | 0 | 11 | 9.1 | 90.9 | 100.0 | 0.0 |
| Total | | 3 | 5 | 3 | 6 | 16 | 4 | 63 | 40 | 140 | 12.1 | 87.9 | 20.0 | 80.0 |

#### R7 and R8 outgoing Tm20

| name | skid | pR7a | pR7b | yR7a | yR7b | pR8a | pR8b | yR8a | yR8b | total | %R7 | %R8 | %p | %y |
|---|---|---|---|---|---|---|---|---|---|---|---|---|---|---|
| Putative Tm20 col I1 11444393 HL | 11444392 | 0 | 2 | 0 | 0 | 0 | 35 | 0 | 0 | 37 | 5.4 | 94.6 | 100.0 | 0.0 |
| Putative Tm20 col H1 11450553 CL | 11450552 | 0 | 0 | 0 | 2 | 0 | 0 | 0 | 35 | 37 | 5.4 | 94.6 | 0.0 | 100.0 |
| Putative Tm20 col A1 10423775 MF | 10423774 | 3 | 0 | 0 | 0 | 32 | 0 | 0 | 0 | 35 | 8.6 | 91.4 | 100.0 | 0.0 |
| Putative Tm20 col G1 11473669 MF | 11473668 | 0 | 0 | 1 | 0 | 0 | 0 | 30 | 0 | 31 | 3.2 | 96.8 | 0.0 | 100.0 |
| Total | | 3 | 2 | 1 | 2 | 32 | 35 | 30 | 35 | 140 | 5.7 | 94.3 | 51.4 | 48.6 |

#### R7 and R8 outgoing Mi15

| name | skid | pR7a | pR7b | yR7a | yR7b | pR8a | pR8b | yR8a | yR8b | total | %R7 | %R8 | %p | %y |
|---|---|---|---|---|---|---|---|---|---|---|---|---|---|---|
| Putative Mi15 11445262 MF | 11445261 | 5 | 0 | 2 | 0 | 13 | 0 | 22 | 0 | 42 | 16.7 | 83.3 | 42.9 | 57.1 |
| Putative Mi15 11450568 CL | 11450567 | 0 | 0 | 0 | 3 | 0 | 0 | 0 | 32 | 35 | 8.6 | 91.4 | 0.0 | 100.0 |
| Putative Mi15 11453819 CL | 11453818 | 0 | 0 | 2 | 0 | 0 | 0 | 25 | 2 | 29 | 6.9 | 93.1 | 0.0 | 100.0 |
| Putative Mi15 11484381 HL | 11484380 | 0 | 1 | 0 | 0 | 0 | 25 | 0 | 0 | 26 | 3.8 | 96.2 | 100.0 | 0.0 |
| Total | | 5 | 1 | 4 | 3 | 13 | 25 | 47 | 34 | 132 | 9.8 | 90.2 | 33.3 | 66.7 |

#### R7 and R8 outgoing Mi4

| name | skid | pR7a | pR7b | yR7a | yR7b | pR8a | pR8b | yR8a | yR8b | total | %R7 | %R8 | %p | %y |
|---|---|---|---|---|---|---|---|---|---|---|---|---|---|---|
| Putative Mi4 col G1 11481479 MF | 11481478 | 0 | 0 | 0 | 0 | 0 | 0 | 30 | 0 | 30 | 0 | 100 | 0 | 100 |
| Putative Mi4 col I1 11467500 HL | 11467499 | 0 | 0 | 0 | 0 | 0 | 29 | 0 | 0 | 29 | 0 | 100 | 100 | 0 |
| Putative Mi4 col A1 11465885 CL | 11465884 | 0 | 0 | 0 | 0 | 29 | 0 | 0 | 0 | 29 | 0 | 100 | 100 | 0 |
| Putative Mi4 col H1 11473654 CL | 11473653 | 0 | 0 | 0 | 0 | 0 | 0 | 0 | 28 | 28 | 0 | 100 | 0 | 100 |
| Total | | 0 | 0 | 0 | 0 | 29 | 29 | 30 | 28 | 116 | 0 | 100 | 50 | 50 |

#### R7 and R8 outgoing ML1

| name | skid | pR7a | pR7b | yR7a | yR7b | pR8a | pR8b | yR8a | yR8b | total | %R7 | %R8 | %p | %y |
|---|---|---|---|---|---|---|---|---|---|---|---|---|---|---|
| Putative ML1 11472158 CL | 11472157 | 0 | 0 | 0 | 0 | 11 | 4 | 13 | 13 | 41 | 0 | 100 | 36.6 | 63.4 |
| Putative ML1 11458491 CL | 11458490 | 0 | 0 | 0 | 0 | 13 | 12 | 4 | 0 | 29 | 0 | 100 | 86.2 | 13.8 |
| Putative ML1 11471220 CL | 11471219 | 0 | 0 | 0 | 0 | 0 | 13 | 4 | 7 | 24 | 0 | 100 | 54.2 | 45.8 |
| Putative ML1 11458827 CL | 11458826 | 0 | 0 | 0 | 0 | 5 | 0 | 0 | 0 | 5 | 0 | 100 | 100.0 | 0.0 |
| Total | | 0 | 0 | 0 | 0 | 29 | 29 | 21 | 20 | 99 | 0 | 100 | 58.6 | 41.4 |

#### R7 and R8 outgoing Dm2

| name | skid | pR7a | pR7b | yR7a | yR7b | pR8a | pR8b | yR8a | yR8b | total | %R7 | %R8 | %p | %y |
|---|---|---|---|---|---|---|---|---|---|---|---|---|---|---|
| Putative Dm2 11448823 HL | 11448822 | 0 | 12 | 0 | 0 | 0 | 24 | 0 | 0 | 36 | 33.3 | 66.7 | 100.0 | 0.0 |
| Putative Dm2 10499250 MF | 10499249 | 6 | 0 | 0 | 0 | 29 | 0 | 0 | 0 | 35 | 17.1 | 82.9 | 100.0 | 0.0 |
| Putative Dm2 11453278 CL | 11453277 | 0 | 0 | 0 | 3 | 0 | 0 | 0 | 15 | 18 | 16.7 | 83.3 | 0.0 | 100.0 |
| Putative Dm2 11466217 CL | 11466216 | 0 | 0 | 0 | 0 | 7 | 0 | 0 | 0 | 7 | 0.0 | 100.0 | 100.0 | 0.0 |
| Total | | 6 | 12 | 0 | 3 | 36 | 24 | 0 | 15 | 96 | 21.9 | 78.1 | 81.2 | 18.8 |

#### R7 and R8 outgoing Dm11

| name | skid | pR7a | pR7b | yR7a | yR7b | pR8a | pR8b | yR8a | yR8b | total | %R7 | %R8 | %p | %y |
|---|---|---|---|---|---|---|---|---|---|---|---|---|---|---|
| Putative Dm11 11450454 CL | 11450453 | 15 | 4 | 9 | 18 | 2 | 0 | 3 | 1 | 52 | 88.5 | 11.5 | 40.4 | 59.6 |
| Putative Dm11 11444399 HL | 11444398 | 0 | 9 | 0 | 0 | 0 | 2 | 0 | 0 | 11 | 81.8 | 18.2 | 100.0 | 0.0 |
| Total | | 15 | 13 | 9 | 18 | 2 | 2 | 3 | 1 | 63 | 87.3 | 12.7 | 50.8 | 49.2 |

#### R7 and R8 outgoing L3

| name | skid | pR7a | pR7b | yR7a | yR7b | pR8a | pR8b | yR8a | yR8b | total | %R7 | %R8 | %p | %y |
|---|---|---|---|---|---|---|---|---|---|---|---|---|---|---|
| Putative L3 col H1 11450470 CL | 11450469 | 0 | 0 | 0 | 12 | 0 | 0 | 0 | 7 | 19 | 63.2 | 36.8 | 0.0 | 100.0 |
| Putative L3 col A1 11445252 MF | 11445251 | 10 | 0 | 0 | 0 | 6 | 0 | 0 | 0 | 16 | 62.5 | 37.5 | 100.0 | 0.0 |
| Putative L3 col I1 11448597 HL | 11448596 | 0 | 9 | 0 | 0 | 0 | 6 | 0 | 0 | 15 | 60.0 | 40.0 | 100.0 | 0.0 |
| Putative L3 col G1 11453914 CL | 11453913 | 0 | 0 | 5 | 0 | 0 | 0 | 6 | 0 | 11 | 45.5 | 54.5 | 0.0 | 100.0 |
| Total | | 10 | 9 | 5 | 12 | 6 | 6 | 6 | 7 | 61 | 59.0 | 41.0 | 50.8 | 49.2 |

#### R7 and R8 outgoing Mi1

| name | skid | pR7a | pR7b | yR7a | yR7b | pR8a | pR8b | yR8a | yR8b | total | %R7 | %R8 | %p | %y |
|---|---|---|---|---|---|---|---|---|---|---|---|---|---|---|
| Putative Mi1 col I1 11471702 HL | 11471701 | 0 | 0 | 0 | 0 | 0 | 17 | 0 | 0 | 17 | 0 | 100 | 100.0 | 0.0 |
| Putative Mi1 col H1 11472781 CL | 11472780 | 0 | 0 | 0 | 0 | 0 | 0 | 0 | 14 | 14 | 0 | 100 | 0.0 | 100.0 |
| Putative Mi1 col G1 11470015 MF | 11470014 | 0 | 0 | 0 | 0 | 0 | 0 | 14 | 0 | 14 | 0 | 100 | 0.0 | 100.0 |
| Putative Mi1 col A1 11458128 CL | 11458127 | 0 | 0 | 0 | 0 | 14 | 0 | 0 | 0 | 14 | 0 | 100 | 100.0 | 0.0 |
| Total | | 0 | 0 | 0 | 0 | 14 | 17 | 14 | 14 | 59 | 0 | 100 | 52.5 | 47.5 |

#### R7 and R8 outgoing R8

| name | skid | pR7a | pR7b | yR7a | yR7b | pR8a | pR8b | yR8a | yR8b | total | %R7 | %R8 | %p | %y |
|---|---|---|---|---|---|---|---|---|---|---|---|---|---|---|
| Putative R8 col A1 10086692 MF | 10086691 | 20 | 0 | 0 | 0 | 1 | 0 | 0 | 0 | 21 | 95.2 | 4.8 | 100.0 | 0.0 |
| Putative R8 col H1 11468319 CL | 11468318 | 0 | 0 | 0 | 19 | 0 | 0 | 0 | 0 | 19 | 100.0 | 0.0 | 0.0 | 100.0 |
| Putative R8 col I1 11466409 HL | 11466408 | 0 | 11 | 0 | 0 | 0 | 0 | 0 | 0 | 11 | 100.0 | 0.0 | 100.0 | 0.0 |
| Putative R8 col G1 10629255 MF | 10629254 | 0 | 0 | 7 | 0 | 0 | 0 | 0 | 0 | 7 | 100.0 | 0.0 | 0.0 | 100.0 |
| Total | | 20 | 11 | 7 | 19 | 1 | 0 | 0 | 0 | 58 | 98.3 | 1.7 | 55.2 | 44.8 |

#### R7 and R8 outgoing Tm5a

| name | skid | pR7a | pR7b | yR7a | yR7b | pR8a | pR8b | yR8a | yR8b | total | %R7 | %R8 | %p | %y |
|---|---|---|---|---|---|---|---|---|---|---|---|---|---|---|
| Putative Tm5a 11447184 MF | 11447183 | 0 | 0 | 29 | 0 | 0 | 0 | 0 | 0 | 29 | 100 | 0 | 0 | 100 |
| Putative Tm5a 11453107 CL | 11453106 | 0 | 0 | 0 | 24 | 0 | 0 | 0 | 0 | 24 | 100 | 0 | 0 | 100 |
| Total | | 0 | 0 | 29 | 24 | 0 | 0 | 0 | 0 | 53 | 100 | 0 | 0 | 100 |

#### R7 and R8 outgoing Tm5b

| name | skid | pR7a | pR7b | yR7a | yR7b | pR8a | pR8b | yR8a | yR8b | total | %R7 | %R8 | %p | %y |
|---|---|---|---|---|---|---|---|---|---|---|---|---|---|---|
| Putative Tm5b 11448828 HL | 11448827 | 0 | 19 | 0 | 0 | 0 | 1 | 0 | 6 | 26 | 73.1 | 26.9 | 76.9 | 23.1 |
| Putative Tm5b 10356413 MF | 10356412 | 20 | 0 | 0 | 0 | 2 | 0 | 0 | 0 | 22 | 90.9 | 9.1 | 100.0 | 0.0 |
| Total | | 20 | 19 | 0 | 0 | 2 | 1 | 0 | 6 | 48 | 81.2 | 18.8 | 87.5 | 12.5 |

#### R7 and R8 outgoing Tm

| name | skid | pR7a | pR7b | yR7a | yR7b | pR8a | pR8b | yR8a | yR8b | total | %R7 | %R8 | %p | %y |
|---|---|---|---|---|---|---|---|---|---|---|---|---|---|---|
| Putative Tm 11544671 HL | 11544670 | 0 | 10 | 0 | 0 | 0 | 0 | 0 | 0 | 10 | 100.0 | 0.0 | 100.0 | 0.0 |
| Putative Tm 11671250 MF | 11671249 | 0 | 0 | 0 | 0 | 0 | 0 | 0 | 7 | 7 | 0.0 | 100.0 | 0.0 | 100.0 |
| Putative Tm 10692408 HL | 10692407 | 0 | 0 | 0 | 0 | 0 | 0 | 3 | 2 | 5 | 0.0 | 100.0 | 0.0 | 100.0 |
| Putative Tm 11474296 CL | 11474295 | 0 | 0 | 0 | 0 | 0 | 0 | 0 | 4 | 4 | 0.0 | 100.0 | 0.0 | 100.0 |
| Putative Tm 11458217 CL | 11458216 | 0 | 0 | 0 | 0 | 4 | 0 | 0 | 0 | 4 | 0.0 | 100.0 | 100.0 | 0.0 |
| Putative Tm 11455044 MF | 11455043 | 0 | 0 | 0 | 4 | 0 | 0 | 0 | 0 | 4 | 100.0 | 0.0 | 0.0 | 100.0 |
| Putative Tm 11445921 MF | 11445920 | 3 | 0 | 0 | 0 | 1 | 0 | 0 | 0 | 4 | 75.0 | 25.0 | 100.0 | 0.0 |
| Putative Tm 11459161 CL | 11459160 | 1 | 0 | 0 | 0 | 2 | 0 | 0 | 0 | 3 | 33.3 | 66.7 | 100.0 | 0.0 |
| Putative Tm 11450248 HL | 11450247 | 0 | 3 | 0 | 0 | 0 | 0 | 0 | 0 | 3 | 100.0 | 0.0 | 100.0 | 0.0 |
| Total | | 4 | 13 | 0 | 4 | 7 | 0 | 3 | 13 | 44 | 47.7 | 52.3 | 54.5 | 45.5 |

#### R7 and R8 outgoing Tm5b-like

| name | skid | pR7a | pR7b | yR7a | yR7b | pR8a | pR8b | yR8a | yR8b | total | %R7 | %R8 | %p | %y |
|---|---|---|---|---|---|---|---|---|---|---|---|---|---|---|
| Putative Tm5b-like 11447921 MF | 11447920 | 0 | 0 | 7 | 0 | 0 | 0 | 16 | 0 | 23 | 30.4 | 69.6 | 0 | 100 |
| Putative Tm5b-like 11447511 CL | 11447510 | 0 | 0 | 4 | 0 | 0 | 0 | 7 | 0 | 11 | 36.4 | 63.6 | 0 | 100 |
| Putative Tm5b-like 11468647 CL | 11468646 | 0 | 0 | 2 | 1 | 0 | 0 | 0 | 4 | 7 | 42.9 | 57.1 | 0 | 100 |
| Total | | 0 | 0 | 13 | 1 | 0 | 0 | 23 | 4 | 41 | 34.1 | 65.9 | 0 | 100 |

#### R7 and R8 outgoing Mi9

| name | skid | pR7a | pR7b | yR7a | yR7b | pR8a | pR8b | yR8a | yR8b | total | %R7 | %R8 | %p | %y |
|---|---|---|---|---|---|---|---|---|---|---|---|---|---|---|
| Putative Mi9 col A1 10422285 MF | 10422284 | 3 | 0 | 0 | 0 | 12 | 0 | 0 | 0 | 15 | 20.0 | 80.0 | 100.0 | 0.0 |
| Putative Mi9 col I1 11467629 HL | 11467628 | 0 | 0 | 0 | 0 | 0 | 14 | 0 | 0 | 14 | 0.0 | 100.0 | 100.0 | 0.0 |
| Putative Mi9 col G1 11447431 CL | 11447430 | 0 | 0 | 5 | 0 | 0 | 0 | 1 | 0 | 6 | 83.3 | 16.7 | 0.0 | 100.0 |
| Putative Mi9 col H1 11453112 CL | 11453111 | 0 | 0 | 0 | 4 | 0 | 0 | 0 | 0 | 4 | 100.0 | 0.0 | 0.0 | 100.0 |
| Total | | 3 | 0 | 5 | 4 | 12 | 14 | 1 | 0 | 39 | 30.8 | 69.2 | 74.4 | 25.6 |

#### R7 and R8 outgoing L1

| name | skid | pR7a | pR7b | yR7a | yR7b | pR8a | pR8b | yR8a | yR8b | total | %R7 | %R8 | %p | %y |
|---|---|---|---|---|---|---|---|---|---|---|---|---|---|---|
| Putative L1 col I1 11472941 HL | 11472940 | 0 | 0 | 0 | 0 | 0 | 13 | 0 | 0 | 13 | 0.0 | 100.0 | 100.0 | 0.0 |
| Putative L1 col A1 10108986 MF | 10108985 | 2 | 0 | 0 | 0 | 8 | 0 | 0 | 0 | 10 | 20.0 | 80.0 | 100.0 | 0.0 |
| Putative L1 col H1 11472771 CL | 11472770 | 0 | 0 | 0 | 1 | 0 | 0 | 0 | 8 | 9 | 11.1 | 88.9 | 0.0 | 100.0 |
| Putative L1 col G1 11470179 MF | 11470178 | 0 | 0 | 0 | 0 | 0 | 0 | 6 | 0 | 6 | 0.0 | 100.0 | 0.0 | 100.0 |
| Total | | 2 | 0 | 0 | 1 | 8 | 13 | 6 | 8 | 38 | 7.9 | 92.1 | 60.5 | 39.5 |

#### R7 and R8 outgoing aMe12

| name | skid | pR7a | pR7b | yR7a | yR7b | pR8a | pR8b | yR8a | yR8b | total | %R7 | %R8 | %p | %y |
|---|---|---|---|---|---|---|---|---|---|---|---|---|---|---|
| Putative aMe12 VPN ME.R to vACA OLCT bilateral 7038036 JS ECM | 7038035 | 2 | 0 | 0 | 0 | 15 | 0 | 0 | 0 | 17 | 11.8 | 88.2 | 100 | 0 |
| Putative aMe12 VPN ME.R to vACA OLCT bilateral 28842 AA GA | 28841 | 1 | 1 | 0 | 0 | 0 | 11 | 0 | 0 | 13 | 15.4 | 84.6 | 100 | 0 |
| Total | | 3 | 1 | 0 | 0 | 15 | 11 | 0 | 0 | 30 | 13.3 | 86.7 | 100 | 0 |

#### R7 and R8 outgoing Dm

| name | skid | pR7a | pR7b | yR7a | yR7b | pR8a | pR8b | yR8a | yR8b | total | %R7 | %R8 | %p | %y |
|---|---|---|---|---|---|---|---|---|---|---|---|---|---|---|
| Putative Dm 11511058 MF | 11511057 | 0 | 4 | 0 | 0 | 0 | 4 | 0 | 0 | 8 | 50.0 | 50.0 | 100.0 | 0.0 |
| Putative Dm 11448963 MF | 11448962 | 2 | 4 | 0 | 0 | 0 | 2 | 0 | 0 | 8 | 75.0 | 25.0 | 100.0 | 0.0 |
| Putative Dm 11474157 CL | 11474156 | 0 | 0 | 0 | 1 | 0 | 0 | 0 | 5 | 6 | 16.7 | 83.3 | 0.0 | 100.0 |
| Putative Dm 10106711 MF | 10106710 | 1 | 0 | 3 | 0 | 0 | 0 | 0 | 0 | 4 | 100.0 | 0.0 | 25.0 | 75.0 |
| Putative Dm 11455002 MF | 11455001 | 0 | 0 | 0 | 3 | 0 | 0 | 0 | 0 | 3 | 100.0 | 0.0 | 0.0 | 100.0 |
| Total | | 3 | 8 | 3 | 4 | 0 | 6 | 0 | 5 | 29 | 62.1 | 37.9 | 58.6 | 41.4 |

#### R7 and R8 outgoing ML-VPN1

| name | skid | pR7a | pR7b | yR7a | yR7b | pR8a | pR8b | yR8a | yR8b | total | %R7 | %R8 | %p | %y |
|---|---|---|---|---|---|---|---|---|---|---|---|---|---|---|
| Putative ML_VPN1 11458373 CL | 11458372 | 0 | 0 | 0 | 0 | 14 | 0 | 0 | 1 | 15 | 0 | 100 | 93.3 | 6.7 |
| Putative ML_VPN1 11467871 HL | 11467870 | 0 | 0 | 0 | 0 | 0 | 7 | 0 | 0 | 7 | 0 | 100 | 100.0 | 0.0 |
| Putative ML_VPN1 11466212 CL | 11466211 | 0 | 0 | 0 | 0 | 5 | 0 | 0 | 0 | 5 | 0 | 100 | 100.0 | 0.0 |
| Total | | 0 | 0 | 0 | 0 | 19 | 7 | 0 | 1 | 27 | 0 | 100 | 96.3 | 3.7 |

#### R7 and R8 outgoing C2

| name | skid | pR7a | pR7b | yR7a | yR7b | pR8a | pR8b | yR8a | yR8b | total | %R7 | %R8 | %p | %y |
|---|---|---|---|---|---|---|---|---|---|---|---|---|---|---|
| Putative C2 11453788 CL | 11453787 | 0 | 6 | 9 | 4 | 0 | 0 | 0 | 0 | 19 | 100.0 | 0.0 | 31.6 | 68.4 |
| Putative C2 11456777 CL | 11456776 | 1 | 0 | 0 | 0 | 6 | 0 | 0 | 0 | 7 | 14.3 | 85.7 | 100.0 | 0.0 |
| Total | | 1 | 6 | 9 | 4 | 6 | 0 | 0 | 0 | 26 | 76.9 | 23.1 | 50.0 | 50.0 |

#### R7 and R8 outgoing Mt-VPN

| name | skid | pR7a | pR7b | yR7a | yR7b | pR8a | pR8b | yR8a | yR8b | total | %R7 | %R8 | %p | %y |
|---|---|---|---|---|---|---|---|---|---|---|---|---|---|---|
| Putative Mt_VPN 11453465 CL | 11453464 | 0 | 0 | 5 | 6 | 0 | 0 | 0 | 0 | 11 | 100.0 | 0.0 | 0 | 100 |
| Putative Mt_VPN 11469482 HL | 11469481 | 0 | 0 | 0 | 0 | 0 | 0 | 4 | 1 | 5 | 0.0 | 100.0 | 0 | 100 |
| Putative Mt_VPN 3401517 MR | 14286406 | 0 | 0 | 1 | 0 | 0 | 0 | 3 | 0 | 4 | 25.0 | 75.0 | 0 | 100 |
| Putative Mt_VPN LP neuron 3509521 CPM | 3509520 | 0 | 0 | 0 | 0 | 0 | 3 | 0 | 0 | 3 | 0.0 | 100.0 | 100 | 0 |
| Total | | 0 | 0 | 6 | 6 | 0 | 3 | 7 | 1 | 23 | 52.2 | 47.8 | 13 | 87 |

#### R7 and R8 outgoing Mti

| name | skid | pR7a | pR7b | yR7a | yR7b | pR8a | pR8b | yR8a | yR8b | total | %R7 | %R8 | %p | %y |
|---|---|---|---|---|---|---|---|---|---|---|---|---|---|---|
| Putative Mti 10289206 MF | 11666155 | 0 | 2 | 0 | 0 | 2 | 5 | 0 | 0 | 9 | 22.2 | 77.8 | 100.0 | 0.0 |
| Putative Mti 11481396 CL | 11481395 | 0 | 0 | 0 | 0 | 0 | 0 | 7 | 0 | 7 | 0.0 | 100.0 | 0.0 | 100.0 |
| Putative Mti 11466227 CL | 11466226 | 0 | 0 | 0 | 0 | 3 | 0 | 0 | 0 | 3 | 0.0 | 100.0 | 100.0 | 0.0 |
| Total | | 0 | 2 | 0 | 0 | 5 | 5 | 7 | 0 | 19 | 10.5 | 89.5 | 63.2 | 36.8 |

#### R7 and R8 outgoing Tm5a-like

| name | skid | pR7a | pR7b | yR7a | yR7b | pR8a | pR8b | yR8a | yR8b | total | %R7 | %R8 | %p | %y |
|---|---|---|---|---|---|---|---|---|---|---|---|---|---|---|
| Putative Tm5a-like 11481725 MF | 11481724 | 0 | 0 | 1 | 0 | 0 | 0 | 16 | 0 | 17 | 5.9 | 94.1 | 0 | 100 |
| Total | | 0 | 0 | 1 | 0 | 0 | 0 | 16 | 0 | 17 | 5.9 | 94.1 | 0 | 100 |

#### R7 and R8 outgoing TmY10

| name | skid | pR7a | pR7b | yR7a | yR7b | pR8a | pR8b | yR8a | yR8b | total | %R7 | %R8 | %p | %y |
|---|---|---|---|---|---|---|---|---|---|---|---|---|---|---|
| Putative TmY10 11449412 HL | 11449411 | 0 | 1 | 0 | 0 | 0 | 6 | 0 | 0 | 7 | 14.3 | 85.7 | 100 | 0 |
| Total | | 0 | 1 | 0 | 0 | 0 | 6 | 0 | 0 | 7 | 14.3 | 85.7 | 100 | 0 |

#### R7 and R8 outgoing Mi10

| name | skid | pR7a | pR7b | yR7a | yR7b | pR8a | pR8b | yR8a | yR8b | total | %R7 | %R8 | %p | %y |
|---|---|---|---|---|---|---|---|---|---|---|---|---|---|---|
| Putative Mi10 11481717 MF | 11481716 | 0 | 0 | 0 | 0 | 0 | 0 | 5 | 0 | 5 | 0 | 100 | 0 | 100 |
| Total | | 0 | 0 | 0 | 0 | 0 | 0 | 5 | 0 | 5 | 0 | 100 | 0 | 100 |

#### R7 and R8 outgoing Mi

| name | skid | pR7a | pR7b | yR7a | yR7b | pR8a | pR8b | yR8a | yR8b | total | %R7 | %R8 | %p | %y |
|---|---|---|---|---|---|---|---|---|---|---|---|---|---|---|
| Putative Mi 11453950 HL | 11453949 | 0 | 2 | 0 | 1 | 0 | 0 | 0 | 0 | 3 | 100 | 0 | 66.7 | 33.3 |
| Total | | 0 | 2 | 0 | 1 | 0 | 0 | 0 | 0 | 3 | 100 | 0 | 66.7 | 33.3 |

#### R7 and R8 outgoing C3

| name | skid | pR7a | pR7b | yR7a | yR7b | pR8a | pR8b | yR8a | yR8b | total | %R7 | %R8 | %p | %y |
|---|---|---|---|---|---|---|---|---|---|---|---|---|---|---|
| Putative C3 11471582 CL | 11471581 | 0 | 0 | 0 | 0 | 0 | 0 | 0 | 3 | 3 | 0 | 100 | 0 | 100 |
| Total | | 0 | 0 | 0 | 0 | 0 | 0 | 0 | 3 | 3 | 0 | 100 | 0 | 100 |

#### R7 and R8 outgoing Identified-<3

| name | skid | pR7a | pR7b | yR7a | yR7b | pR8a | pR8b | yR8a | yR8b | total | %R7 | %R8 | %p | %y |
|---|---|---|---|---|---|---|---|---|---|---|---|---|---|---|
| Putative Tm 11469206 HL | 11469205 | 0 | 0 | 0 | 0 | 0 | 2 | 0 | 0 | 2 | 0.0 | 100.0 | 100.0 | 0.0 |
| Putative Tm 11458353 MF | 11458352 | 0 | 0 | 0 | 0 | 2 | 0 | 0 | 0 | 2 | 0.0 | 100.0 | 100.0 | 0.0 |
| Putative Tm 11448401 MF | 11448400 | 0 | 0 | 2 | 0 | 0 | 0 | 0 | 0 | 2 | 100.0 | 0.0 | 0.0 | 100.0 |
| Putative Tm 10649077 MF | 10649076 | 0 | 0 | 0 | 0 | 0 | 0 | 2 | 0 | 2 | 0.0 | 100.0 | 0.0 | 100.0 |
| Putative Mt_VPN 4711709 ME | 4711708 | 0 | 0 | 0 | 0 | 0 | 2 | 0 | 0 | 2 | 0.0 | 100.0 | 100.0 | 0.0 |
| Putative ML_VPN2 11474247 CL | 11671334 | 0 | 0 | 0 | 0 | 0 | 0 | 0 | 2 | 2 | 0.0 | 100.0 | 0.0 | 100.0 |
| Putative MeTu 14838260 AT | 14838259 | 0 | 0 | 2 | 0 | 0 | 0 | 0 | 0 | 2 | 100.0 | 0.0 | 0.0 | 100.0 |
| Putative Dm2 11449695 HL | 11449694 | 0 | 2 | 0 | 0 | 0 | 0 | 0 | 0 | 2 | 100.0 | 0.0 | 100.0 | 0.0 |
| Putative Dm11 11455072 HL | 11455071 | 0 | 0 | 0 | 2 | 0 | 0 | 0 | 0 | 2 | 100.0 | 0.0 | 0.0 | 100.0 |
| Putative Dm 10562975 MF | 10562974 | 0 | 2 | 0 | 0 | 0 | 0 | 0 | 0 | 2 | 100.0 | 0.0 | 100.0 | 0.0 |
| Putative C2 11472486 HL | 11472485 | 0 | 0 | 0 | 0 | 0 | 2 | 0 | 0 | 2 | 0.0 | 100.0 | 100.0 | 0.0 |
| Putative Tm3 14359935 CL | 14359934 | 0 | 0 | 1 | 0 | 0 | 0 | 0 | 0 | 1 | 100.0 | 0.0 | 0.0 | 100.0 |
| Putative Tm1 8942830690 CL | 14653782 | 0 | 0 | 0 | 0 | 1 | 0 | 0 | 0 | 1 | 0.0 | 100.0 | 100.0 | 0.0 |
| Putative Tm 9725279989 CL | 14767205 | 0 | 0 | 0 | 1 | 0 | 0 | 0 | 0 | 1 | 100.0 | 0.0 | 0.0 | 100.0 |
| Putative Tm 15805232 HL | 15805231 | 0 | 0 | 1 | 0 | 0 | 0 | 0 | 0 | 1 | 100.0 | 0.0 | 0.0 | 100.0 |
| Putative Tm 11749586 HL | 11749585 | 0 | 0 | 1 | 0 | 0 | 0 | 0 | 0 | 1 | 100.0 | 0.0 | 0.0 | 100.0 |
| Putative Tm 10657485 HL | 10657484 | 0 | 0 | 1 | 0 | 0 | 0 | 0 | 0 | 1 | 100.0 | 0.0 | 0.0 | 100.0 |
| Putative Tm 10547529 MF | 10547528 | 0 | 0 | 1 | 0 | 0 | 0 | 0 | 0 | 1 | 100.0 | 0.0 | 0.0 | 100.0 |
| Putative T1 11454504 MF | 11454503 | 0 | 0 | 0 | 0 | 1 | 0 | 0 | 0 | 1 | 0.0 | 100.0 | 100.0 | 0.0 |
| Putative Mi14 8421422362 CL | 14746351 | 0 | 0 | 0 | 0 | 1 | 0 | 0 | 0 | 1 | 0.0 | 100.0 | 100.0 | 0.0 |
| Putative Mi1 8809115577 CL | 14880472 | 0 | 0 | 0 | 0 | 1 | 0 | 0 | 0 | 1 | 0.0 | 100.0 | 100.0 | 0.0 |
| Putative MeTu 10656251 MF | 10656250 | 0 | 0 | 0 | 1 | 0 | 0 | 0 | 0 | 1 | 100.0 | 0.0 | 0.0 | 100.0 |
| Putative LaWF1 8286417189 CL | 14624795 | 0 | 0 | 0 | 0 | 0 | 1 | 0 | 0 | 1 | 0.0 | 100.0 | 100.0 | 0.0 |
| Putative Dm 11723760 MF | 11723759 | 0 | 0 | 0 | 0 | 1 | 0 | 0 | 0 | 1 | 0.0 | 100.0 | 100.0 | 0.0 |
| Putative Dm 10638023 HL | 10638022 | 0 | 0 | 1 | 0 | 0 | 0 | 0 | 0 | 1 | 100.0 | 0.0 | 0.0 | 100.0 |
| Putative Dm 10537971 MF | 10537970 | 0 | 1 | 0 | 0 | 0 | 0 | 0 | 0 | 1 | 100.0 | 0.0 | 100.0 | 0.0 |
| Total | | 0 | 5 | 10 | 4 | 7 | 7 | 2 | 2 | 37 | 51.4 | 48.6 | 51.4 | 48.6 |

#### R7 and R8 outgoing Unidentified->=3

| name | skid | pR7a | pR7b | yR7a | yR7b | pR8a | pR8b | yR8a | yR8b | total | %R7 | %R8 | %p | %y |
|---|---|---|---|---|---|---|---|---|---|---|---|---|---|---|
| Neuron 15934960 HL | 15934959 | 0 | 0 | 0 | 0 | 0 | 5 | 0 | 0 | 5 | 0 | 100 | 100.0 | 0.0 |
| neuron 11652951 HL | 11469649 | 0 | 0 | 0 | 0 | 0 | 0 | 2 | 1 | 3 | 0 | 100 | 0.0 | 100.0 |
| Total | | 0 | 0 | 0 | 0 | 0 | 5 | 2 | 1 | 8 | 0 | 100 | 62.5 | 37.5 |

#### R7 and R8 outgoing Unidentified-<3

| name | skid | pR7a | pR7b | yR7a | yR7b | pR8a | pR8b | yR8a | yR8b | total | %R7 | %R8 | %p | %y |
|---|---|---|---|---|---|---|---|---|---|---|---|---|---|---|
| neuron 8674787766 HL | 15942569 | 0 | 0 | 0 | 0 | 0 | 0 | 0 | 2 | 2 | 0.0 | 100.0 | 0.0 | 100.0 |
| neuron 8454892 SMA | 8454891 | 0 | 0 | 0 | 0 | 0 | 0 | 0 | 2 | 2 | 0.0 | 100.0 | 0.0 | 100.0 |
| Neuron 217388042 HL | 15984291 | 0 | 2 | 0 | 0 | 0 | 0 | 0 | 0 | 2 | 100.0 | 0.0 | 100.0 | 0.0 |
| neuron 17131131 MF | 17131130 | 0 | 0 | 0 | 0 | 0 | 0 | 2 | 0 | 2 | 0.0 | 100.0 | 0.0 | 100.0 |
| neuron 11483242 HL | 11483241 | 0 | 0 | 0 | 0 | 0 | 0 | 2 | 0 | 2 | 0.0 | 100.0 | 0.0 | 100.0 |
| neuron 11470831 HL | 11470830 | 0 | 0 | 0 | 0 | 0 | 2 | 0 | 0 | 2 | 0.0 | 100.0 | 100.0 | 0.0 |
| neuron 11326962 CL | 11326961 | 0 | 0 | 1 | 0 | 0 | 0 | 1 | 0 | 2 | 50.0 | 50.0 | 0.0 | 100.0 |
| neuron 10563217 MF | 10563216 | 0 | 2 | 0 | 0 | 0 | 0 | 0 | 0 | 2 | 100.0 | 0.0 | 100.0 | 0.0 |
| neuron 17159865 | 17159864 | 0 | 0 | 0 | 0 | 1 | 0 | 0 | 0 | 1 | 0.0 | 100.0 | 100.0 | 0.0 |
| neuron 17131177 MF | 17131176 | 1 | 0 | 0 | 0 | 0 | 0 | 0 | 0 | 1 | 100.0 | 0.0 | 100.0 | 0.0 |
| Neuron 15901693 | 15901692 | 0 | 0 | 1 | 0 | 0 | 0 | 0 | 0 | 1 | 100.0 | 0.0 | 0.0 | 100.0 |
| Neuron 11598593 | 11598592 | 0 | 0 | 0 | 0 | 1 | 0 | 0 | 0 | 1 | 0.0 | 100.0 | 100.0 | 0.0 |
| neuron 11512294 HL | 11512293 | 0 | 0 | 0 | 0 | 0 | 1 | 0 | 0 | 1 | 0.0 | 100.0 | 100.0 | 0.0 |
| neuron 11511048 | 11511047 | 0 | 1 | 0 | 0 | 0 | 0 | 0 | 0 | 1 | 100.0 | 0.0 | 100.0 | 0.0 |
| neuron 11481388 | 11481387 | 0 | 0 | 0 | 0 | 0 | 0 | 1 | 0 | 1 | 0.0 | 100.0 | 0.0 | 100.0 |
| neuron 11480446 | 11480445 | 0 | 0 | 0 | 0 | 0 | 0 | 1 | 0 | 1 | 0.0 | 100.0 | 0.0 | 100.0 |
| neuron 11475567 | 11475566 | 0 | 0 | 0 | 0 | 0 | 0 | 1 | 0 | 1 | 0.0 | 100.0 | 0.0 | 100.0 |
| neuron 11474876 | 11474875 | 0 | 0 | 0 | 0 | 0 | 0 | 1 | 0 | 1 | 0.0 | 100.0 | 0.0 | 100.0 |
| neuron 11474415 | 11474414 | 0 | 0 | 0 | 0 | 0 | 0 | 0 | 1 | 1 | 0.0 | 100.0 | 0.0 | 100.0 |
| neuron 11474213 CL | 11474212 | 0 | 0 | 0 | 0 | 0 | 0 | 0 | 1 | 1 | 0.0 | 100.0 | 0.0 | 100.0 |
| neuron 11474053 | 11474052 | 0 | 0 | 0 | 0 | 0 | 0 | 1 | 0 | 1 | 0.0 | 100.0 | 0.0 | 100.0 |
| neuron 11474033 | 11474032 | 0 | 0 | 0 | 0 | 0 | 0 | 1 | 0 | 1 | 0.0 | 100.0 | 0.0 | 100.0 |
| neuron 11474028 | 11474027 | 0 | 0 | 0 | 0 | 0 | 0 | 1 | 0 | 1 | 0.0 | 100.0 | 0.0 | 100.0 |
| neuron 11473958 | 11473957 | 0 | 0 | 0 | 0 | 0 | 0 | 1 | 0 | 1 | 0.0 | 100.0 | 0.0 | 100.0 |
| neuron 11473474 CL | 11473473 | 0 | 0 | 0 | 0 | 0 | 0 | 0 | 1 | 1 | 0.0 | 100.0 | 0.0 | 100.0 |
| neuron 11472913 | 11472912 | 0 | 0 | 0 | 0 | 0 | 0 | 1 | 0 | 1 | 0.0 | 100.0 | 0.0 | 100.0 |
| neuron 11472423 | 11472422 | 0 | 0 | 0 | 0 | 0 | 0 | 0 | 1 | 1 | 0.0 | 100.0 | 0.0 | 100.0 |
| neuron 11471813 HL | 11471812 | 0 | 0 | 0 | 0 | 0 | 1 | 0 | 0 | 1 | 0.0 | 100.0 | 100.0 | 0.0 |
| neuron 11471528 | 11471527 | 0 | 0 | 0 | 0 | 0 | 1 | 0 | 0 | 1 | 0.0 | 100.0 | 100.0 | 0.0 |
| neuron 11469592 | 11469591 | 0 | 0 | 0 | 0 | 0 | 1 | 0 | 0 | 1 | 0.0 | 100.0 | 100.0 | 0.0 |
| neuron 11469561 | 11469560 | 0 | 0 | 0 | 0 | 0 | 0 | 1 | 0 | 1 | 0.0 | 100.0 | 0.0 | 100.0 |
| neuron 11468662 | 11468661 | 0 | 0 | 0 | 0 | 0 | 0 | 0 | 1 | 1 | 0.0 | 100.0 | 0.0 | 100.0 |
| neuron 11467899 | 11467898 | 0 | 0 | 0 | 0 | 0 | 1 | 0 | 0 | 1 | 0.0 | 100.0 | 100.0 | 0.0 |
| neuron 11467356 | 11467355 | 0 | 0 | 0 | 0 | 0 | 1 | 0 | 0 | 1 | 0.0 | 100.0 | 100.0 | 0.0 |
| neuron 11459213 | 11459212 | 0 | 0 | 0 | 0 | 1 | 0 | 0 | 0 | 1 | 0.0 | 100.0 | 100.0 | 0.0 |
| neuron 11459079 | 11459078 | 0 | 0 | 0 | 0 | 1 | 0 | 0 | 0 | 1 | 0.0 | 100.0 | 100.0 | 0.0 |
| neuron 11458409 | 11458408 | 0 | 0 | 0 | 0 | 1 | 0 | 0 | 0 | 1 | 0.0 | 100.0 | 100.0 | 0.0 |
| neuron 11458393 CL | 11458392 | 0 | 0 | 0 | 0 | 1 | 0 | 0 | 0 | 1 | 0.0 | 100.0 | 100.0 | 0.0 |
| neuron 11458262 | 11458261 | 0 | 0 | 0 | 0 | 1 | 0 | 0 | 0 | 1 | 0.0 | 100.0 | 100.0 | 0.0 |
| neuron 11458257 | 11458256 | 0 | 0 | 0 | 0 | 1 | 0 | 0 | 0 | 1 | 0.0 | 100.0 | 100.0 | 0.0 |
| neuron 11458247 | 11458246 | 0 | 0 | 0 | 0 | 1 | 0 | 0 | 0 | 1 | 0.0 | 100.0 | 100.0 | 0.0 |
| neuron 11458187 | 11458186 | 0 | 0 | 0 | 0 | 1 | 0 | 0 | 0 | 1 | 0.0 | 100.0 | 100.0 | 0.0 |
| neuron 11458100 | 11458099 | 0 | 0 | 0 | 0 | 1 | 0 | 0 | 0 | 1 | 0.0 | 100.0 | 100.0 | 0.0 |
| neuron 11458095 | 11458094 | 0 | 0 | 0 | 0 | 1 | 0 | 0 | 0 | 1 | 0.0 | 100.0 | 100.0 | 0.0 |
| neuron 11457627 MF | 11457626 | 0 | 0 | 0 | 0 | 1 | 0 | 0 | 0 | 1 | 0.0 | 100.0 | 100.0 | 0.0 |
| neuron 11457032 MF | 11457031 | 0 | 0 | 0 | 0 | 1 | 0 | 0 | 0 | 1 | 0.0 | 100.0 | 100.0 | 0.0 |
| neuron 11455152 | 11455151 | 0 | 0 | 0 | 1 | 0 | 0 | 0 | 0 | 1 | 100.0 | 0.0 | 0.0 | 100.0 |
| neuron 11454942 | 11454941 | 0 | 0 | 0 | 1 | 0 | 0 | 0 | 0 | 1 | 100.0 | 0.0 | 0.0 | 100.0 |
| neuron 11454742 | 11454741 | 0 | 0 | 0 | 1 | 0 | 0 | 0 | 0 | 1 | 100.0 | 0.0 | 0.0 | 100.0 |
| neuron 11453858 | 11453857 | 0 | 0 | 0 | 1 | 0 | 0 | 0 | 0 | 1 | 100.0 | 0.0 | 0.0 | 100.0 |
| neuron 11453555 | 11453554 | 0 | 0 | 0 | 1 | 0 | 0 | 0 | 0 | 1 | 100.0 | 0.0 | 0.0 | 100.0 |
| neuron 11453195 | 11453194 | 0 | 0 | 0 | 1 | 0 | 0 | 0 | 0 | 1 | 100.0 | 0.0 | 0.0 | 100.0 |
| neuron 11452412 | 11452411 | 0 | 0 | 1 | 0 | 0 | 0 | 0 | 0 | 1 | 100.0 | 0.0 | 0.0 | 100.0 |
| neuron 11449753 | 11449752 | 0 | 1 | 0 | 0 | 0 | 0 | 0 | 0 | 1 | 100.0 | 0.0 | 100.0 | 0.0 |
| neuron 11448908 | 11448907 | 0 | 1 | 0 | 0 | 0 | 0 | 0 | 0 | 1 | 100.0 | 0.0 | 100.0 | 0.0 |
| neuron 11448383 | 11448382 | 0 | 0 | 1 | 0 | 0 | 0 | 0 | 0 | 1 | 100.0 | 0.0 | 0.0 | 100.0 |
| neuron 11448211 | 11448210 | 0 | 0 | 1 | 0 | 0 | 0 | 0 | 0 | 1 | 100.0 | 0.0 | 0.0 | 100.0 |
| neuron 11448038 | 11448037 | 0 | 0 | 1 | 0 | 0 | 0 | 0 | 0 | 1 | 100.0 | 0.0 | 0.0 | 100.0 |
| neuron 11447605 | 11447604 | 0 | 0 | 1 | 0 | 0 | 0 | 0 | 0 | 1 | 100.0 | 0.0 | 0.0 | 100.0 |
| neuron 11447258 HL | 11447257 | 0 | 0 | 1 | 0 | 0 | 0 | 0 | 0 | 1 | 100.0 | 0.0 | 0.0 | 100.0 |
| neuron 11446424 HL | 11446423 | 1 | 0 | 0 | 0 | 0 | 0 | 0 | 0 | 1 | 100.0 | 0.0 | 100.0 | 0.0 |
| Google: 9334377939 HL | 15901648 | 0 | 0 | 1 | 0 | 0 | 0 | 0 | 0 | 1 | 100.0 | 0.0 | 0.0 | 100.0 |
| Total | | 2 | 7 | 9 | 6 | 14 | 8 | 15 | 9 | 70 | 34.3 | 65.7 | 44.3 | 55.7 |

#### R7 and R8 incoming Dm9

| name | skid | pR7a | pR7b | yR7a | yR7b | pR8a | pR8b | yR8a | yR8b | total | %R7 | %R8 | %p | %y |
|---|---|---|---|---|---|---|---|---|---|---|---|---|---|---|
| Putative Dm9 11452428 MF | 11452427 | 45 | 35 | 39 | 35 | 33 | 36 | 29 | 29 | 281 | 54.8 | 45.2 | 53.0 | 47.0 |
| Putative Dm9 11447062 CL | 11447061 | 2 | 0 | 9 | 0 | 0 | 0 | 7 | 0 | 18 | 61.1 | 38.9 | 11.1 | 88.9 |
| Putative Dm9 11450496 CL | 11450495 | 0 | 0 | 3 | 4 | 0 | 0 | 2 | 3 | 12 | 58.3 | 41.7 | 0.0 | 100.0 |
| Putative Dm9 11484680 MF | 11484679 | 4 | 0 | 0 | 0 | 5 | 0 | 0 | 0 | 9 | 44.4 | 55.6 | 100.0 | 0.0 |
| Putative Dm9 11454715 CL | 11454714 | 0 | 0 | 0 | 2 | 0 | 3 | 0 | 2 | 7 | 28.6 | 71.4 | 42.9 | 57.1 |
| Putative Dm9 11444387 HL | 11444386 | 0 | 4 | 0 | 0 | 0 | 2 | 0 | 0 | 6 | 66.7 | 33.3 | 100.0 | 0.0 |
| Total | | 51 | 39 | 51 | 41 | 38 | 41 | 38 | 34 | 333 | 54.7 | 45.3 | 50.8 | 49.2 |

#### R7 and R8 incoming R8

| name | skid | pR7a | pR7b | yR7a | yR7b | pR8a | pR8b | yR8a | yR8b | total | %R7 | %R8 | %p | %y |
|---|---|---|---|---|---|---|---|---|---|---|---|---|---|---|
| Putative R8 col G1 10629255 MF | 10629254 | 0 | 0 | 46 | 0 | 0 | 0 | 0 | 0 | 46 | 100.0 | 0.0 | 0.0 | 100.0 |
| Putative R8 col A1 10086692 MF | 10086691 | 39 | 0 | 0 | 0 | 1 | 0 | 0 | 0 | 40 | 97.5 | 2.5 | 100.0 | 0.0 |
| Putative R8 col I1 11466409 HL | 11466408 | 0 | 36 | 0 | 0 | 0 | 0 | 0 | 0 | 36 | 100.0 | 0.0 | 100.0 | 0.0 |
| Putative R8 col H1 11468319 CL | 11468318 | 0 | 0 | 0 | 36 | 0 | 0 | 0 | 0 | 36 | 100.0 | 0.0 | 0.0 | 100.0 |
| Total | | 39 | 36 | 46 | 36 | 1 | 0 | 0 | 0 | 158 | 99.4 | 0.6 | 48.1 | 51.9 |

#### R7 and R8 incoming R7

| name | skid | pR7a | pR7b | yR7a | yR7b | pR8a | pR8b | yR8a | yR8b | total | %R7 | %R8 | %p | %y |
|---|---|---|---|---|---|---|---|---|---|---|---|---|---|---|
| Putative R7 col A1 10082583 MF | 10082582 | 0 | 0 | 0 | 0 | 20 | 0 | 0 | 0 | 20 | 0 | 100 | 100.0 | 0.0 |
| Putative R7 col H1 10653594 MF | 10653593 | 0 | 0 | 0 | 0 | 0 | 0 | 0 | 19 | 19 | 0 | 100 | 0.0 | 100.0 |
| Putative R7 col I1 10538511 MF | 10538510 | 0 | 0 | 0 | 0 | 0 | 11 | 0 | 0 | 11 | 0 | 100 | 100.0 | 0.0 |
| Putative R7 col G1 10585941 MF | 10585940 | 0 | 0 | 0 | 0 | 0 | 0 | 7 | 0 | 7 | 0 | 100 | 0.0 | 100.0 |
| Total | | 0 | 0 | 0 | 0 | 20 | 11 | 7 | 19 | 57 | 0 | 100 | 54.4 | 45.6 |

#### R7 and R8 incoming Mt-VPN

| name | skid | pR7a | pR7b | yR7a | yR7b | pR8a | pR8b | yR8a | yR8b | total | %R7 | %R8 | %p | %y |
|---|---|---|---|---|---|---|---|---|---|---|---|---|---|---|
| Putative Mt_VPN 3401517 MR | 14286406 | 0 | 0 | 2 | 0 | 0 | 0 | 3 | 0 | 5 | 40 | 60 | 0 | 100 |
| Total | | 0 | 0 | 2 | 0 | 0 | 0 | 3 | 0 | 5 | 40 | 60 | 0 | 100 |

#### R7 and R8 incoming C2

| name | skid | pR7a | pR7b | yR7a | yR7b | pR8a | pR8b | yR8a | yR8b | total | %R7 | %R8 | %p | %y |
|---|---|---|---|---|---|---|---|---|---|---|---|---|---|---|
| Putative C2 11453788 CL | 11453787 | 0 | 2 | 1 | 2 | 0 | 0 | 0 | 0 | 5 | 100 | 0 | 40 | 60 |
| Total | | 0 | 2 | 1 | 2 | 0 | 0 | 0 | 0 | 5 | 100 | 0 | 40 | 60 |

#### R7 and R8 incoming L3

| name | skid | pR7a | pR7b | yR7a | yR7b | pR8a | pR8b | yR8a | yR8b | total | %R7 | %R8 | %p | %y |
|---|---|---|---|---|---|---|---|---|---|---|---|---|---|---|
| Putative L3 col A1 11445252 MF | 11445251 | 0 | 0 | 0 | 0 | 4 | 0 | 0 | 0 | 4 | 0 | 100 | 100 | 0 |
| Total | | 0 | 0 | 0 | 0 | 4 | 0 | 0 | 0 | 4 | 0 | 100 | 100 | 0 |

#### R7 and R8 incoming Identified-<3

| name | skid | pR7a | pR7b | yR7a | yR7b | pR8a | pR8b | yR8a | yR8b | total | %R7 | %R8 | %p | %y |
|---|---|---|---|---|---|---|---|---|---|---|---|---|---|---|
| Putative Dm8 10109587 MF | 10109586 | 2 | 0 | 0 | 0 | 0 | 0 | 0 | 0 | 2 | 100.0 | 0.0 | 100.0 | 0.0 |
| Putative R7 10653781 MF | 10653780 | 0 | 0 | 0 | 2 | 0 | 0 | 0 | 0 | 2 | 100.0 | 0.0 | 0.0 | 100.0 |
| Putative L3 col I1 11448597 HL | 11448596 | 0 | 0 | 0 | 0 | 0 | 2 | 0 | 0 | 2 | 0.0 | 100.0 | 100.0 | 0.0 |
| Putative C2 11472486 HL | 11472485 | 0 | 0 | 0 | 0 | 0 | 2 | 0 | 0 | 2 | 0.0 | 100.0 | 100.0 | 0.0 |
| Putative Mi15 11453819 CL | 11453818 | 0 | 0 | 0 | 0 | 0 | 0 | 2 | 0 | 2 | 0.0 | 100.0 | 0.0 | 100.0 |
| Putative L3 col H1 11450470 CL | 11450469 | 0 | 0 | 0 | 0 | 0 | 0 | 0 | 2 | 2 | 0.0 | 100.0 | 0.0 | 100.0 |
| Putative Mi15 11450568 CL | 11450567 | 0 | 0 | 0 | 0 | 0 | 0 | 0 | 2 | 2 | 0.0 | 100.0 | 0.0 | 100.0 |
| Putative C2 11456777 CL | 11456776 | 1 | 0 | 0 | 0 | 0 | 0 | 0 | 0 | 1 | 100.0 | 0.0 | 100.0 | 0.0 |
| Putative Mi15 11484381 HL | 11484380 | 0 | 1 | 0 | 0 | 0 | 0 | 0 | 0 | 1 | 100.0 | 0.0 | 100.0 | 0.0 |
| Putative Tm5b-like 11447921 MF | 11447920 | 0 | 0 | 0 | 0 | 0 | 0 | 1 | 0 | 1 | 0.0 | 100.0 | 0.0 | 100.0 |
| Putative Dm11 11450454 CL | 11450453 | 0 | 0 | 0 | 0 | 0 | 0 | 0 | 1 | 1 | 0.0 | 100.0 | 0.0 | 100.0 |
| Total | | 3 | 1 | 0 | 2 | 0 | 4 | 3 | 5 | 18 | 33.3 | 66.7 | 44.4 | 55.6 |

#### R7 and R8 incoming Unidentified-<3

| name | skid | pR7a | pR7b | yR7a | yR7b | pR8a | pR8b | yR8a | yR8b | total | %R7 | %R8 | %p | %y |
|---|---|---|---|---|---|---|---|---|---|---|---|---|---|---|
| neuron 11456493 | 11456492 | 0 | 1 | 0 | 0 | 0 | 0 | 0 | 0 | 1 | 100 | 0 | 100 | 0 |
| neuron 11679982 | 11679981 | 0 | 0 | 1 | 0 | 0 | 0 | 0 | 0 | 1 | 100 | 0 | 0 | 100 |
| neuron 11472423 | 11472422 | 0 | 0 | 0 | 1 | 0 | 0 | 0 | 0 | 1 | 100 | 0 | 0 | 100 |
| neuron 11466245 | 11466244 | 0 | 0 | 0 | 0 | 1 | 0 | 0 | 0 | 1 | 0 | 100 | 100 | 0 |
| Total | | 0 | 1 | 1 | 1 | 1 | 0 | 0 | 0 | 4 | 75 | 25 | 50 | 50 |

#### R7-DRA and R8-DRA outgoing Dm-DRA1

| name | skid | R7-DRA-a | R7-DRA-b | R7-DRA-c | R8-DRA-a | R8-DRA-b | R8-DRA-c | total | %R7-DRA | %R8-DRA |
|---|---|---|---|---|---|---|---|---|---|---|
| Putative Dm-DRA1 10440161 TO | 10440160 | 0 | 0 | 21 | 0 | 0 | 0 | 21 | 100 | 0 |
| Putative Dm-DRA1 11896102 EK | 11896101 | 0 | 18 | 0 | 0 | 0 | 0 | 18 | 100 | 0 |
| Putative Dm-DRA1 16766813 EK | 16766812 | 0 | 0 | 16 | 0 | 0 | 0 | 16 | 100 | 0 |
| Putative Dm-DRA1 12106450 TO | 12106449 | 16 | 0 | 0 | 0 | 0 | 0 | 16 | 100 | 0 |
| Putative Dm-DRA1 10247371 TO | 10247370 | 5 | 10 | 0 | 0 | 0 | 0 | 15 | 100 | 0 |
| Putative Dm-DRA1 17156428 EK | 17156427 | 0 | 0 | 14 | 0 | 0 | 0 | 14 | 100 | 0 |
| Putative Dm-DRA1 17155322 EK | 17155321 | 0 | 0 | 13 | 0 | 0 | 0 | 13 | 100 | 0 |
| Putative Dm-DRA1 14065299 GS-EK | 14065298 | 4 | 9 | 0 | 0 | 0 | 0 | 13 | 100 | 0 |
| Putative Dm-DRA1 10265807 TO EK | 10265806 | 8 | 5 | 0 | 0 | 0 | 0 | 13 | 100 | 0 |
| Putative Dm-DRA1 11903983 GS | 11903982 | 0 | 10 | 0 | 0 | 0 | 0 | 10 | 100 | 0 |
| Putative Dm-DRA1 11992844 EK | 11992843 | 9 | 0 | 0 | 0 | 0 | 0 | 9 | 100 | 0 |
| Putative Dm-DRA1 11141230 GS | 11141229 | 0 | 9 | 0 | 0 | 0 | 0 | 9 | 100 | 0 |
| Putative Dm-DRA1 13979722 EK | 13979721 | 6 | 2 | 0 | 0 | 0 | 0 | 8 | 100 | 0 |
| Putative Dm-DRA1 11993799 EK | 11993798 | 7 | 0 | 0 | 0 | 0 | 0 | 7 | 100 | 0 |
| Putative Dm-DRA1 11993077 EK | 11993076 | 7 | 0 | 0 | 0 | 0 | 0 | 7 | 100 | 0 |
| Putative Dm-DRA1 13607721 EK | 13607720 | 6 | 0 | 0 | 0 | 0 | 0 | 6 | 100 | 0 |
| Putative Dm-DRA1 11769715 TO EK | 11769714 | 6 | 0 | 0 | 0 | 0 | 0 | 6 | 100 | 0 |
| Putative Dm-DRA1 17170764 EK | 17170763 | 0 | 0 | 5 | 0 | 0 | 0 | 5 | 100 | 0 |
| Putative Dm-DRA1 15976818 EK | 15976817 | 0 | 3 | 0 | 0 | 0 | 0 | 3 | 100 | 0 |
| Putative Dm-DRA1 11993696 EK | 11993695 | 3 | 0 | 0 | 0 | 0 | 0 | 3 | 100 | 0 |
| Total | | 77 | 66 | 69 | 0 | 0 | 0 | 212 | 100 | 0 |

#### R7-DRA and R8-DRA outgoing Dm9

| name | skid | R7-DRA-a | R7-DRA-b | R7-DRA-c | R8-DRA-a | R8-DRA-b | R8-DRA-c | total | %R7-DRA | %R8-DRA |
|---|---|---|---|---|---|---|---|---|---|---|
| Putative Dm9 12013130 EK | 12013129 | 0 | 0 | 31 | 0 | 0 | 36 | 67 | 46.3 | 53.7 |
| Putative Dm9 11916196 EK | 11916195 | 26 | 0 | 0 | 36 | 0 | 0 | 62 | 41.9 | 58.1 |
| Putative Dm9 10657501 GS | 10657500 | 0 | 17 | 0 | 0 | 35 | 0 | 52 | 32.7 | 67.3 |
| Putative Dm9 14310275 GS | 14310274 | 0 | 16 | 0 | 0 | 5 | 0 | 21 | 76.2 | 23.8 |
| Putative Dm9 14933803 EK | 14933802 | 0 | 3 | 0 | 0 | 1 | 0 | 4 | 75.0 | 25.0 |
| Putative Dm9 10655889 EK | 10655888 | 0 | 2 | 0 | 0 | 2 | 0 | 4 | 50.0 | 50.0 |
| Total | | 26 | 38 | 31 | 36 | 43 | 36 | 210 | 45.2 | 54.8 |

#### R7-DRA and R8-DRA outgoing MeTu-DRA

| name | skid | R7-DRA-a | R7-DRA-b | R7-DRA-c | R8-DRA-a | R8-DRA-b | R8-DRA-c | total | %R7-DRA | %R8-DRA |
|---|---|---|---|---|---|---|---|---|---|---|
| Putative MeTu_DRA 11995074 EK | 11995073 | 14 | 0 | 0 | 0 | 0 | 0 | 14 | 100 | 0 |
| Putative MeTu_DRA 11994449 EK | 11994448 | 12 | 0 | 0 | 0 | 0 | 0 | 12 | 100 | 0 |
| Putative MeTu_DRA 11908711 GS | 11908710 | 0 | 10 | 0 | 0 | 0 | 0 | 10 | 100 | 0 |
| Putative MeTu_DRA 11695294 PA | 11695293 | 0 | 0 | 10 | 0 | 0 | 0 | 10 | 100 | 0 |
| Putative MeTu_DRA 10737979 TO | 15749047 | 10 | 0 | 0 | 0 | 0 | 0 | 10 | 100 | 0 |
| Putative MeTu_DRA 12106384 EK | 12106383 | 0 | 0 | 9 | 0 | 0 | 0 | 9 | 100 | 0 |
| Putative MeTu_DRA 11994564 EK | 11994563 | 8 | 0 | 0 | 0 | 0 | 0 | 8 | 100 | 0 |
| Putative MeTu_DRA 10438340 TO | 10438339 | 0 | 0 | 8 | 0 | 0 | 0 | 8 | 100 | 0 |
| Putative MeTu_DRA 7749081264 EK | 15952007 | 0 | 0 | 7 | 0 | 0 | 0 | 7 | 100 | 0 |
| Putative MeTu_DRA 13136157 EK | 13136156 | 0 | 7 | 0 | 0 | 0 | 0 | 7 | 100 | 0 |
| Putative MeTu_DRA 11995103 EK | 11995102 | 7 | 0 | 0 | 0 | 0 | 0 | 7 | 100 | 0 |
| Putative MeTu_DRA 15616478 EK | 15616477 | 0 | 6 | 0 | 0 | 0 | 0 | 6 | 100 | 0 |
| Putative MeTu_DRA 15600899 EK | 15600898 | 0 | 6 | 0 | 0 | 0 | 0 | 6 | 100 | 0 |
| Putative MeTu_DRA 11993031 EK | 11993030 | 6 | 0 | 0 | 0 | 0 | 0 | 6 | 100 | 0 |
| Putative MeTu_DRA 10479162 TO | 15698953 | 0 | 6 | 0 | 0 | 0 | 0 | 6 | 100 | 0 |
| Putative MeTu_DRA 12127370 TO | 12127369 | 5 | 0 | 0 | 0 | 0 | 0 | 5 | 100 | 0 |
| Putative MeTu_DRA 11993383 EK | 11993382 | 5 | 0 | 0 | 0 | 0 | 0 | 5 | 100 | 0 |
| Putative MeTu_DRA 11992933 EK | 11992932 | 5 | 0 | 0 | 0 | 0 | 0 | 5 | 100 | 0 |
| Putative MeTu_DRA 6844453075 EK | 15949451 | 0 | 0 | 4 | 0 | 0 | 0 | 4 | 100 | 0 |
| Putative MeTu_DRA 17250637 EK | 17250636 | 0 | 0 | 4 | 0 | 0 | 0 | 4 | 100 | 0 |
| Putative MeTu_DRA 14889232 EK | 14889231 | 0 | 0 | 4 | 0 | 0 | 0 | 4 | 100 | 0 |
| Putative MeTu_DRA 11995359 EK | 11995358 | 4 | 0 | 0 | 0 | 0 | 0 | 4 | 100 | 0 |
| Putative MeTu_DRA 11993353 EK | 11993352 | 4 | 0 | 0 | 0 | 0 | 0 | 4 | 100 | 0 |
| Putative MeTu_DRA 11993315 EK | 11993314 | 2 | 2 | 0 | 0 | 0 | 0 | 4 | 100 | 0 |
| Putative MeTu_DRA 11908744 GS | 11908743 | 0 | 4 | 0 | 0 | 0 | 0 | 4 | 100 | 0 |
| Putative MeTu_DRA 9706730253 EK | 14864504 | 0 | 3 | 0 | 0 | 0 | 0 | 3 | 100 | 0 |
| Putative MeTu_DRA 14936635 EK | 14936634 | 0 | 0 | 3 | 0 | 0 | 0 | 3 | 100 | 0 |
| Putative MeTu_DRA 11993065 EK | 11993064 | 3 | 0 | 0 | 0 | 0 | 0 | 3 | 100 | 0 |
| Putative MeTu_DRA 11908669 GS | 11908668 | 0 | 3 | 0 | 0 | 0 | 0 | 3 | 100 | 0 |
| Putative MeTu_DRA 11903993 EK | 11903992 | 0 | 3 | 0 | 0 | 0 | 0 | 3 | 100 | 0 |
| Total | | 85 | 50 | 49 | 0 | 0 | 0 | 184 | 100 | 0 |

#### R7-DRA and R8-DRA outgoing R7-DRA

| name | skid | R7-DRA-a | R7-DRA-b | R7-DRA-c | R8-DRA-a | R8-DRA-b | R8-DRA-c | total | %R7-DRA | %R8-DRA |
|---|---|---|---|---|---|---|---|---|---|---|
| Putative R7_DRA 11728780 TO | 11728779 | 0 | 0 | 1 | 0 | 0 | 37 | 38 | 2.6 | 97.4 |
| Putative R7_DRA 10300950 TO | 10300949 | 0 | 0 | 0 | 0 | 38 | 0 | 38 | 0.0 | 100.0 |
| Putative R7_DRA 10191736 TO | 10191735 | 0 | 0 | 0 | 33 | 0 | 0 | 33 | 0.0 | 100.0 |
| Total | | 0 | 0 | 1 | 33 | 38 | 37 | 109 | 0.9 | 99.1 |

#### R7-DRA and R8-DRA outgoing Dm-DRA2

| name | skid | R7-DRA-a | R7-DRA-b | R7-DRA-c | R8-DRA-a | R8-DRA-b | R8-DRA-c | total | %R7-DRA | %R8-DRA |
|---|---|---|---|---|---|---|---|---|---|---|
| Putative Dm-DRA2 10411789 TO | 10411788 | 0 | 1 | 0 | 0 | 31 | 0 | 32 | 3.1 | 96.9 |
| Putative Dm-DRA2 10483849 TO | 10483848 | 0 | 0 | 0 | 19 | 0 | 0 | 19 | 0.0 | 100.0 |
| Putative Dm-DRA2 11710980 TO | 11710979 | 0 | 0 | 1 | 0 | 0 | 12 | 13 | 7.7 | 92.3 |
| Putative Dm-DRA2 10449077 TO | 10449076 | 0 | 0 | 0 | 9 | 0 | 0 | 9 | 0.0 | 100.0 |
| Putative Dm-DRA2 10484772 TO | 10484771 | 0 | 0 | 0 | 0 | 8 | 0 | 8 | 0.0 | 100.0 |
| Putative Dm-DRA2 17163677 EK | 17163676 | 0 | 0 | 0 | 0 | 0 | 7 | 7 | 0.0 | 100.0 |
| Putative Dm-DRA2 13262260 EK | 13262259 | 0 | 0 | 0 | 7 | 0 | 0 | 7 | 0.0 | 100.0 |
| Putative Dm-DRA2 14311484 GS | 14311483 | 0 | 1 | 0 | 0 | 5 | 0 | 6 | 16.7 | 83.3 |
| Putative Dm-DRA2 11918437 EK | 11918436 | 0 | 0 | 0 | 5 | 0 | 0 | 5 | 0.0 | 100.0 |
| Total | | 0 | 2 | 1 | 40 | 44 | 19 | 106 | 2.8 | 97.2 |

#### R7-DRA and R8-DRA outgoing Dm2

| name | skid | R7-DRA-a | R7-DRA-b | R7-DRA-c | R8-DRA-a | R8-DRA-b | R8-DRA-c | total | %R7-DRA | %R8-DRA |
|---|---|---|---|---|---|---|---|---|---|---|
| Putative Dm2 17162383 EK | 17162382 | 0 | 0 | 6 | 0 | 0 | 15 | 21 | 28.6 | 71.4 |
| Putative Dm2 11918131 EK | 11918130 | 5 | 0 | 0 | 7 | 0 | 0 | 12 | 41.7 | 58.3 |
| Putative Dm2 14065354 EK | 14065353 | 0 | 10 | 0 | 0 | 1 | 0 | 11 | 90.9 | 9.1 |
| Putative Dm2 10655669 TO | 10655668 | 0 | 0 | 0 | 0 | 4 | 0 | 4 | 0.0 | 100.0 |
| Total | | 5 | 10 | 6 | 7 | 5 | 15 | 48 | 43.8 | 56.2 |

#### R7-DRA and R8-DRA outgoing R8-DRA

| name | skid | R7-DRA-a | R7-DRA-b | R7-DRA-c | R8-DRA-a | R8-DRA-b | R8-DRA-c | total | %R7-DRA | %R8-DRA |
|---|---|---|---|---|---|---|---|---|---|---|
| Putative R8_DRA 10300964 TO | 10300963 | 0 | 16 | 0 | 0 | 0 | 0 | 16 | 100.0 | 0.0 |
| Putative R8_DRA 11728828 TO | 11728827 | 0 | 0 | 15 | 0 | 0 | 0 | 15 | 100.0 | 0.0 |
| Putative R8_DRA 10190509 TO | 10190508 | 10 | 0 | 0 | 1 | 0 | 0 | 11 | 90.9 | 9.1 |
| Total | | 10 | 16 | 15 | 1 | 0 | 0 | 42 | 97.6 | 2.4 |

#### R7-DRA and R8-DRA outgoing Mi15

| name | skid | R7-DRA-a | R7-DRA-b | R7-DRA-c | R8-DRA-a | R8-DRA-b | R8-DRA-c | total | %R7-DRA | %R8-DRA |
|---|---|---|---|---|---|---|---|---|---|---|
| Putative Mi15 10655927 TO | 10655926 | 0 | 8 | 0 | 0 | 10 | 0 | 18 | 44.4 | 55.6 |
| Putative Mi15 12017151 EK | 12017150 | 0 | 0 | 5 | 0 | 0 | 5 | 10 | 50.0 | 50.0 |
| Putative Mi15 11916191 EK | 11916190 | 4 | 0 | 0 | 3 | 0 | 0 | 7 | 57.1 | 42.9 |
| Putative Mi15 12002140 EK | 12002139 | 0 | 0 | 4 | 0 | 0 | 0 | 4 | 100.0 | 0.0 |
| Total | | 4 | 8 | 9 | 3 | 10 | 5 | 39 | 53.8 | 46.2 |

#### R7-DRA and R8-DRA outgoing Mti-DRA-1

| name | skid | R7-DRA-a | R7-DRA-b | R7-DRA-c | R8-DRA-a | R8-DRA-b | R8-DRA-c | total | %R7-DRA | %R8-DRA |
|---|---|---|---|---|---|---|---|---|---|---|
| Putative Mti_DRA_1 11993472 EK | 11993471 | 6 | 7 | 4 | 0 | 0 | 0 | 17 | 100 | 0 |
| Putative Mti_DRA_1 14430737 EK | 14430736 | 4 | 2 | 1 | 0 | 0 | 0 | 7 | 100 | 0 |
| Putative Mti_DRA_1 11994581 EK | 11994580 | 2 | 0 | 2 | 0 | 0 | 0 | 4 | 100 | 0 |
| Putative Mti_DRA_1 11995267 EK | 11995266 | 1 | 0 | 2 | 0 | 0 | 0 | 3 | 100 | 0 |
| Putative Mti_DRA_1 11686948 EK | 11686947 | 0 | 1 | 2 | 0 | 0 | 0 | 3 | 100 | 0 |
| Putative Mti_DRA_1 10474754 PA | 10474753 | 0 | 0 | 3 | 0 | 0 | 0 | 3 | 100 | 0 |
| Total | | 13 | 10 | 14 | 0 | 0 | 0 | 37 | 100 | 0 |

#### R7-DRA and R8-DRA outgoing MeMe-DRA

| name | skid | R7-DRA-a | R7-DRA-b | R7-DRA-c | R8-DRA-a | R8-DRA-b | R8-DRA-c | total | %R7-DRA | %R8-DRA |
|---|---|---|---|---|---|---|---|---|---|---|
| Putative MeMe_DRA 10439443 EK | 10439442 | 6 | 0 | 21 | 0 | 0 | 0 | 27 | 100 | 0 |
| Putative MeMe_DRA 11993544 EK | 11993543 | 9 | 0 | 0 | 0 | 0 | 0 | 9 | 100 | 0 |
| Total | | 15 | 0 | 21 | 0 | 0 | 0 | 36 | 100 | 0 |

#### R7-DRA and R8-DRA outgoing L3

| name | skid | R7-DRA-a | R7-DRA-b | R7-DRA-c | R8-DRA-a | R8-DRA-b | R8-DRA-c | total | %R7-DRA | %R8-DRA |
|---|---|---|---|---|---|---|---|---|---|---|
| Putative L3 12018070 EK | 12018069 | 0 | 0 | 7 | 0 | 0 | 6 | 13 | 53.8 | 46.2 |
| Putative L3 10653986 EK | 10653985 | 0 | 5 | 0 | 0 | 7 | 0 | 12 | 41.7 | 58.3 |
| Putative L3 11917228 EK | 11917227 | 4 | 0 | 0 | 3 | 0 | 0 | 7 | 57.1 | 42.9 |
| Total | | 4 | 5 | 7 | 3 | 7 | 6 | 32 | 50.0 | 50.0 |

#### R7-DRA and R8-DRA outgoing VPN-DRA

| name | skid | R7-DRA-a | R7-DRA-b | R7-DRA-c | R8-DRA-a | R8-DRA-b | R8-DRA-c | total | %R7-DRA | %R8-DRA |
|---|---|---|---|---|---|---|---|---|---|---|
| Putative VPN_DRA ME.R 11992908 EK | 11992907 | 2 | 0 | 6 | 0 | 0 | 0 | 8 | 100 | 0 |
| Putative VPN_DRA ME.R 11904074 GS EK | 11904073 | 5 | 2 | 0 | 0 | 0 | 0 | 7 | 100 | 0 |
| Putative VPN_DRA ME.R 4271777 EK | 4271776 | 1 | 4 | 0 | 0 | 0 | 0 | 5 | 100 | 0 |
| Putative VPN_DRA ME.R 17165716 EK | 17165715 | 0 | 0 | 4 | 0 | 0 | 0 | 4 | 100 | 0 |
| Putative VPN_DRA ME.R 8541155621 EK | 15969128 | 3 | 0 | 0 | 0 | 0 | 0 | 3 | 100 | 0 |
| Putative VPN_DRA ME.R 16886431 PA | 16886430 | 0 | 0 | 3 | 0 | 0 | 0 | 3 | 100 | 0 |
| Total | | 11 | 6 | 13 | 0 | 0 | 0 | 30 | 100 | 0 |

#### R7-DRA and R8-DRA outgoing L1

| name | skid | R7-DRA-a | R7-DRA-b | R7-DRA-c | R8-DRA-a | R8-DRA-b | R8-DRA-c | total | %R7-DRA | %R8-DRA |
|---|---|---|---|---|---|---|---|---|---|---|
| Putative L1 10654563 EK-GS | 10654562 | 0 | 3 | 0 | 0 | 5 | 0 | 8 | 37.5 | 62.5 |
| Putative L1 11915726 EK | 11915725 | 3 | 0 | 0 | 4 | 0 | 0 | 7 | 42.9 | 57.1 |
| Putative L1 12019445 EK | 12019444 | 0 | 0 | 3 | 0 | 0 | 0 | 3 | 100.0 | 0.0 |
| Total | | 3 | 3 | 3 | 4 | 5 | 0 | 18 | 50.0 | 50.0 |

#### R7-DRA and R8-DRA outgoing Tm20

| name | skid | R7-DRA-a | R7-DRA-b | R7-DRA-c | R8-DRA-a | R8-DRA-b | R8-DRA-c | total | %R7-DRA | %R8-DRA |
|---|---|---|---|---|---|---|---|---|---|---|
| Putative Tm20 12018770 EK | 12018769 | 0 | 0 | 0 | 0 | 0 | 8 | 8 | 0 | 100 |
| Putative Tm20 11918620 EK | 11918619 | 0 | 0 | 0 | 4 | 0 | 0 | 4 | 0 | 100 |
| Putative Tm20 10655482 EK | 10655481 | 0 | 0 | 0 | 0 | 4 | 0 | 4 | 0 | 100 |
| Total | | 0 | 0 | 0 | 4 | 4 | 8 | 16 | 0 | 100 |

#### R7-DRA and R8-DRA outgoing Mti-DRA-2

| name | skid | R7-DRA-a | R7-DRA-b | R7-DRA-c | R8-DRA-a | R8-DRA-b | R8-DRA-c | total | %R7-DRA | %R8-DRA |
|---|---|---|---|---|---|---|---|---|---|---|
| Putative Mti_DRA_2 11995289 EK | 11995288 | 5 | 0 | 0 | 0 | 0 | 0 | 5 | 100 | 0 |
| Putative Mti_DRA_2 11693572 PA | 11693571 | 0 | 0 | 4 | 0 | 0 | 0 | 4 | 100 | 0 |
| Putative Mti_DRA_2 11993445 EK | 11993444 | 3 | 0 | 0 | 0 | 0 | 0 | 3 | 100 | 0 |
| Putative Mti_DRA_2 11903803 GS | 11903802 | 0 | 3 | 0 | 0 | 0 | 0 | 3 | 100 | 0 |
| Total | | 8 | 3 | 4 | 0 | 0 | 0 | 15 | 100 | 0 |

#### R7-DRA and R8-DRA outgoing Mi1

| name | skid | R7-DRA-a | R7-DRA-b | R7-DRA-c | R8-DRA-a | R8-DRA-b | R8-DRA-c | total | %R7-DRA | %R8-DRA |
|---|---|---|---|---|---|---|---|---|---|---|
| Putative Mi1 11829851 HL | 11829850 | 0 | 0 | 1 | 0 | 0 | 5 | 6 | 16.7 | 83.3 |
| Putative Mi1 13294452 EK | 13294451 | 3 | 0 | 0 | 2 | 0 | 0 | 5 | 60.0 | 40.0 |
| Putative Mi1 11660604824 CL | 14811021 | 0 | 2 | 0 | 0 | 2 | 0 | 4 | 50.0 | 50.0 |
| Total | | 3 | 2 | 1 | 2 | 2 | 5 | 15 | 40.0 | 60.0 |

#### R7-DRA and R8-DRA outgoing MeTu

| name | skid | R7-DRA-a | R7-DRA-b | R7-DRA-c | R8-DRA-a | R8-DRA-b | R8-DRA-c | total | %R7-DRA | %R8-DRA |
|---|---|---|---|---|---|---|---|---|---|---|
| Putative MeTu 16730683 AT | 16730682 | 0 | 0 | 8 | 0 | 0 | 0 | 8 | 100 | 0 |
| Putative MeTu 14888321 AT | 14888320 | 0 | 0 | 5 | 0 | 0 | 0 | 5 | 100 | 0 |
| Total | | 0 | 0 | 13 | 0 | 0 | 0 | 13 | 100 | 0 |

#### R7-DRA and R8-DRA outgoing Tm5-like

| name | skid | R7-DRA-a | R7-DRA-b | R7-DRA-c | R8-DRA-a | R8-DRA-b | R8-DRA-c | total | %R7-DRA | %R8-DRA |
|---|---|---|---|---|---|---|---|---|---|---|
| Putative Tm5-like 12016007 EK | 12016006 | 0 | 0 | 0 | 0 | 0 | 12 | 12 | 0 | 100 |
| Total | | 0 | 0 | 0 | 0 | 0 | 12 | 12 | 0 | 100 |

#### R7-DRA and R8-DRA outgoing Mi9

| name | skid | R7-DRA-a | R7-DRA-b | R7-DRA-c | R8-DRA-a | R8-DRA-b | R8-DRA-c | total | %R7-DRA | %R8-DRA |
|---|---|---|---|---|---|---|---|---|---|---|
| Putative Mi9 12015967 EK | 12015966 | 0 | 0 | 0 | 0 | 0 | 9 | 9 | 0 | 100 |
| Putative Mi9 11918591 EK | 11918590 | 0 | 0 | 0 | 3 | 0 | 0 | 3 | 0 | 100 |
| Total | | 0 | 0 | 0 | 3 | 0 | 9 | 12 | 0 | 100 |

#### R7-DRA and R8-DRA outgoing Dm11

| name | skid | R7-DRA-a | R7-DRA-b | R7-DRA-c | R8-DRA-a | R8-DRA-b | R8-DRA-c | total | %R7-DRA | %R8-DRA |
|---|---|---|---|---|---|---|---|---|---|---|
| Putative Dm11 17161736 EK | 17161735 | 0 | 0 | 4 | 0 | 0 | 8 | 12 | 33.3 | 66.7 |
| Total | | 0 | 0 | 4 | 0 | 0 | 8 | 12 | 33.3 | 66.7 |

#### R7-DRA and R8-DRA outgoing aMe12

| name | skid | R7-DRA-a | R7-DRA-b | R7-DRA-c | R8-DRA-a | R8-DRA-b | R8-DRA-c | total | %R7-DRA | %R8-DRA |
|---|---|---|---|---|---|---|---|---|---|---|
| Putative aMe12 VPN ME.R to vACA OLCT bilateral 164545 GA ECM | 164544 | 0 | 1 | 2 | 0 | 1 | 0 | 4 | 75 | 25 |
| Total | | 0 | 1 | 2 | 0 | 1 | 0 | 4 | 75 | 25 |

#### R7-DRA and R8-DRA outgoing TmY

| name | skid | R7-DRA-a | R7-DRA-b | R7-DRA-c | R8-DRA-a | R8-DRA-b | R8-DRA-c | total | %R7-DRA | %R8-DRA |
|---|---|---|---|---|---|---|---|---|---|---|
| Putative TmY 12043438 EK | 12043437 | 0 | 0 | 0 | 3 | 0 | 0 | 3 | 0 | 100 |
| Total | | 0 | 0 | 0 | 3 | 0 | 0 | 3 | 0 | 100 |

#### R7-DRA and R8-DRA outgoing ML-VPN2

| name | skid | R7-DRA-a | R7-DRA-b | R7-DRA-c | R8-DRA-a | R8-DRA-b | R8-DRA-c | total | %R7-DRA | %R8-DRA |
|---|---|---|---|---|---|---|---|---|---|---|
| Putative ML_VPN2 11995243 EK | 11995242 | 3 | 0 | 0 | 0 | 0 | 0 | 3 | 100 | 0 |
| Total | | 3 | 0 | 0 | 0 | 0 | 0 | 3 | 100 | 0 |

#### R7-DRA and R8-DRA outgoing C2

| name | skid | R7-DRA-a | R7-DRA-b | R7-DRA-c | R8-DRA-a | R8-DRA-b | R8-DRA-c | total | %R7-DRA | %R8-DRA |
|---|---|---|---|---|---|---|---|---|---|---|
| Putative C2 11906498 GS | 11906497 | 0 | 2 | 0 | 0 | 1 | 0 | 3 | 66.7 | 33.3 |
| Total | | 0 | 2 | 0 | 0 | 1 | 0 | 3 | 66.7 | 33.3 |

#### R7-DRA and R8-DRA outgoing Identified-<3

| name | skid | R7-DRA-a | R7-DRA-b | R7-DRA-c | R8-DRA-a | R8-DRA-b | R8-DRA-c | total | %R7-DRA | %R8-DRA |
|---|---|---|---|---|---|---|---|---|---|---|
| Putative Tm5-like 12017355 EK | 12017354 | 0 | 0 | 2 | 0 | 0 | 0 | 2 | 100.0 | 0.0 |
| Putative Mi9 10655487 EK | 10655486 | 0 | 0 | 0 | 0 | 2 | 0 | 2 | 0.0 | 100.0 |
| Putative MeTu_DRA 11993247 EK | 15749075 | 2 | 0 | 0 | 0 | 0 | 0 | 2 | 100.0 | 0.0 |
| Putative MeTu 10806612 EK | 10806611 | 2 | 0 | 0 | 0 | 0 | 0 | 2 | 100.0 | 0.0 |
| Putative MeTu 10473597 EK | 10473596 | 0 | 0 | 2 | 0 | 0 | 0 | 2 | 100.0 | 0.0 |
| Putative L2 12043826 EK | 12043825 | 0 | 0 | 2 | 0 | 0 | 0 | 2 | 100.0 | 0.0 |
| Putative L2 11915907 EK | 11915906 | 1 | 0 | 0 | 1 | 0 | 0 | 2 | 50.0 | 50.0 |
| Putative Dm2 17254551 EK | 17254550 | 0 | 0 | 2 | 0 | 0 | 0 | 2 | 100.0 | 0.0 |
| Putative Dm2 14305319 EK | 14305318 | 0 | 0 | 0 | 2 | 0 | 0 | 2 | 0.0 | 100.0 |
| Putative Dm 16110050 EK | 16110049 | 0 | 2 | 0 | 0 | 0 | 0 | 2 | 100.0 | 0.0 |
| Putative Dm 11993804 EK | 11993803 | 1 | 0 | 0 | 1 | 0 | 0 | 2 | 50.0 | 50.0 |
| Putative Dm 10479098 EK | 10479097 | 2 | 0 | 0 | 0 | 0 | 0 | 2 | 100.0 | 0.0 |
| Putative Dm-DRA2 11981476 EK | 11981475 | 0 | 0 | 0 | 2 | 0 | 0 | 2 | 0.0 | 100.0 |
| Putative C2 12044213 EK | 12044212 | 0 | 0 | 1 | 0 | 0 | 1 | 2 | 50.0 | 50.0 |
| Putative aMe12 VPN ME.R to vACA OLCT bilateral 28842 AA GA | 28841 | 0 | 2 | 0 | 0 | 0 | 0 | 2 | 100.0 | 0.0 |
| Putative VPN_DRA ME.R 8656926956 EK | 15997281 | 0 | 1 | 0 | 0 | 0 | 0 | 1 | 100.0 | 0.0 |
| Putative VPN_DRA ME.R 16215492 EK | 16215491 | 0 | 1 | 0 | 0 | 0 | 0 | 1 | 100.0 | 0.0 |
| Putative VPN_DRA ME.R 12188203 EK | 12188202 | 1 | 0 | 0 | 0 | 0 | 0 | 1 | 100.0 | 0.0 |
| Putative VPN_DRA ME.R 10088063089 EK | 15984100 | 1 | 0 | 0 | 0 | 0 | 0 | 1 | 100.0 | 0.0 |
| Putative T2 17252542 EK | 17252541 | 0 | 0 | 1 | 0 | 0 | 0 | 1 | 100.0 | 0.0 |
| Putative Mt_DRA_2 12109027 EK | 12109026 | 1 | 0 | 0 | 0 | 0 | 0 | 1 | 100.0 | 0.0 |
| Putative Mt_DRA_2 11993433 EK | 11993432 | 1 | 0 | 0 | 0 | 0 | 0 | 1 | 100.0 | 0.0 |
| Putative Mt_DRA_2 11992771 EK | 11992770 | 1 | 0 | 0 | 0 | 0 | 0 | 1 | 100.0 | 0.0 |
| Putative Mi9 11947153 EK | 11947152 | 0 | 0 | 0 | 1 | 0 | 0 | 1 | 0.0 | 100.0 |
| Putative Mi4 12045603 EK | 12045602 | 0 | 0 | 0 | 0 | 1 | 0 | 1 | 0.0 | 100.0 |
| Putative MeTu_DRA 10820917 EK | 10820916 | 1 | 0 | 0 | 0 | 0 | 0 | 1 | 100.0 | 0.0 |
| Putative MeTu_DRA 10755132 EK | 10755131 | 1 | 0 | 0 | 0 | 0 | 0 | 1 | 100.0 | 0.0 |
| Putative MeTu 4735813145 EK | 15978559 | 0 | 1 | 0 | 0 | 0 | 0 | 1 | 100.0 | 0.0 |
| Putative MeTu 14726393 MF | 14726392 | 1 | 0 | 0 | 0 | 0 | 0 | 1 | 100.0 | 0.0 |
| Putative L3 6831613493 EK | 15970720 | 0 | 1 | 0 | 0 | 0 | 0 | 1 | 100.0 | 0.0 |
| Putative L3 17253296 EK | 17253295 | 0 | 0 | 0 | 0 | 0 | 1 | 1 | 0.0 | 100.0 |
| Putative Dm3 12018854 EK | 12018853 | 0 | 0 | 0 | 0 | 0 | 1 | 1 | 0.0 | 100.0 |
| Putative C3 12002623 EK | 12002622 | 0 | 0 | 0 | 0 | 0 | 1 | 1 | 0.0 | 100.0 |
| Total | | 16 | 8 | 10 | 7 | 3 | 4 | 48 | 70.8 | 29.2 |

#### R7-DRA and R8-DRA outgoing Unidentified->=3

| name | skid | R7-DRA-a | R7-DRA-b | R7-DRA-c | R8-DRA-a | R8-DRA-b | R8-DRA-c | total | %R7-DRA | %R8-DRA |
|---|---|---|---|---|---|---|---|---|---|---|
| Neuron 15007628 (not traceable) EK | 15007627 | 0 | 5 | 0 | 0 | 0 | 0 | 5 | 100 | 0 |
| 10655594 EK (untraceable) | 10655593 | 0 | 0 | 0 | 0 | 5 | 0 | 5 | 0 | 100 |
| Total | | 0 | 5 | 0 | 0 | 5 | 0 | 10 | 50 | 50 |

#### R7-DRA and R8-DRA outgoing Unidentified-<3

| name | skid | R7-DRA-a | R7-DRA-b | R7-DRA-c | R8-DRA-a | R8-DRA-b | R8-DRA-c | total | %R7-DRA | %R8-DRA |
|---|---|---|---|---|---|---|---|---|---|---|
| Neuron 15767888 | 15767887 | 0 | 2 | 0 | 0 | 0 | 0 | 2 | 100.0 | 0.0 |
| Neuron 14867404 | 14867403 | 0 | 2 | 0 | 0 | 0 | 0 | 2 | 100.0 | 0.0 |
| Neuron 14788296 EK | 14788295 | 0 | 2 | 0 | 0 | 0 | 0 | 2 | 100.0 | 0.0 |
| neuron 11994901 EK | 11994900 | 0 | 0 | 0 | 2 | 0 | 0 | 2 | 0.0 | 100.0 |
| neuron 11947310 EK (untraceable) | 11947309 | 0 | 0 | 0 | 2 | 0 | 0 | 2 | 0.0 | 100.0 |
| neuron 11918119 EK (untraceable) | 11918118 | 0 | 0 | 0 | 2 | 0 | 0 | 2 | 0.0 | 100.0 |
| neuron 11918097 EK (untraceable) | 11918096 | 0 | 0 | 0 | 2 | 0 | 0 | 2 | 0.0 | 100.0 |
| neuron 11904799 GS (untraceable) | 11904798 | 0 | 2 | 0 | 0 | 0 | 0 | 2 | 100.0 | 0.0 |
| neuron 11904043 GS (not traceable) | 11904042 | 0 | 2 | 0 | 0 | 0 | 0 | 2 | 100.0 | 0.0 |
| neuron 11903818 GS (not traceable) | 11903817 | 0 | 2 | 0 | 0 | 0 | 0 | 2 | 100.0 | 0.0 |
| Neuron 14991389 | 14991388 | 0 | 1 | 0 | 0 | 0 | 0 | 1 | 100.0 | 0.0 |
| Neuron 14798841 | 14798840 | 0 | 1 | 0 | 0 | 0 | 0 | 1 | 100.0 | 0.0 |
| Neuron 14315234 | 14315233 | 0 | 0 | 0 | 0 | 1 | 0 | 1 | 0.0 | 100.0 |
| Neuron 14081457 GS | 14081456 | 0 | 1 | 0 | 0 | 0 | 0 | 1 | 100.0 | 0.0 |
| Neuron 13938782 | 13938781 | 0 | 1 | 0 | 0 | 0 | 0 | 1 | 100.0 | 0.0 |
| Neuron 13433448 EK (untraceable) | 13433447 | 1 | 0 | 0 | 0 | 0 | 0 | 1 | 100.0 | 0.0 |
| neuron 13179813 EK | 13179812 | 0 | 1 | 0 | 0 | 0 | 0 | 1 | 100.0 | 0.0 |
| neuron 12043433 EK | 12043432 | 0 | 0 | 0 | 1 | 0 | 0 | 1 | 0.0 | 100.0 |
| neuron 12019263 | 12019262 | 0 | 0 | 1 | 0 | 0 | 0 | 1 | 100.0 | 0.0 |
| neuron 12019256 | 12019255 | 0 | 0 | 0 | 0 | 0 | 1 | 1 | 0.0 | 100.0 |
| neuron 12018755 | 12018754 | 0 | 0 | 0 | 0 | 0 | 1 | 1 | 0.0 | 100.0 |
| neuron 12017797 | 12017796 | 0 | 0 | 1 | 0 | 0 | 0 | 1 | 100.0 | 0.0 |
| neuron 12017523 | 12017522 | 0 | 0 | 1 | 0 | 0 | 0 | 1 | 100.0 | 0.0 |
| neuron 12017236 | 12017235 | 0 | 0 | 0 | 0 | 0 | 1 | 1 | 0.0 | 100.0 |
| neuron 12017107 | 12017106 | 0 | 0 | 0 | 0 | 0 | 1 | 1 | 0.0 | 100.0 |
| neuron 12016873 | 12016872 | 0 | 0 | 0 | 0 | 0 | 1 | 1 | 0.0 | 100.0 |
| neuron 12016687 | 12016686 | 0 | 0 | 1 | 0 | 0 | 0 | 1 | 100.0 | 0.0 |
| neuron 12016413 | 12016412 | 0 | 0 | 0 | 0 | 0 | 1 | 1 | 0.0 | 100.0 |
| neuron 12015890 | 12015889 | 0 | 0 | 0 | 0 | 0 | 1 | 1 | 0.0 | 100.0 |
| neuron 12015018 | 12015017 | 0 | 0 | 1 | 0 | 0 | 0 | 1 | 100.0 | 0.0 |
| neuron 12014001 | 12014000 | 0 | 0 | 1 | 0 | 0 | 0 | 1 | 100.0 | 0.0 |
| neuron 12013715 | 12013714 | 0 | 0 | 1 | 0 | 0 | 0 | 1 | 100.0 | 0.0 |
| neuron 12002539 | 12002538 | 0 | 0 | 1 | 0 | 0 | 0 | 1 | 100.0 | 0.0 |
| neuron 12002237 | 12002236 | 0 | 0 | 1 | 0 | 0 | 0 | 1 | 100.0 | 0.0 |
| neuron 12002170 | 12002169 | 0 | 0 | 1 | 0 | 0 | 0 | 1 | 100.0 | 0.0 |
| neuron 12002160 | 12002159 | 0 | 0 | 1 | 0 | 0 | 0 | 1 | 100.0 | 0.0 |
| neuron 11995171 EK | 11995170 | 1 | 0 | 0 | 0 | 0 | 0 | 1 | 100.0 | 0.0 |
| neuron 11995069 EK | 11995068 | 1 | 0 | 0 | 0 | 0 | 0 | 1 | 100.0 | 0.0 |
| neuron 11995028 EK | 11995027 | 1 | 0 | 0 | 0 | 0 | 0 | 1 | 100.0 | 0.0 |
| neuron 11994592 EK | 11994591 | 1 | 0 | 0 | 0 | 0 | 0 | 1 | 100.0 | 0.0 |
| neuron 11994244 EK | 11994243 | 1 | 0 | 0 | 0 | 0 | 0 | 1 | 100.0 | 0.0 |
| neuron 11993824 EK | 11993823 | 1 | 0 | 0 | 0 | 0 | 0 | 1 | 100.0 | 0.0 |
| neuron 11992865 EK | 11992864 | 1 | 0 | 0 | 0 | 0 | 0 | 1 | 100.0 | 0.0 |
| neuron 11981310 EK | 11981309 | 0 | 0 | 0 | 1 | 0 | 0 | 1 | 0.0 | 100.0 |
| neuron 11981299 EK | 11981298 | 0 | 0 | 0 | 1 | 0 | 0 | 1 | 0.0 | 100.0 |
| neuron 11908911 GS | 11908910 | 0 | 0 | 0 | 0 | 1 | 0 | 1 | 0.0 | 100.0 |
| neuron 11908664 | 11908663 | 0 | 1 | 0 | 0 | 0 | 0 | 1 | 100.0 | 0.0 |
| neuron 11904702 | 11904701 | 0 | 1 | 0 | 0 | 0 | 0 | 1 | 100.0 | 0.0 |
| neuron 11904687 | 11904686 | 0 | 1 | 0 | 0 | 0 | 0 | 1 | 100.0 | 0.0 |
| neuron 11904395 | 11904394 | 0 | 1 | 0 | 0 | 0 | 0 | 1 | 100.0 | 0.0 |
| neuron 11904241 | 11904240 | 0 | 1 | 0 | 0 | 0 | 0 | 1 | 100.0 | 0.0 |
| neuron 11904203 | 11904202 | 0 | 1 | 0 | 0 | 0 | 0 | 1 | 100.0 | 0.0 |
| neuron 11904153 | 11904152 | 0 | 1 | 0 | 0 | 0 | 0 | 1 | 100.0 | 0.0 |
| neuron 11904033 | 11904032 | 0 | 1 | 0 | 0 | 0 | 0 | 1 | 100.0 | 0.0 |
| neuron 11903739 | 11903738 | 0 | 1 | 0 | 0 | 0 | 0 | 1 | 100.0 | 0.0 |
| neuron 11902130 | 11902129 | 0 | 1 | 0 | 0 | 0 | 0 | 1 | 100.0 | 0.0 |
| neuron 11829995 | 11829994 | 0 | 0 | 0 | 0 | 0 | 1 | 1 | 0.0 | 100.0 |
| Total | | 8 | 27 | 11 | 11 | 2 | 8 | 67 | 68.7 | 31.3 |

#### R7-DRA and R8-DRA incoming Dm9

| name | skid | R7-DRA-a | R7-DRA-b | R7-DRA-c | R8-DRA-a | R8-DRA-b | R8-DRA-c | total | %R7-DRA | %R8-DRA |
|---|---|---|---|---|---|---|---|---|---|---|
| Putative Dm9 11916196 EK | 11916195 | 39 | 0 | 0 | 34 | 0 | 0 | 73 | 53.4 | 46.6 |
| Putative Dm9 12013130 EK | 12013129 | 0 | 0 | 34 | 0 | 0 | 35 | 69 | 49.3 | 50.7 |
| Putative Dm9 10657501 GS | 10657500 | 0 | 30 | 0 | 0 | 36 | 0 | 66 | 45.5 | 54.5 |
| Putative Dm9 10655889 EK | 10655888 | 0 | 2 | 0 | 0 | 1 | 0 | 3 | 66.7 | 33.3 |
| Total | | 39 | 32 | 34 | 34 | 37 | 35 | 211 | 49.8 | 50.2 |

#### R7-DRA and R8-DRA incoming R8-DRA

| name | skid | R7-DRA-a | R7-DRA-b | R7-DRA-c | R8-DRA-a | R8-DRA-b | R8-DRA-c | total | %R7-DRA | %R8-DRA |
|---|---|---|---|---|---|---|---|---|---|---|
| Putative R8_DRA 10300964 TO | 10300963 | 0 | 38 | 0 | 0 | 0 | 0 | 38 | 100.0 | 0.0 |
| Putative R8_DRA 11728828 TO | 11728827 | 0 | 0 | 37 | 0 | 0 | 0 | 37 | 100.0 | 0.0 |
| Putative R8_DRA 10190509 TO | 10190508 | 33 | 0 | 0 | 1 | 0 | 0 | 34 | 97.1 | 2.9 |
| Total | | 33 | 38 | 37 | 1 | 0 | 0 | 109 | 99.1 | 0.9 |

#### R7-DRA and R8-DRA incoming R7-DRA

| name | skid | R7-DRA-a | R7-DRA-b | R7-DRA-c | R8-DRA-a | R8-DRA-b | R8-DRA-c | total | %R7-DRA | %R8-DRA |
|---|---|---|---|---|---|---|---|---|---|---|
| Putative R7_DRA 11728780 TO | 11728779 | 0 | 0 | 1 | 0 | 0 | 15 | 16 | 6.2 | 93.8 |
| Putative R7_DRA 10300950 TO | 10300949 | 0 | 0 | 0 | 0 | 16 | 0 | 16 | 0.0 | 100.0 |
| Putative R7_DRA 10191736 TO | 10191735 | 0 | 0 | 0 | 10 | 0 | 0 | 10 | 0.0 | 100.0 |
| Total | | 0 | 0 | 1 | 10 | 16 | 15 | 42 | 2.4 | 97.6 |

#### R7-DRA and R8-DRA incoming C2

| name | skid | R7-DRA-a | R7-DRA-b | R7-DRA-c | R8-DRA-a | R8-DRA-b | R8-DRA-c | total | %R7-DRA | %R8-DRA |
|---|---|---|---|---|---|---|---|---|---|---|
| Putative C2 12044213 EK | 12044212 | 0 | 0 | 2 | 0 | 0 | 3 | 5 | 40.0 | 60.0 |
| Putative C2 11906498 GS | 11906497 | 0 | 2 | 0 | 0 | 1 | 0 | 3 | 66.7 | 33.3 |
| Total | | 0 | 2 | 2 | 0 | 1 | 3 | 8 | 50.0 | 50.0 |

#### R7-DRA and R8-DRA incoming Mi15

| name | skid | R7-DRA-a | R7-DRA-b | R7-DRA-c | R8-DRA-a | R8-DRA-b | R8-DRA-c | total | %R7-DRA | %R8-DRA |
|---|---|---|---|---|---|---|---|---|---|---|
| Putative Mi15 10655927 TO | 10655926 | 0 | 2 | 0 | 0 | 2 | 0 | 4 | 50 | 50 |
| Total | | 0 | 2 | 0 | 0 | 2 | 0 | 4 | 50 | 50 |

#### R7-DRA and R8-DRA incoming Identified-<3

| name | skid | R7-DRA-a | R7-DRA-b | R7-DRA-c | R8-DRA-a | R8-DRA-b | R8-DRA-c | total | %R7-DRA | %R8-DRA |
|---|---|---|---|---|---|---|---|---|---|---|
| Putative Dm-DRA1 11993077 EK | 11993076 | 1 | 0 | 0 | 0 | 0 | 0 | 1 | 100.0 | 0.0 |
| Putative Dm-DRA1 11993696 EK | 11993695 | 1 | 0 | 0 | 0 | 0 | 0 | 1 | 100.0 | 0.0 |
| Putative Dm9 14933803 EK | 14933802 | 0 | 1 | 0 | 0 | 0 | 0 | 1 | 100.0 | 0.0 |
| Putative L3 12018070 EK | 12018069 | 0 | 0 | 1 | 0 | 0 | 0 | 1 | 100.0 | 0.0 |
| Putative Dm-DRA1 16766813 EK | 16766812 | 0 | 0 | 1 | 0 | 0 | 0 | 1 | 100.0 | 0.0 |
| Putative Dm-DRA1 17155322 EK | 17155321 | 0 | 0 | 1 | 0 | 0 | 0 | 1 | 100.0 | 0.0 |
| Putative L1 11915726 EK | 11915725 | 0 | 0 | 0 | 1 | 0 | 0 | 1 | 0.0 | 100.0 |
| Putative Mi15 11916191 EK | 11916190 | 0 | 0 | 0 | 1 | 0 | 0 | 1 | 0.0 | 100.0 |
| Putative L3 10653986 EK | 10653985 | 0 | 0 | 0 | 0 | 1 | 0 | 1 | 0.0 | 100.0 |
| Total | | 2 | 1 | 3 | 2 | 1 | 0 | 9 | 66.7 | 33.3 |

#### R7-DRA and R8-DRA incoming Unidentified-<3

| name | skid | R7-DRA-a | R7-DRA-b | R7-DRA-c | R8-DRA-a | R8-DRA-b | R8-DRA-c | total | %R7-DRA | %R8-DRA |
|---|---|---|---|---|---|---|---|---|---|---|
| neuron 11915809 | 11915808 | 1 | 0 | 0 | 1 | 0 | 0 | 2 | 50.0 | 50.0 |
| neuron 11903225 | 11903224 | 0 | 1 | 0 | 0 | 0 | 0 | 1 | 100.0 | 0.0 |
| Total | | 1 | 1 | 0 | 1 | 0 | 0 | 3 | 66.7 | 33.3 |
