## Supplementary File 2 for "Synaptic targets of photoreceptors specialized to detect color and skylight polarization in *Drosophila*"

### Supplementary File 2: Gallery plots of all seed column R7, R8, R7-DRA and R8-DRA target cells by type

### Central seed column Dm9

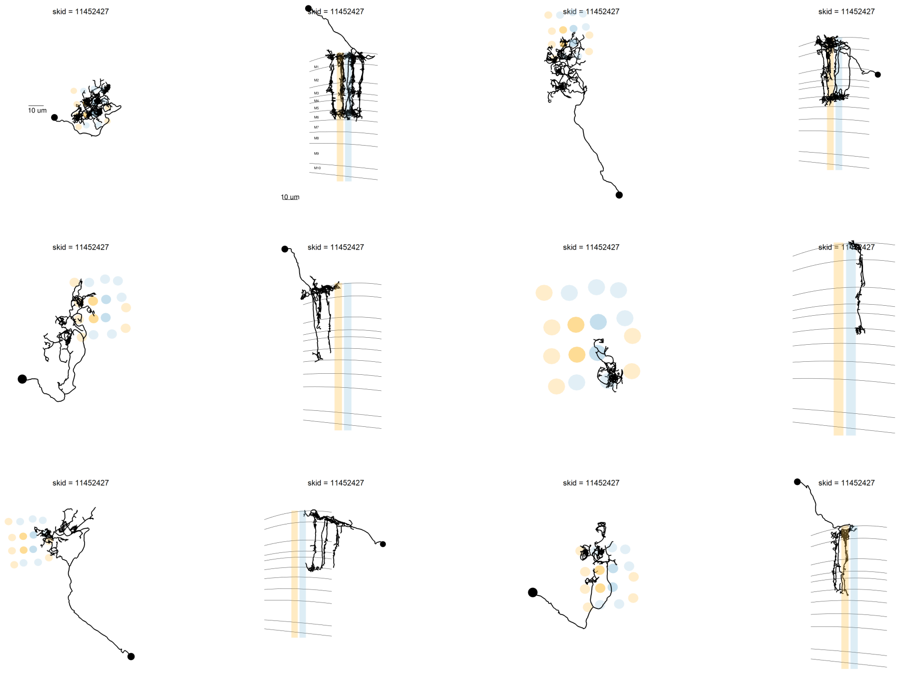

[1] 6 cells

### Central seed column Dm8

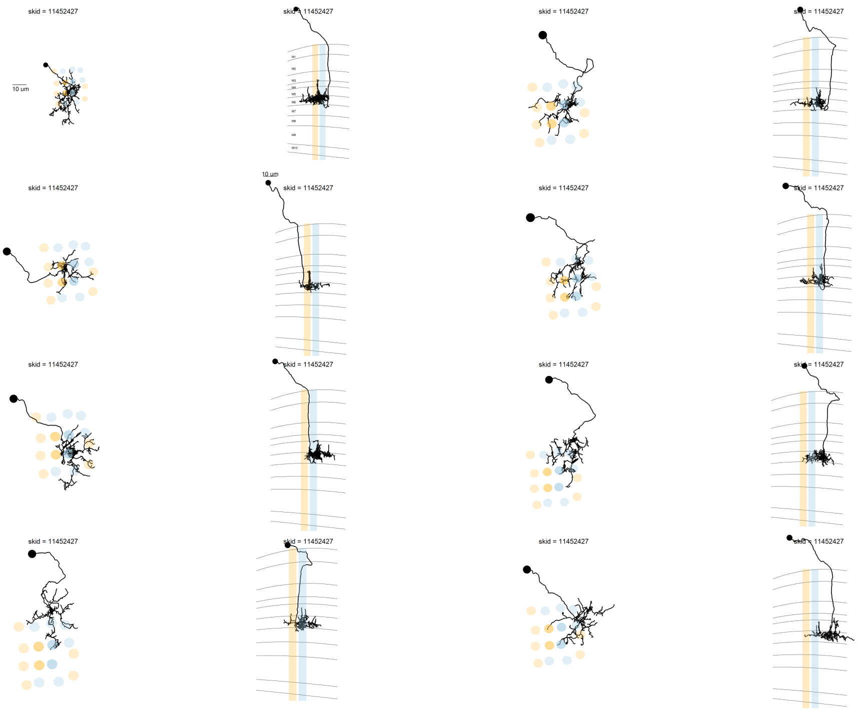

[1] 15 cells

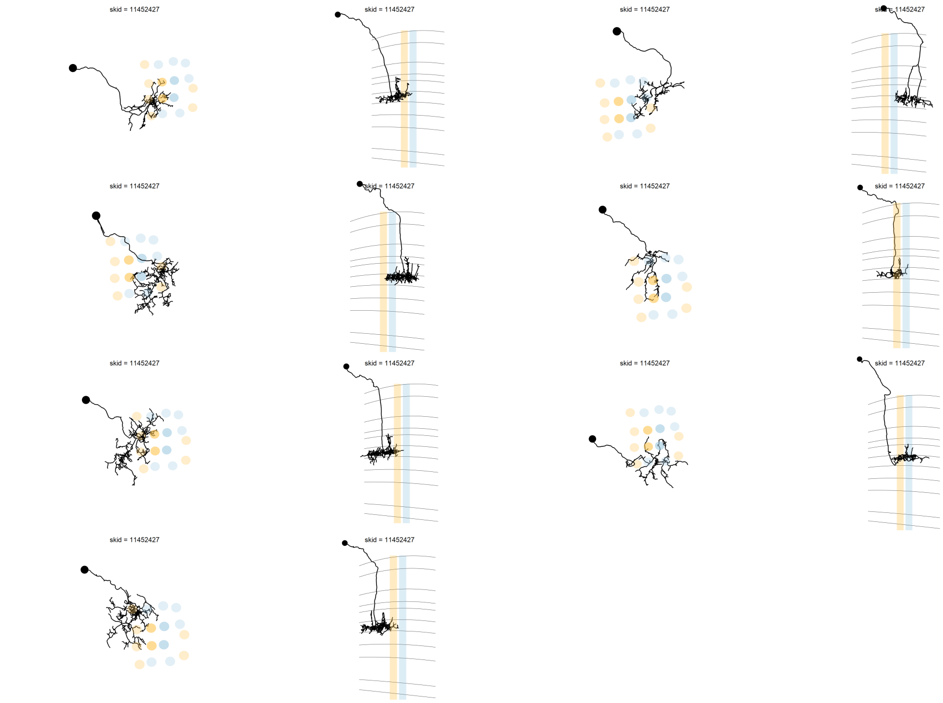
skid = 11452427
skid = 11452427
skid = 11452427
skid = 11452427
skid = 11452427
skid = 11452427
skid = 11452427
skid = 11452427
skid = 11452427
skid = 11452427
skid = 11452427
skid = 11452427
skid = 11452427
skid = 11452427

[1] 15 cells

### Central seed column MeTu

skid = 11452427

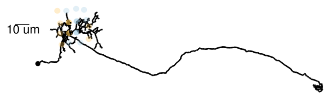

skid = 11452427

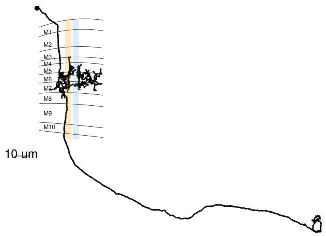

skid = 11452427

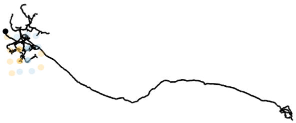

skid = 11452427

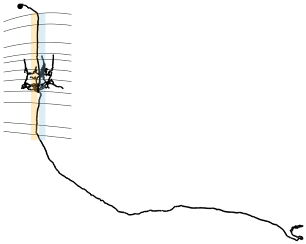

[1] 7 cells

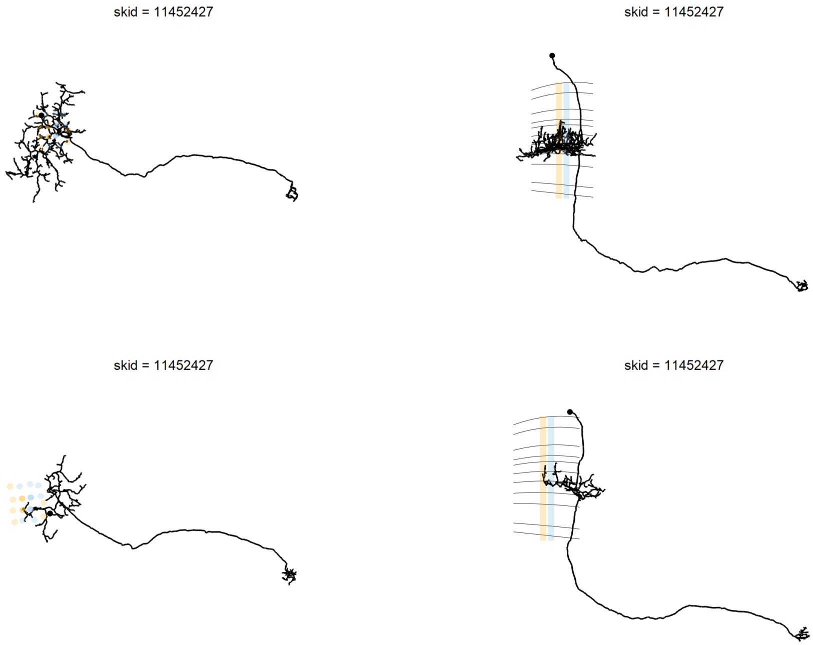

[1] 7 cells

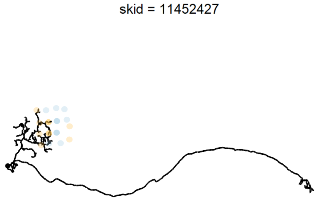

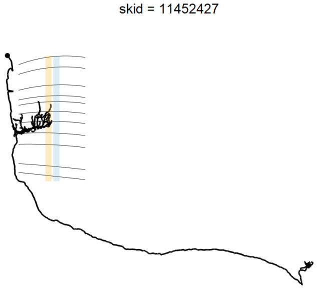

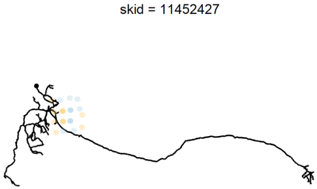

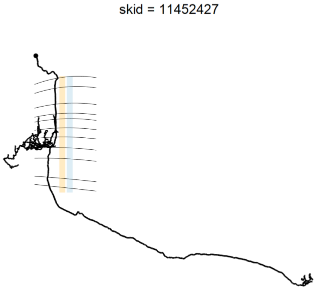

[1] 7 cells

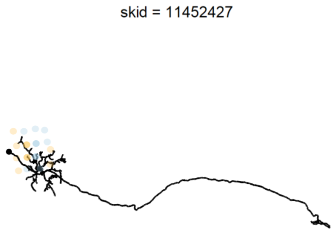

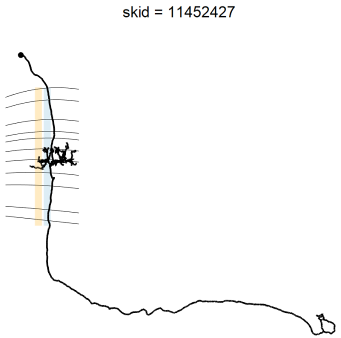

[1] 7 cells

### Central seed column R7

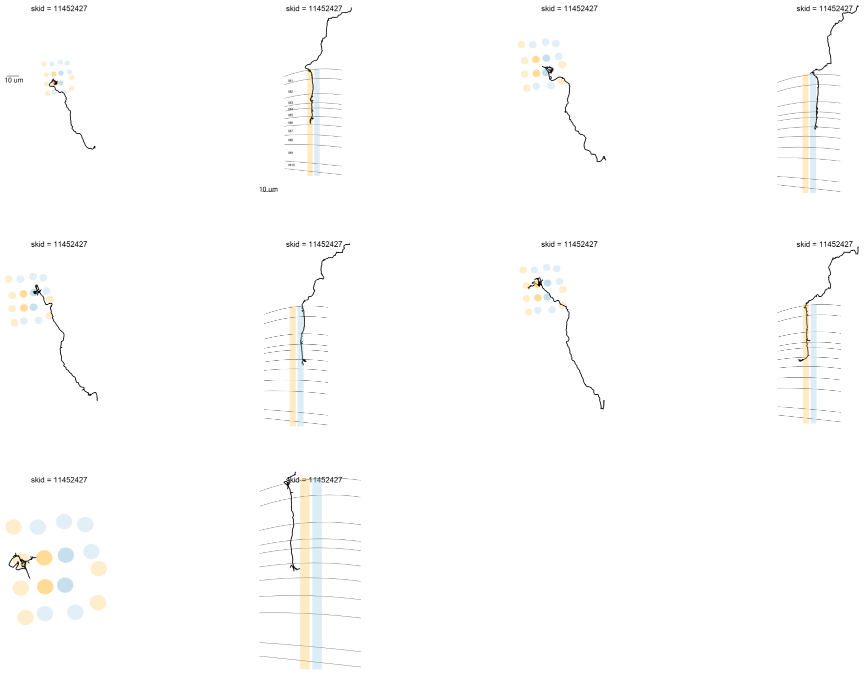

[1] 5 cells

### Central seed column Tm5c

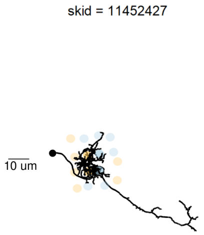

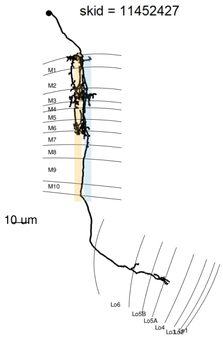

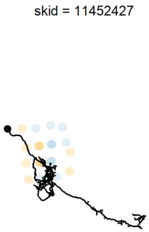

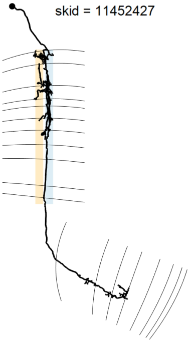

[1] 6 cells

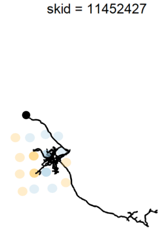

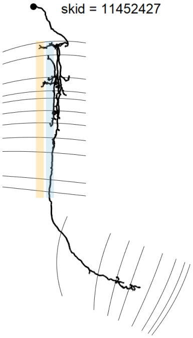

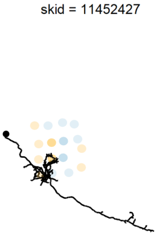

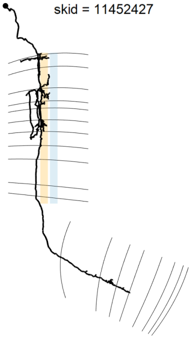

[1] 6 cells

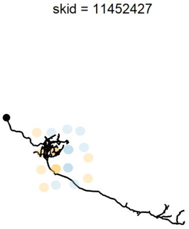

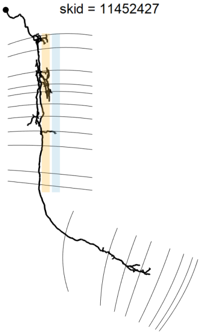

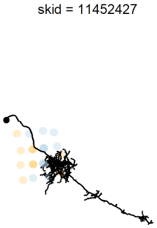

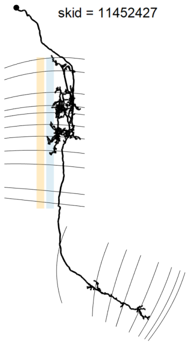

[1] 6 cells

### Central seed column Tm20

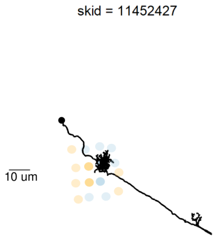

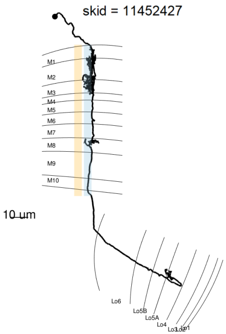

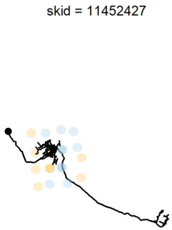

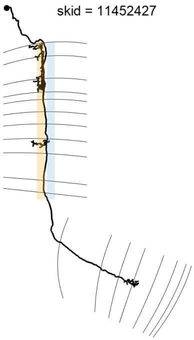

[1] 4 cells

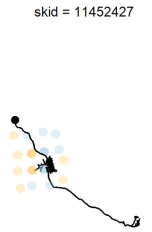

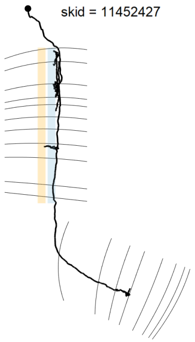

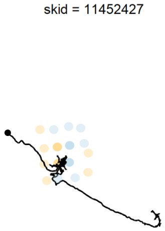

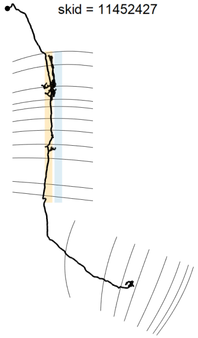

[1] 4 cells

### Central seed column Mi15

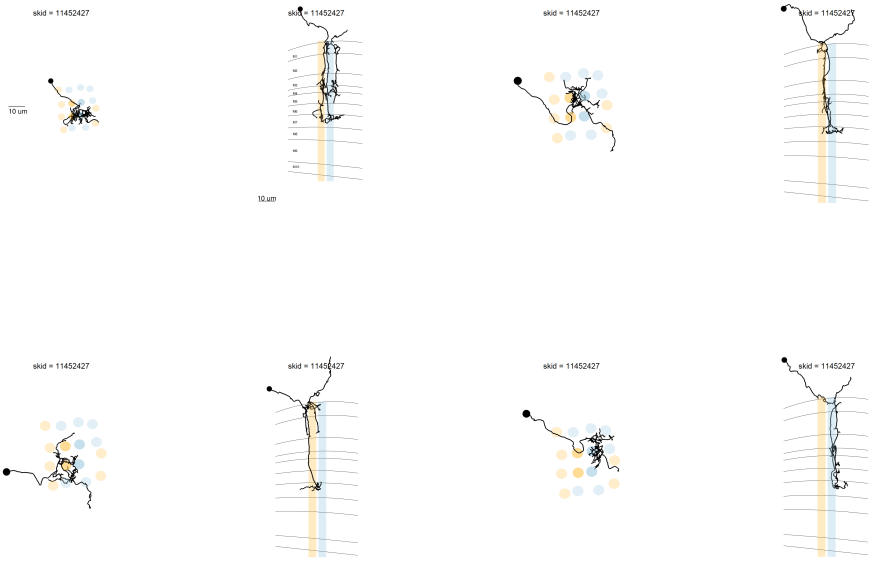

[1] 4 cells

### Central seed column Mi4

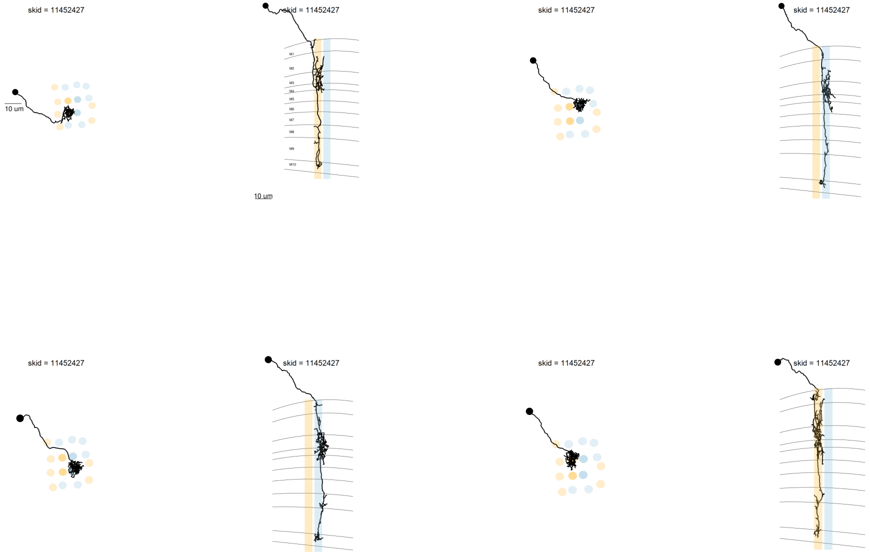

[1] 4 cells

### Central seed column ML1

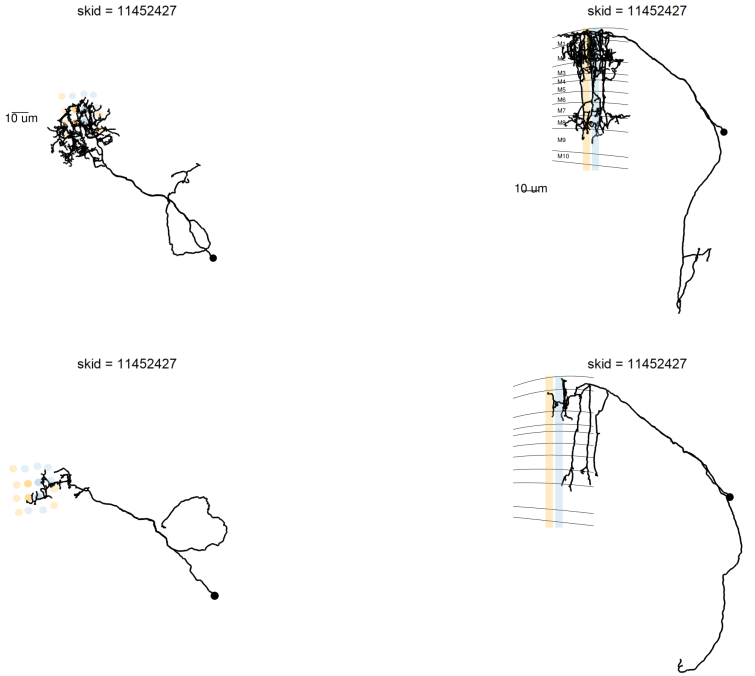

[1] 4 cells

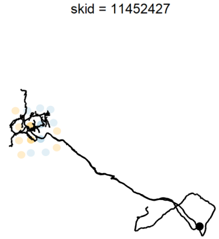

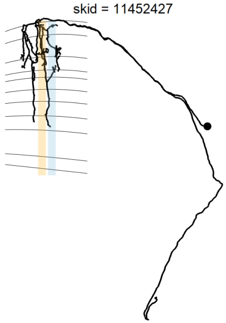

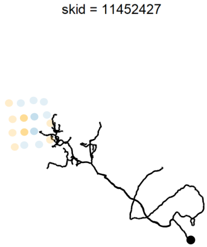

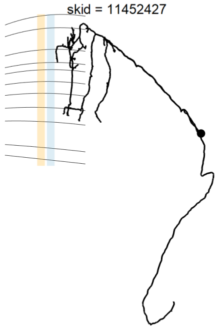

[1] 4 cells

### Central seed column Dm2

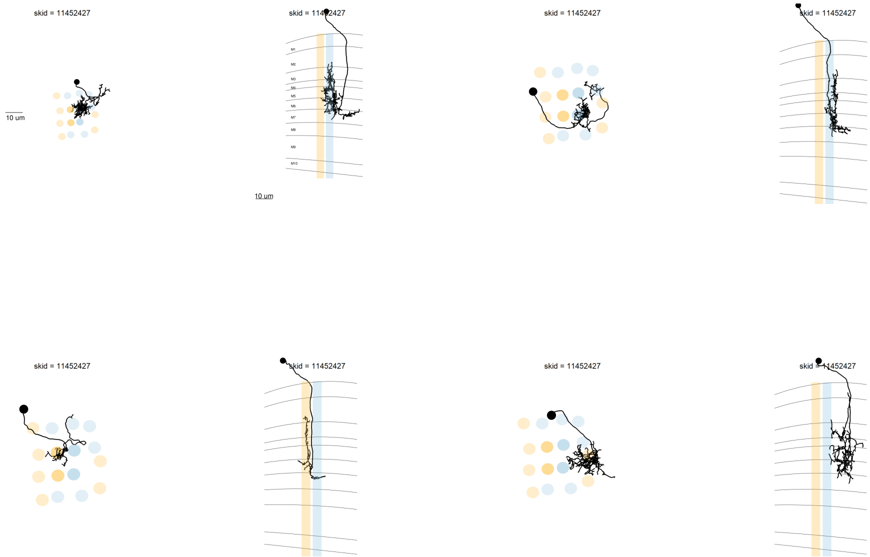

[1] 4 cells

### Central seed column Dm11

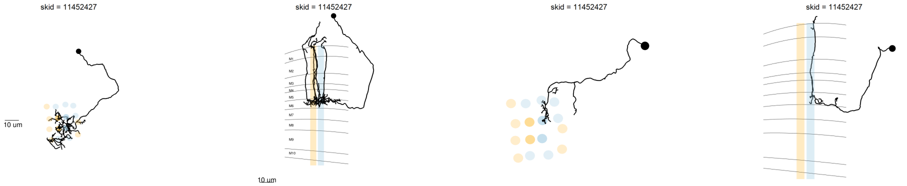

[1] 2 cells

### Central seed column L3

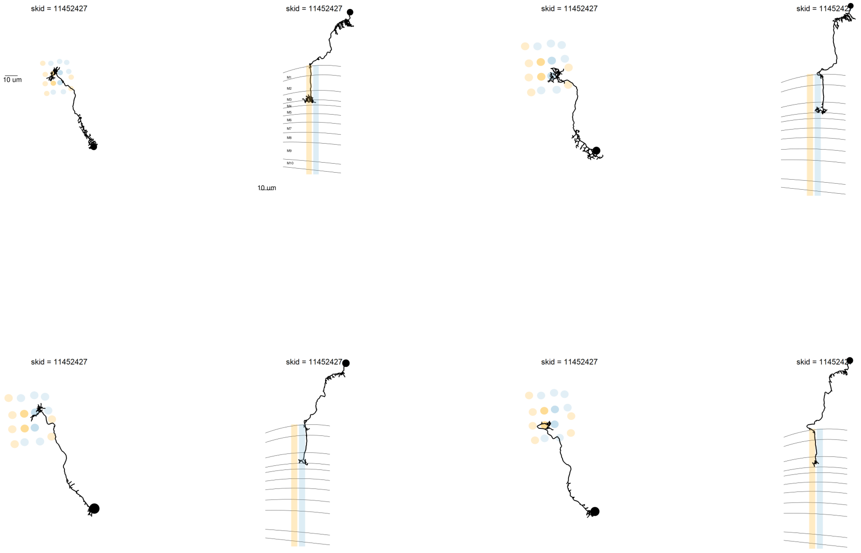

[1] 4 cells

### Central seed column Mi1

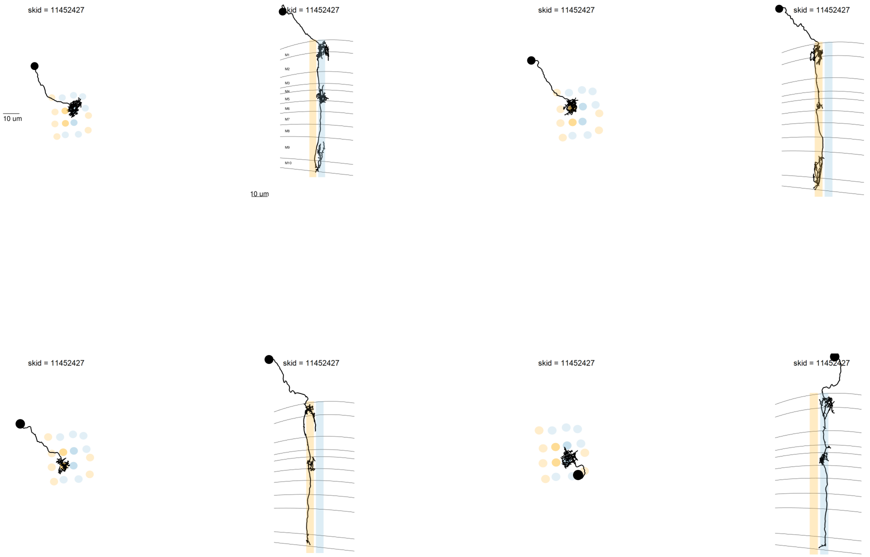

[1] 4 cells

### Central seed column R8

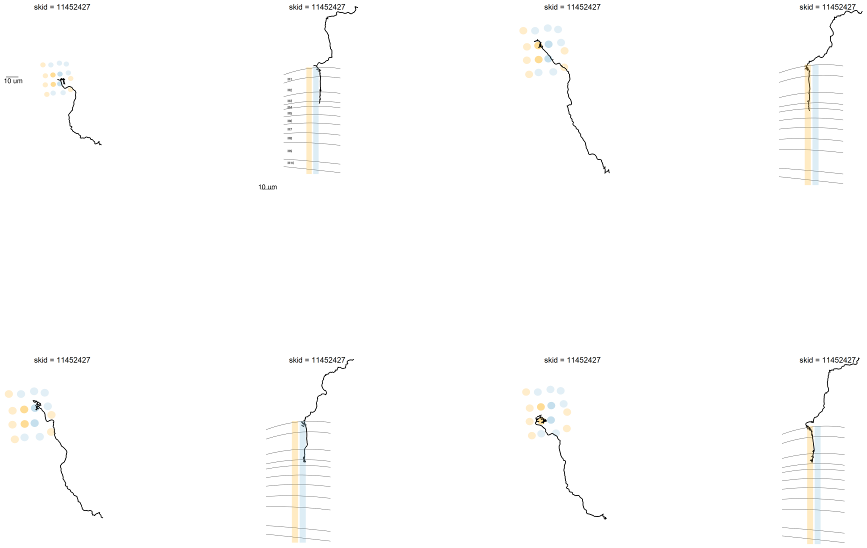

[1] 4 cells

### Central seed column Tm5a

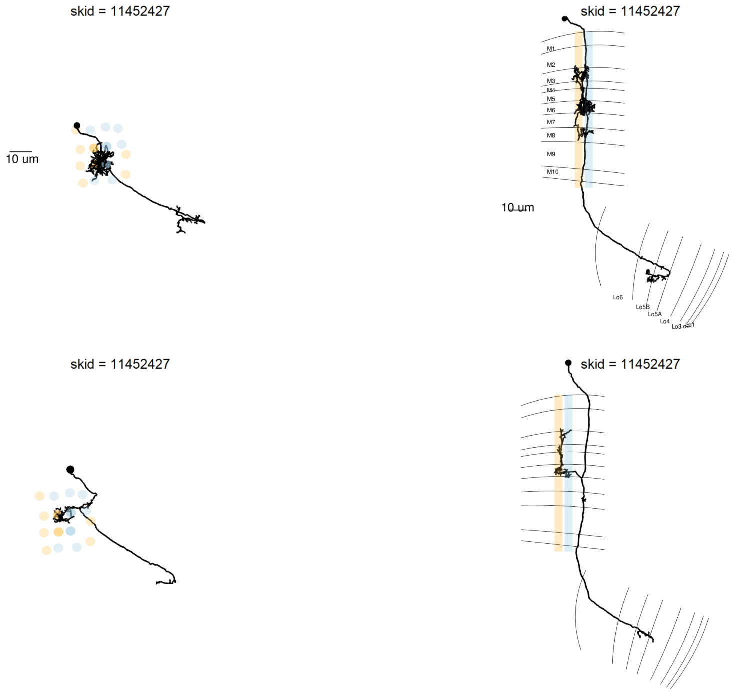

[1] 2 cells

### Central seed column Tm5b

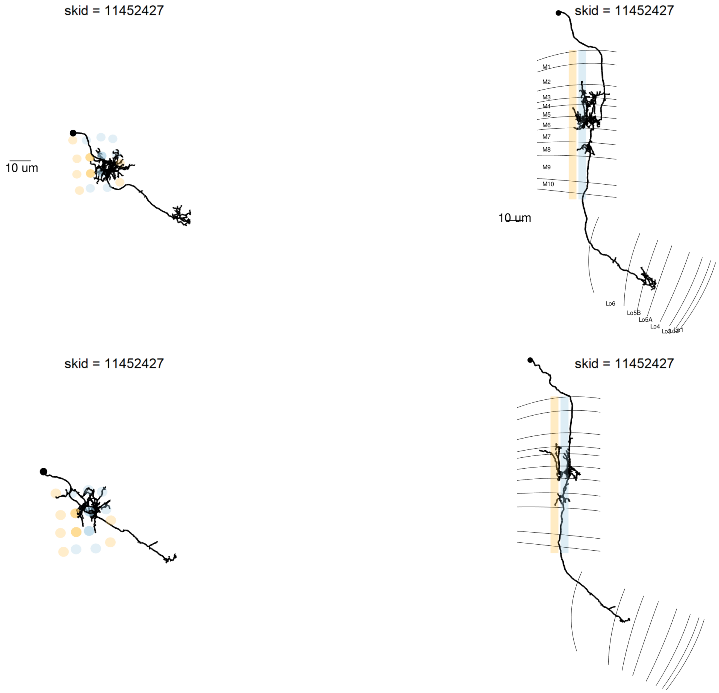

[1] 2 cells

### Central seed column Tm

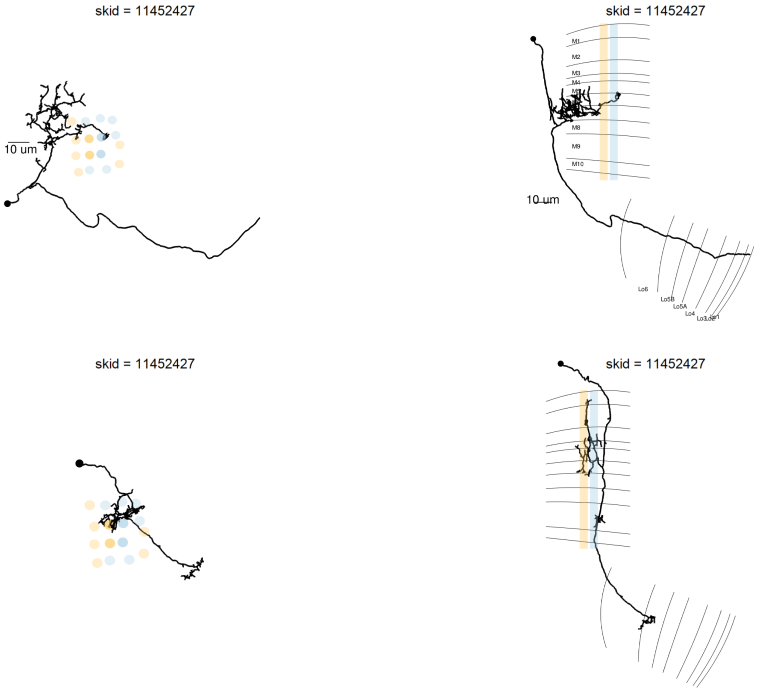

[1] 9 cells

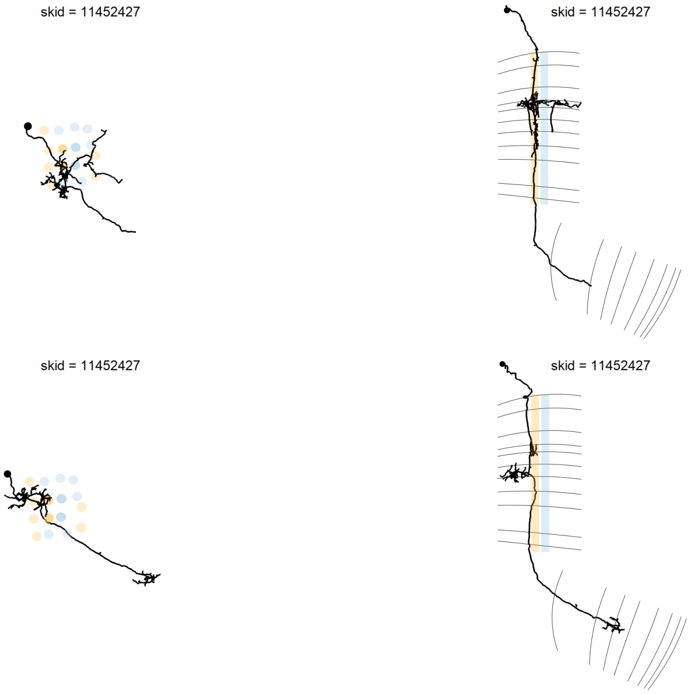

[1] 9 cells

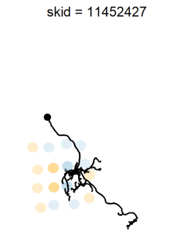

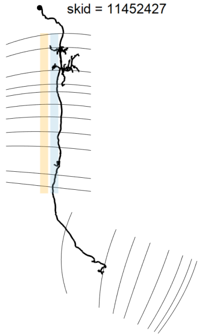

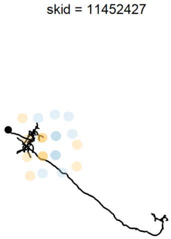

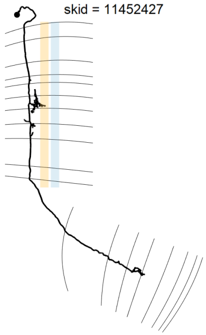

[1] 9 cells

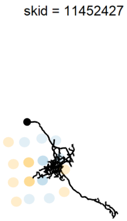

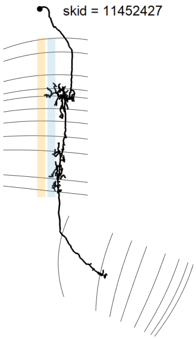

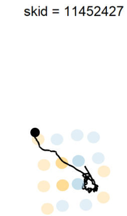

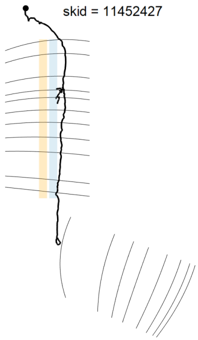

[1] 9 cells

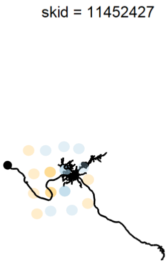

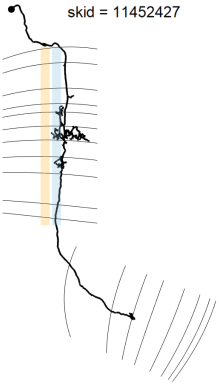

[1] 9 cells

### Central seed column Tm5b-like

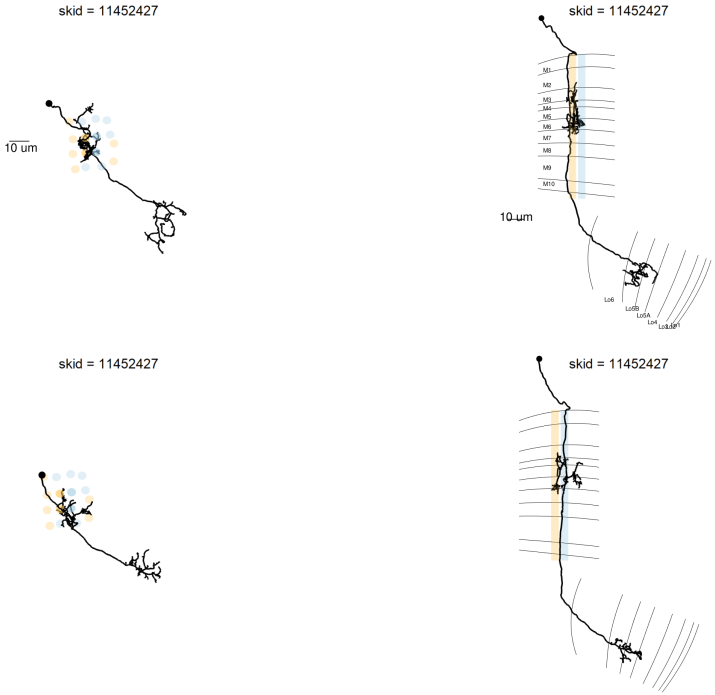

[1] 3 cells

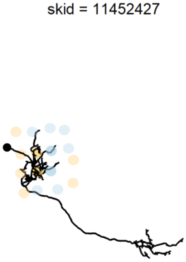

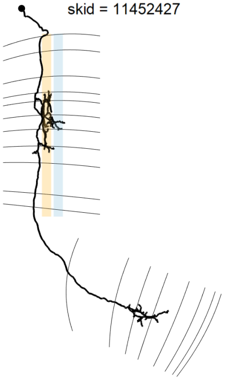

[1] 3 cells

### Central seed column Mi9

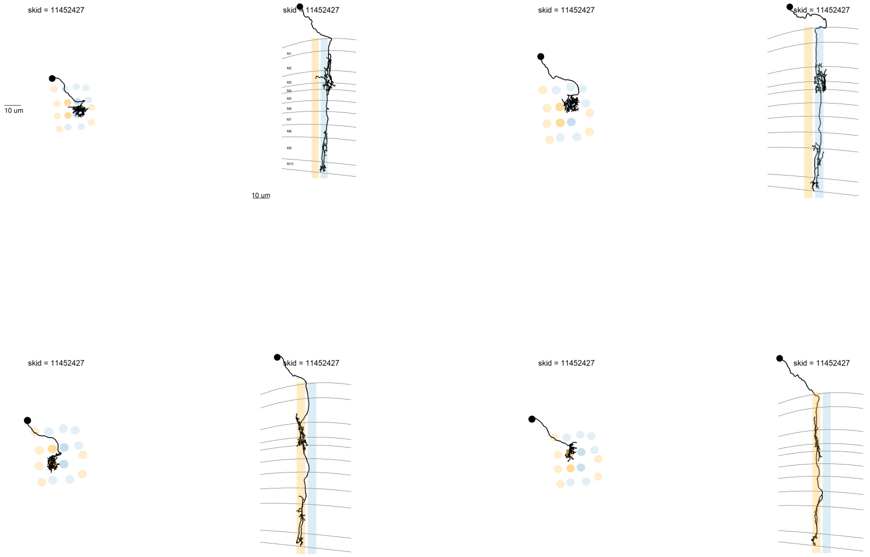

[1] 4 cells

### Central seed column L1

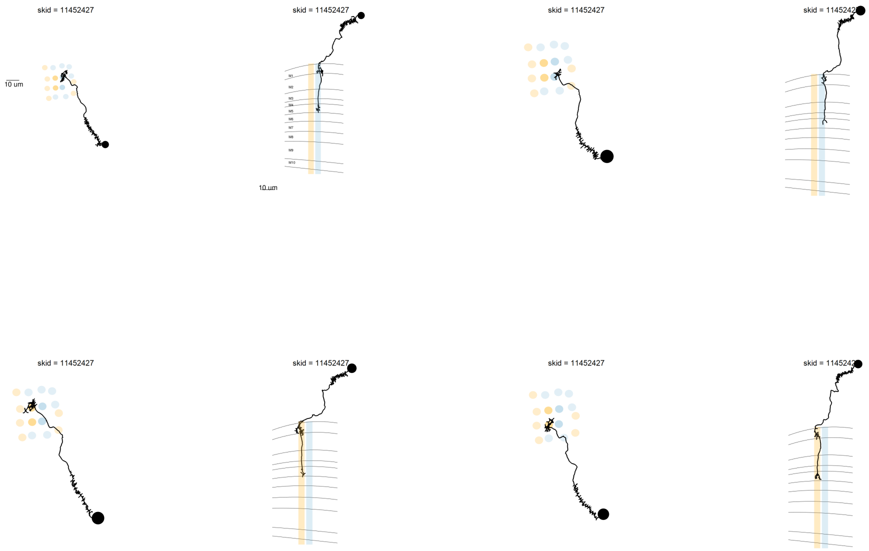

[1] 4 cells

### Central seed column aMe12

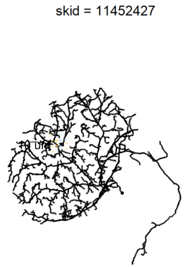

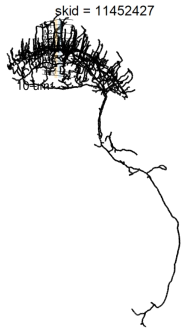

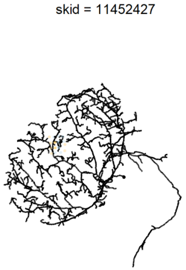

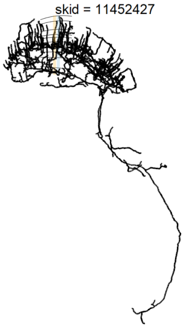

[1] 3 cells

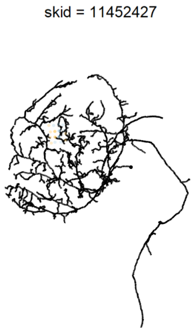

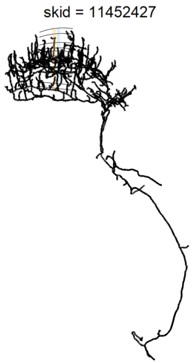

[1] 3 cells

### Central seed column Dm

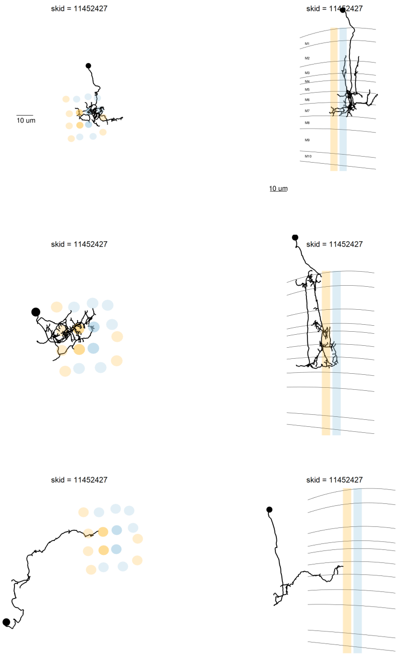

[1] 5 cells

### Central seed column ML-VPN1

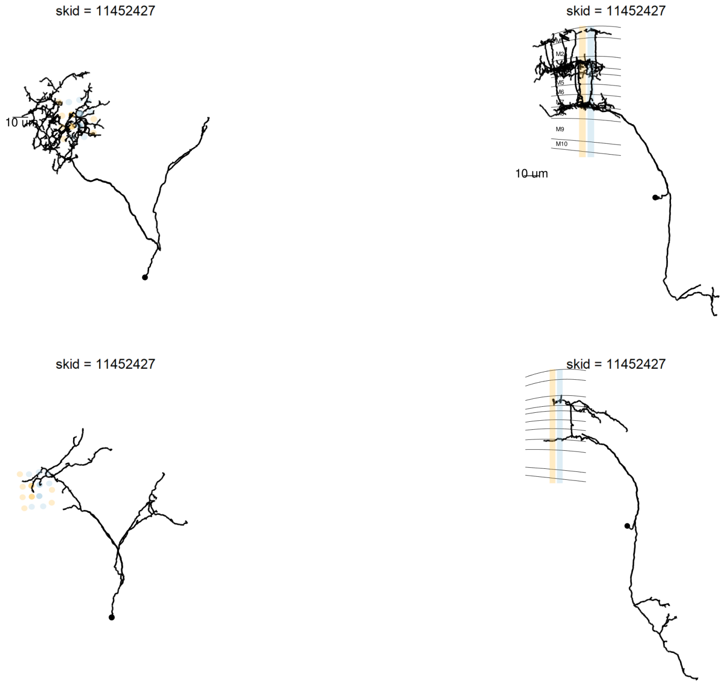

[1] 3 cells

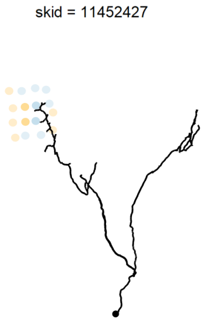

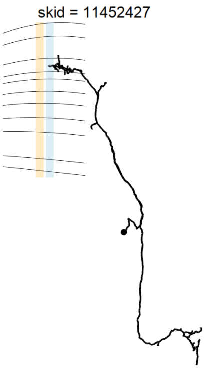

[1] 3 cells

### Central seed column C2

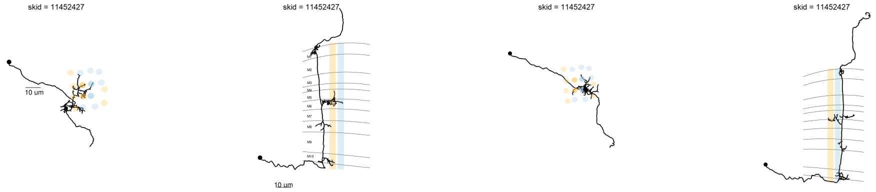

[1] 2 cells

### Central seed column Mt-VPN

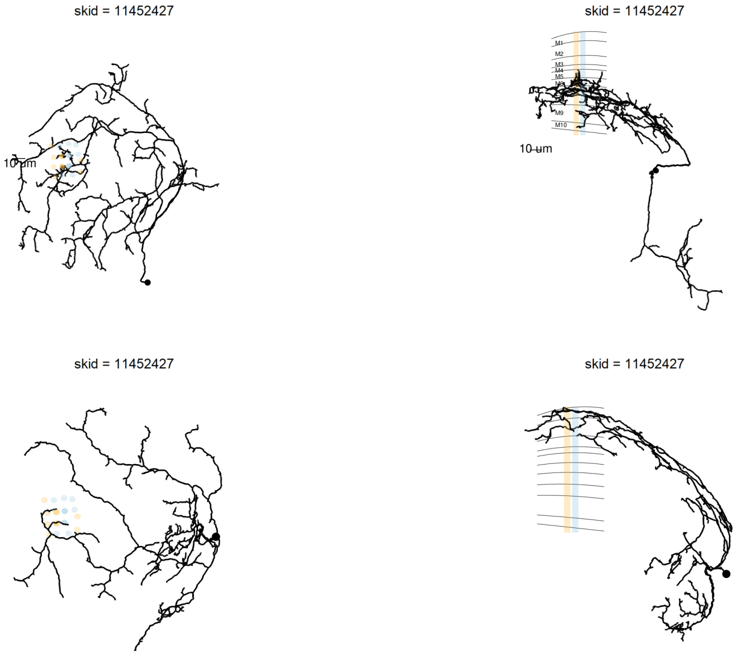

[1] 4 cells

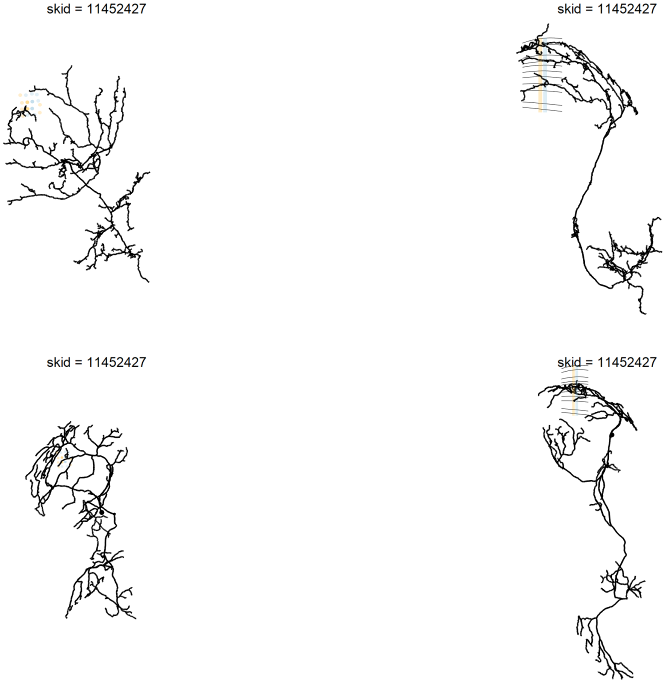

[1] 4 cells

### Central seed column Mti

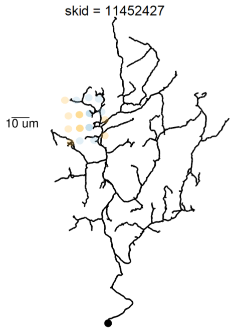

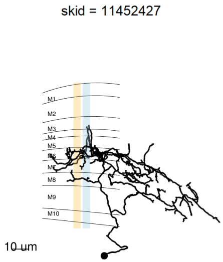

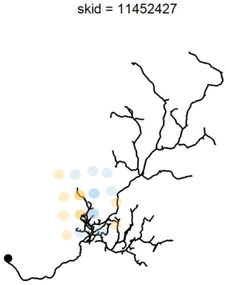

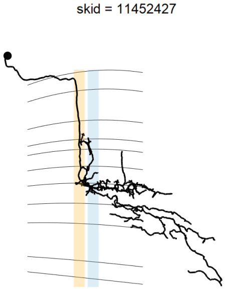

[1] 3 cells

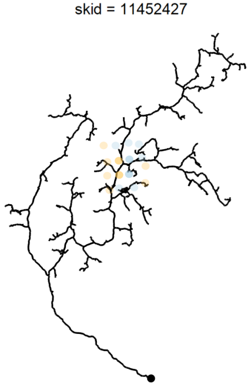

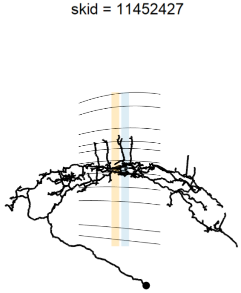

[1] 3 cells

### Central seed column Tm5a-like

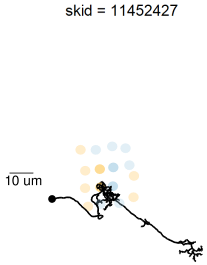

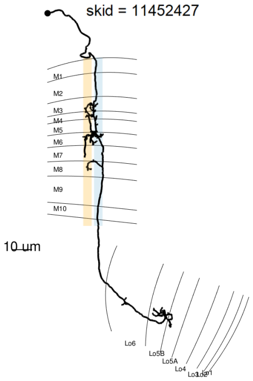

[1] 1 cells

### Central seed column TmY10

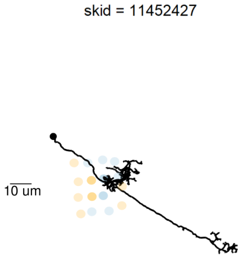

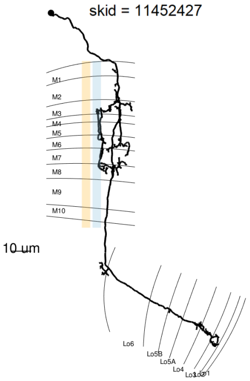

[1] 1 cells

### Central seed column Mi10

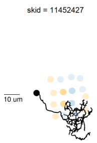

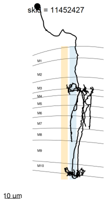

[1] 1 cells

### Central seed column Mi

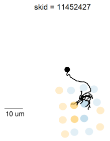

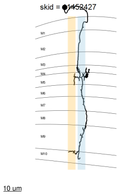

[1] 1 cells

### Central seed column C3

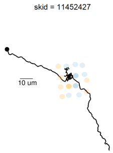

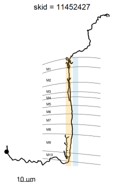

[1] 1 cells

### Central seed column Identified-<3

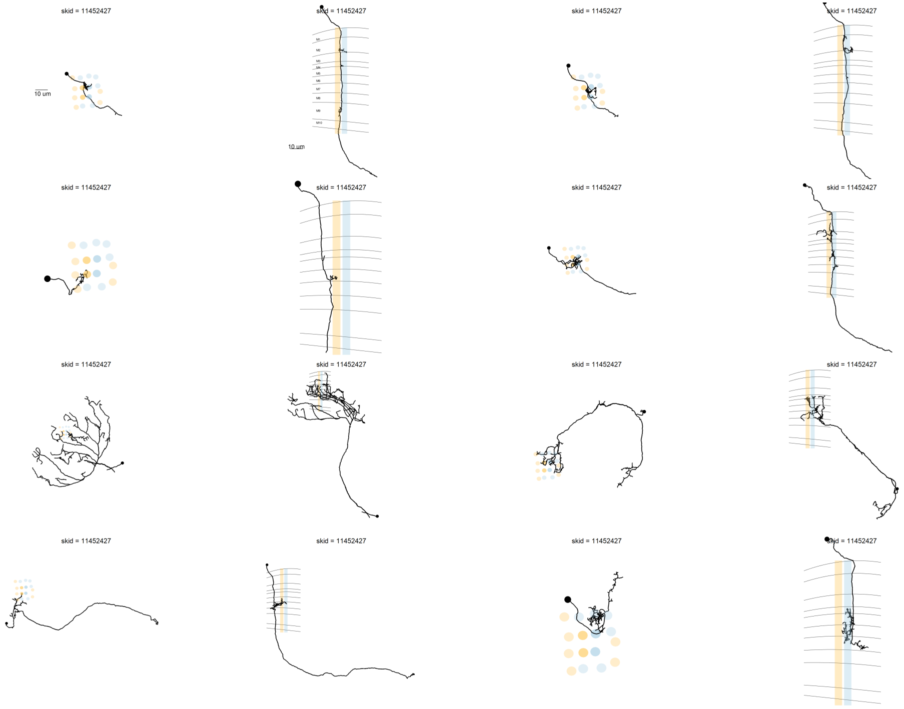

[1] 26 cells

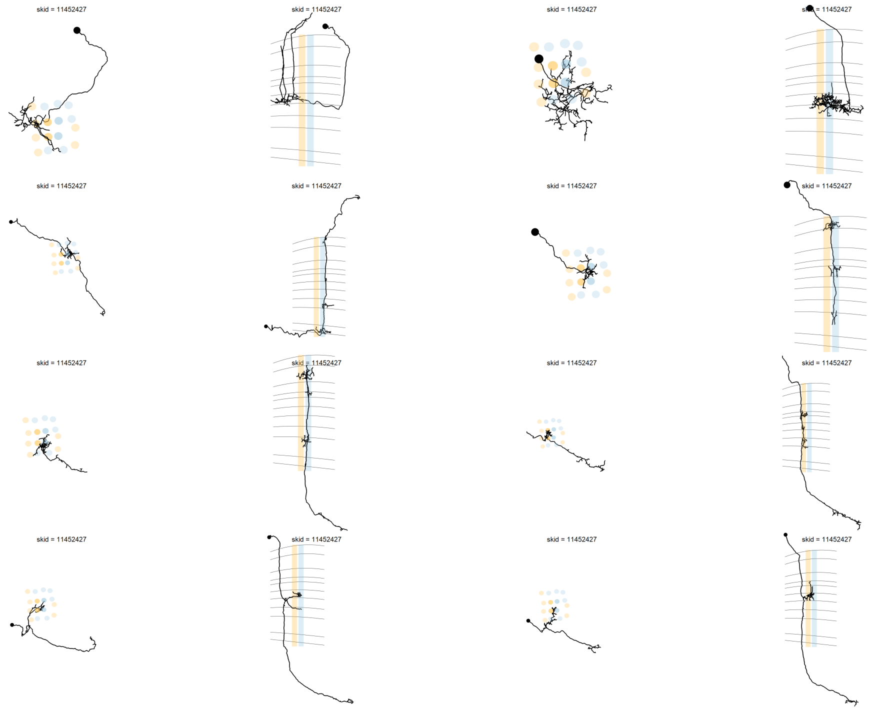
skid = 11452427
skid = 11452427
skid = 11452427
skid = 11452427
skid = 11452427
skid = 11452427
skid = 11452427
skid = 11452427
skid = 11452427
skid = 11452427
skid = 11452427
skid = 11452427
skid = 11452427
skid = 11452427
skid = 11452427
skid = 11452427

[1] 26 cells

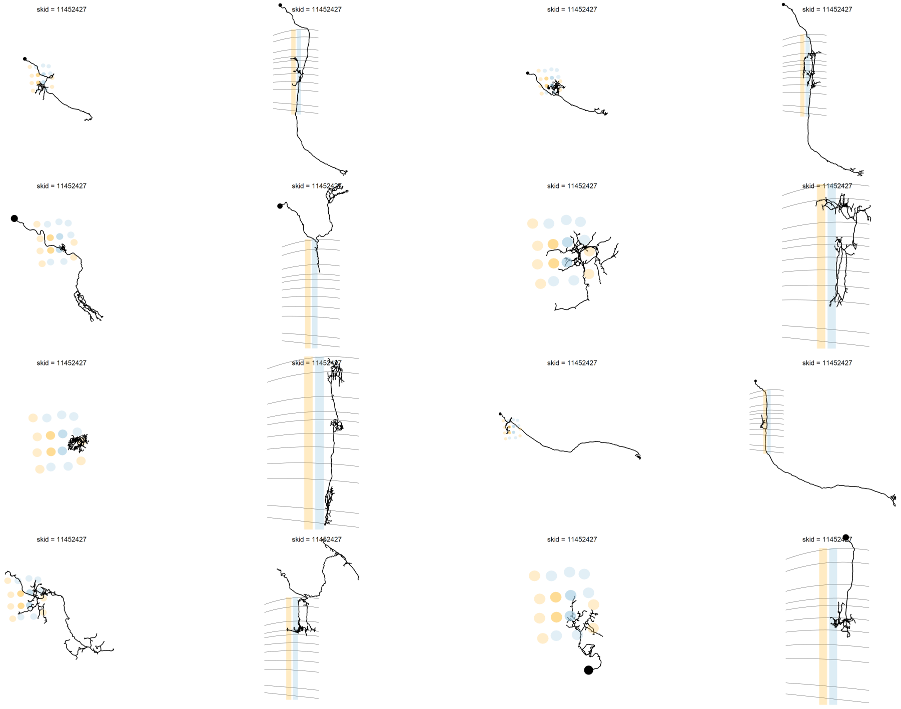
skid = 11452427
skid = 11452427
skid = 11452427
skid = 11452427
skid = 11452427
skid = 11452427
skid = 11452427
skid = 11452427
skid = 11452427
skid = 11452427
skid = 11452427
skid = 11452427
skid = 11452427
skid = 11452427
skid = 11452427
skid = 11452427

[1] 26 cells

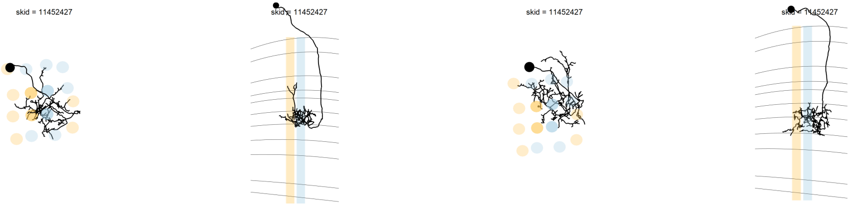

[1] 26 cells

### Central seed column Unidentified->=3

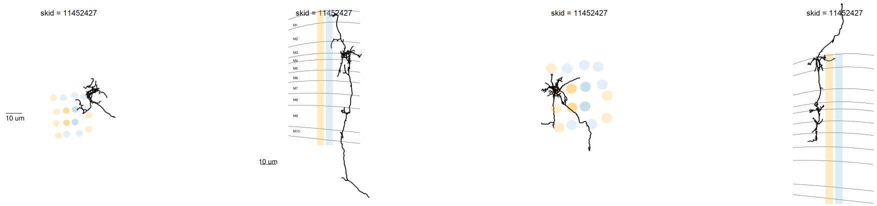

[1] 2 cells

### Central seed column Unidentified-<3

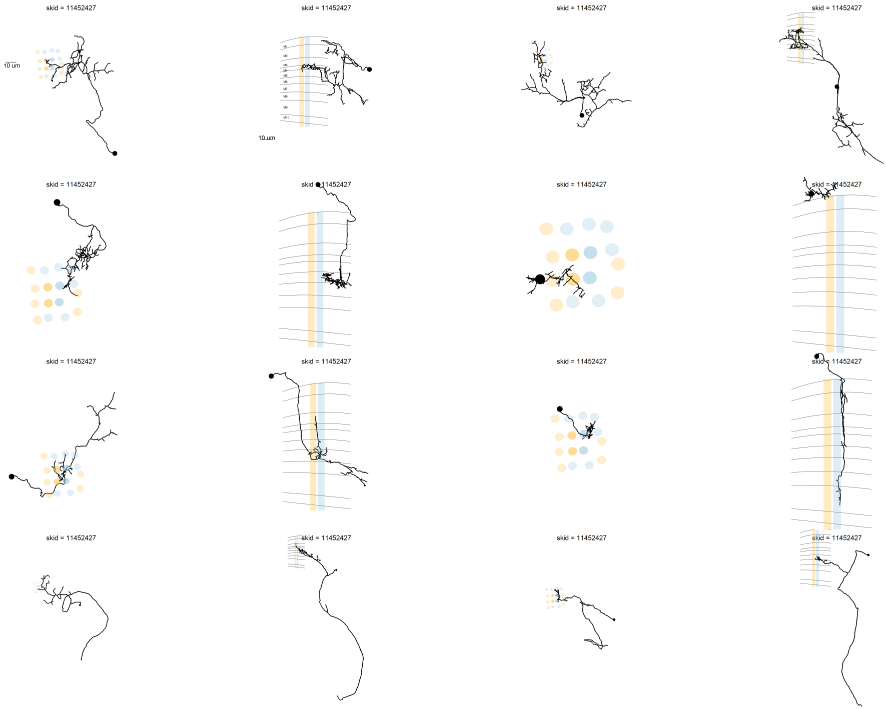

[1] 62 cells

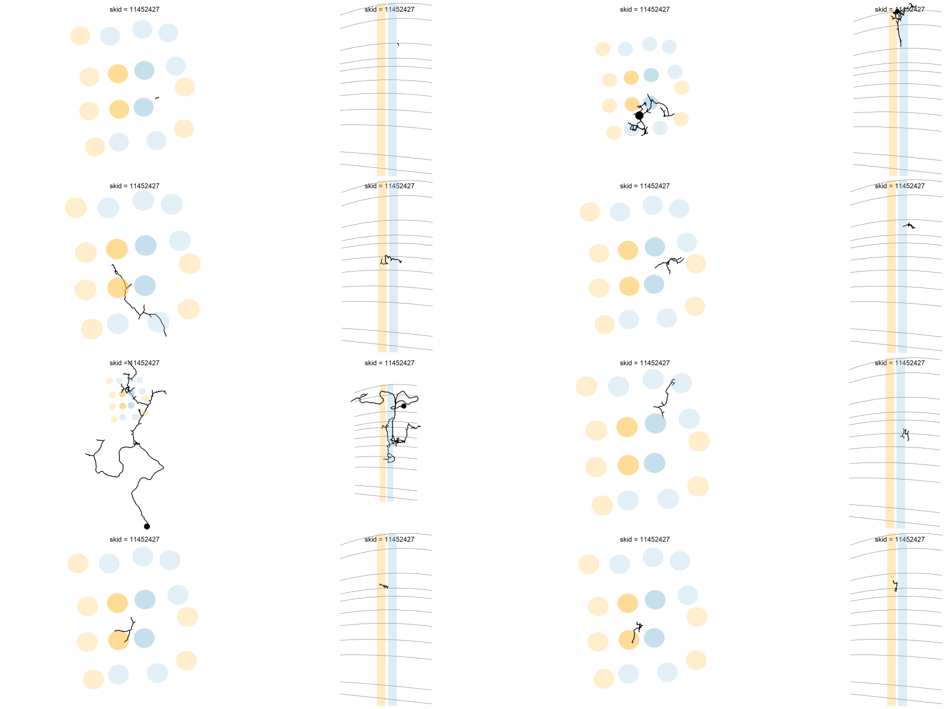
skid = 11452427
skid = 11452427
skid = 11452427
skid = 11452427
skid = 11452427
skid = 11452427
skid = 11452427
skid = 11452427
skid = 11452427
skid = 11452427
skid = 11452427
skid = 11452427
skid = 11452427
skid = 11452427
skid = 11452427
skid = 11452427

[1] 62 cells

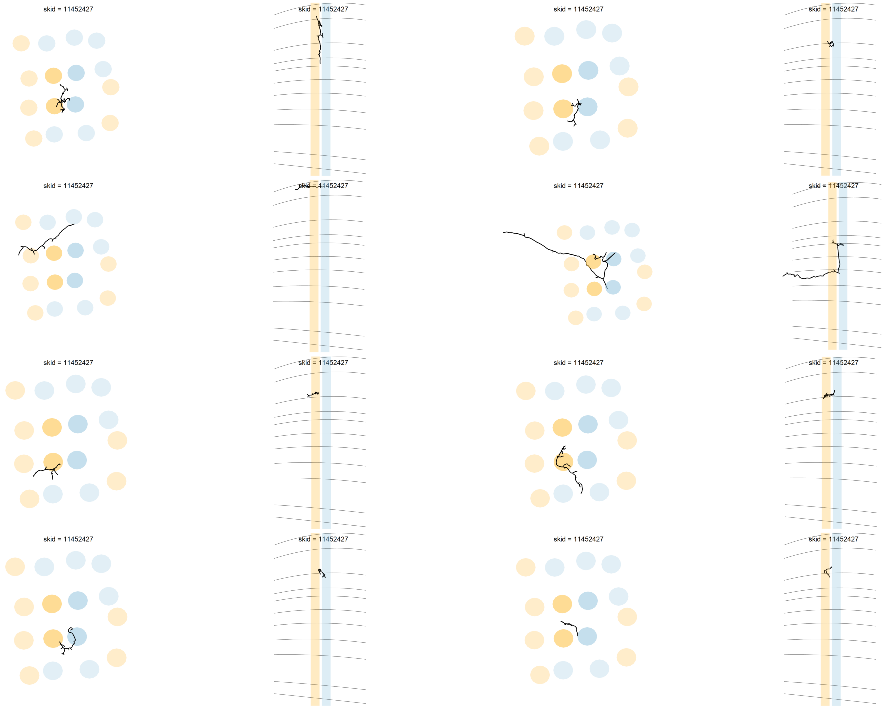
skid = 11452427
skid = 11452427
skid = 11452427
skid = 11452427
skid = 11452427
skid = 11452427
skid = 11452427
skid = 11452427
skid = 11452427
skid = 11452427
skid = 11452427
skid = 11452427
skid = 11452427
skid = 11452427
skid = 11452427
skid = 11452427

[1] 62 cells

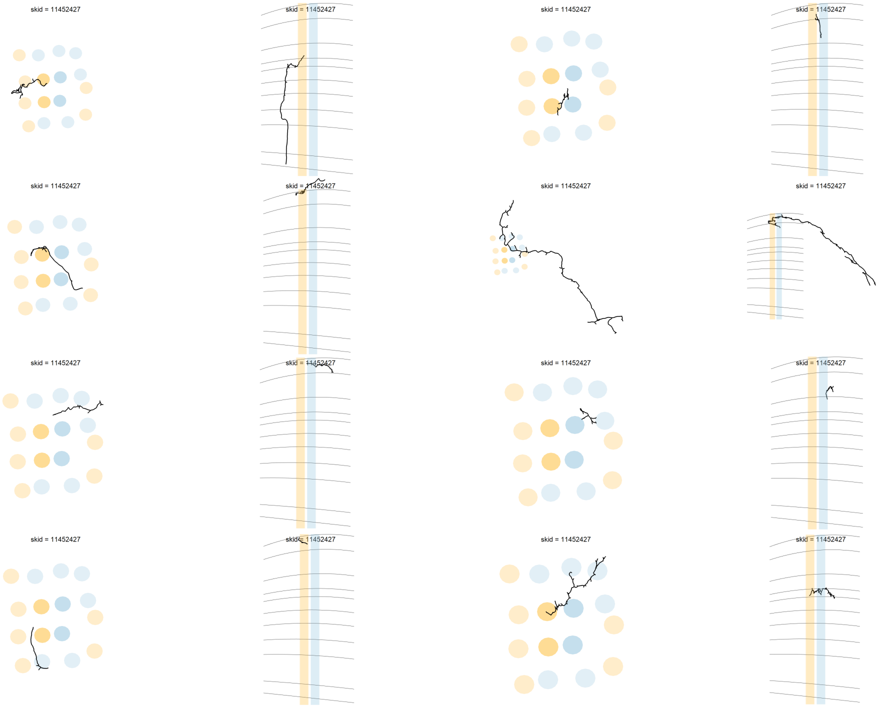
skid = 11452427
skid = 11452427
skid = 11452427
skid = 11452427
skid = 11452427
skid = 11452427
skid = 11452427
skid = 11452427
skid = 11452427
skid = 11452427
skid = 11452427
skid = 11452427
skid = 11452427
skid = 11452427
skid = 11452427
skid = 11452427

[1] 62 cells

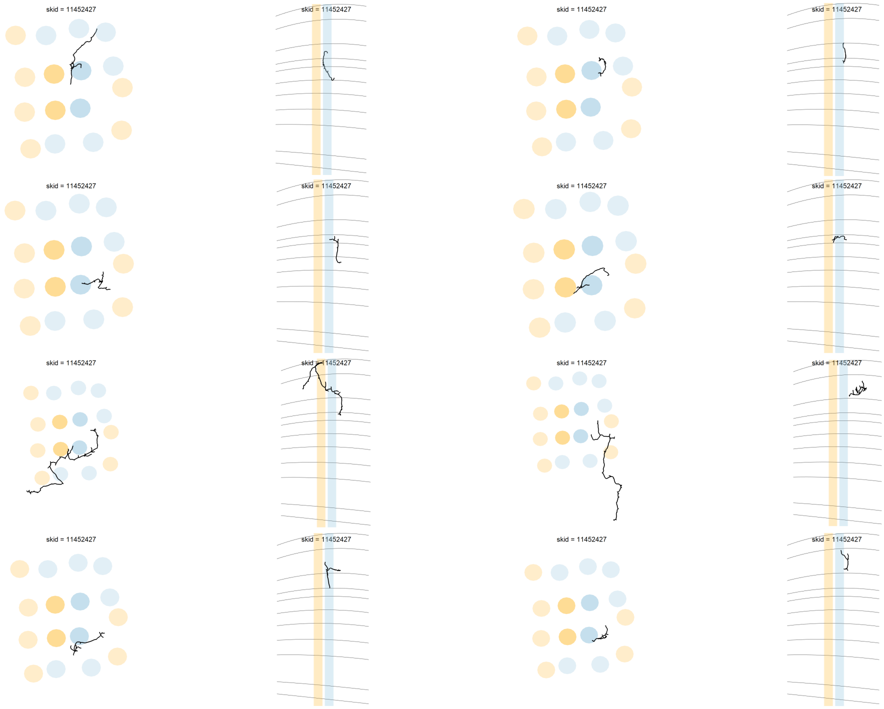
skid = 11452427
skid = 11452427
skid = 11452427
skid = 11452427
skid = 11452427
skid = 11452427
skid = 11452427
skid = 11452427
skid = 11452427
skid = 11452427
skid = 11452427
skid = 11452427
skid = 11452427
skid = 11452427
skid = 11452427
skid = 11452427

[1] 62 cells

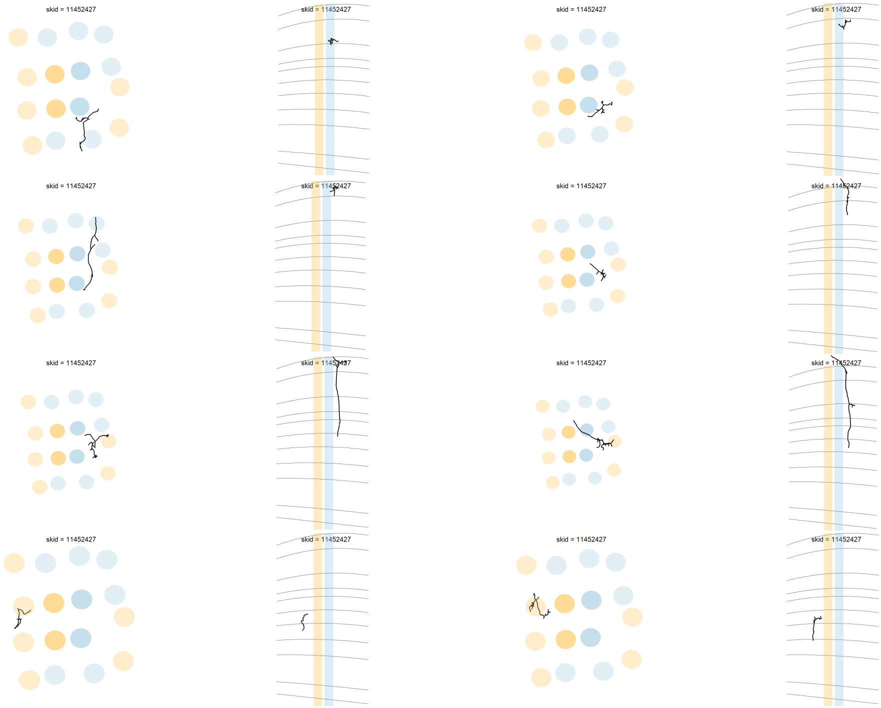
skid = 11452427
skid = 11452427
skid = 11452427
skid = 11452427
skid = 11452427
skid = 11452427
skid = 11452427
skid = 11452427
skid = 11452427
skid = 11452427
skid = 11452427
skid = 11452427
skid = 11452427
skid = 11452427
skid = 11452427
skid = 11452427

[1] 62 cells

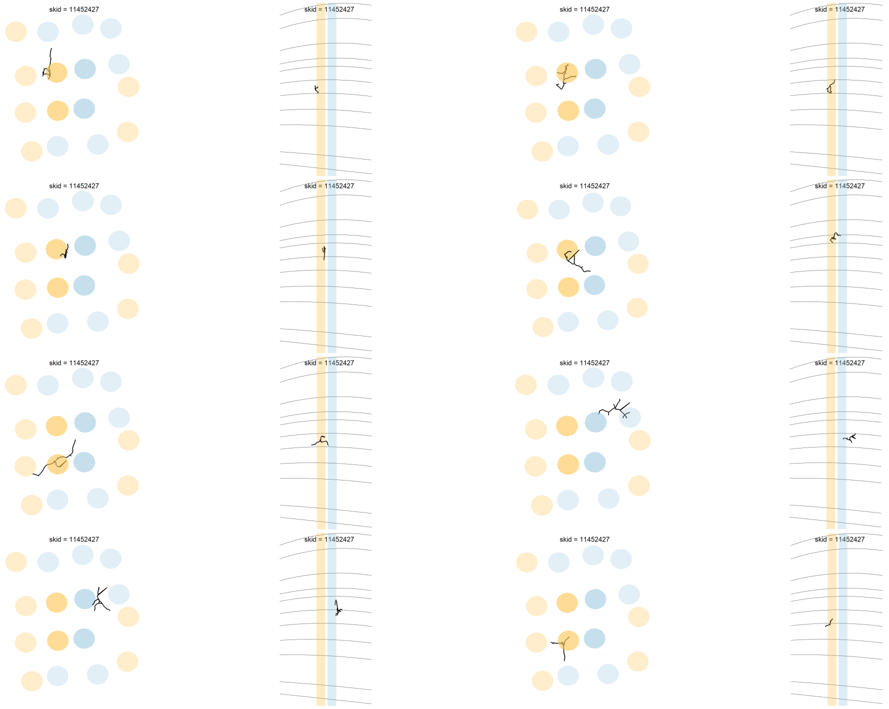
skid = 11452427
skid = 11452427
skid = 11452427
skid = 11452427
skid = 11452427
skid = 11452427
skid = 11452427
skid = 11452427
skid = 11452427
skid = 11452427
skid = 11452427
skid = 11452427
skid = 11452427
skid = 11452427
skid = 11452427
skid = 11452427

[1] 62 cells

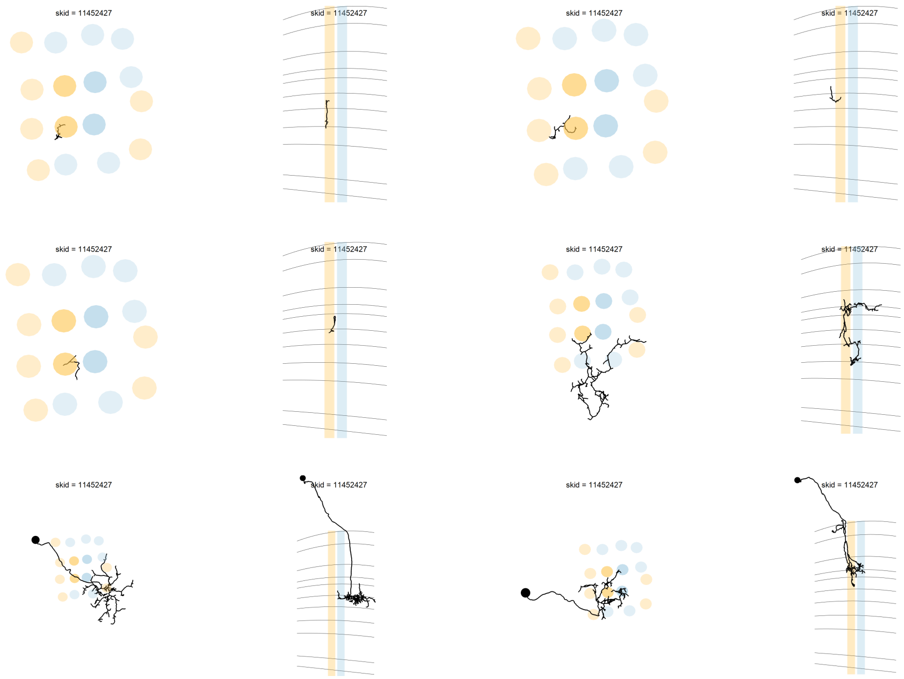
skid = 11452427
skid = 11452427
skid = 11452427
skid = 11452427
skid = 11452427
skid = 11452427
skid = 11452427
skid = 11452427
skid = 11452427
skid = 11452427
skid = 11452427
skid = 11452427

[1] 62 cells

### DRA seed column Dm-DRA1

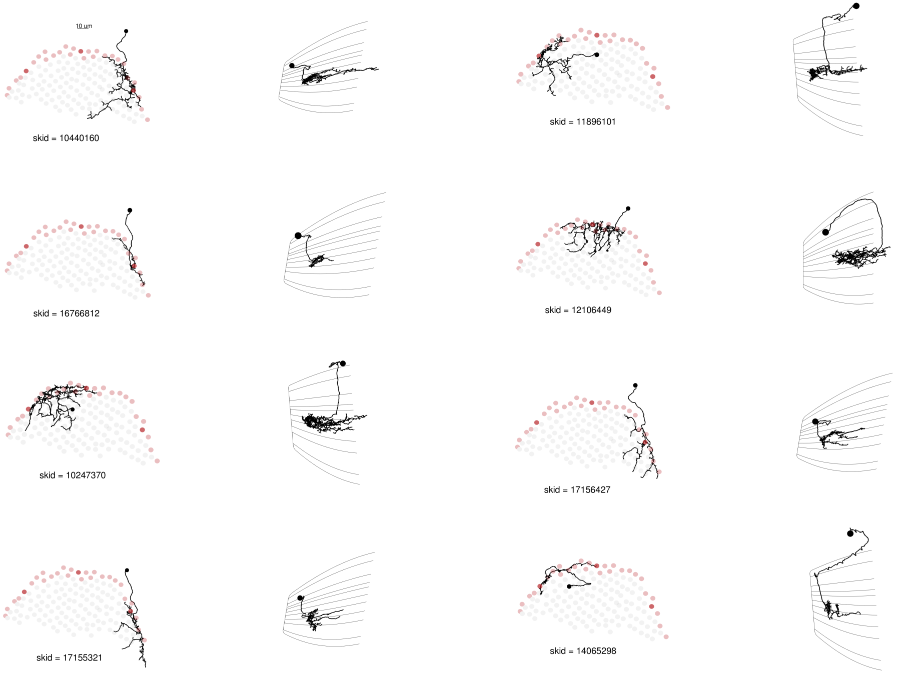

[1] 20 cells

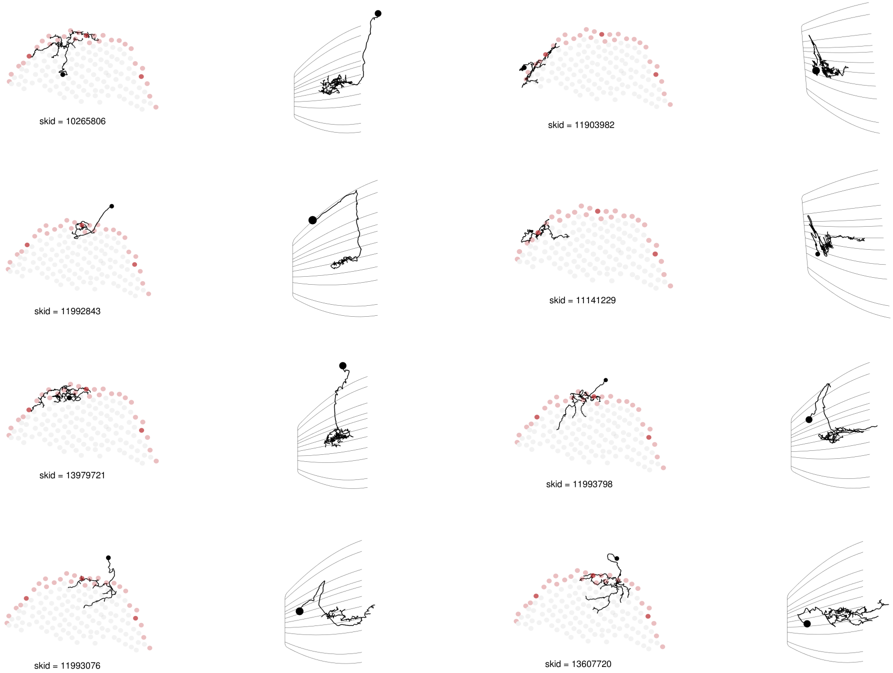
skid = 10265806
skid = 11903982
skid = 11992843
skid = 11141229
skid = 13979721
skid = 11993798
skid = 11993076
skid = 13607720

[1] 20 cells

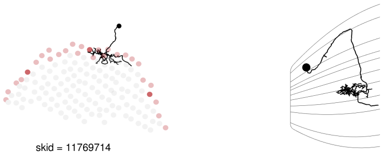

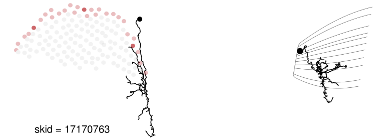

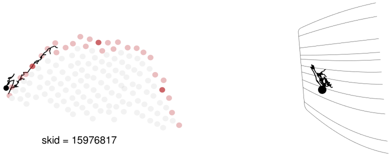

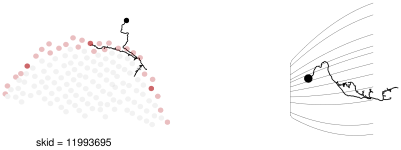

[1] 20 cells

### DRA seed column Dm9

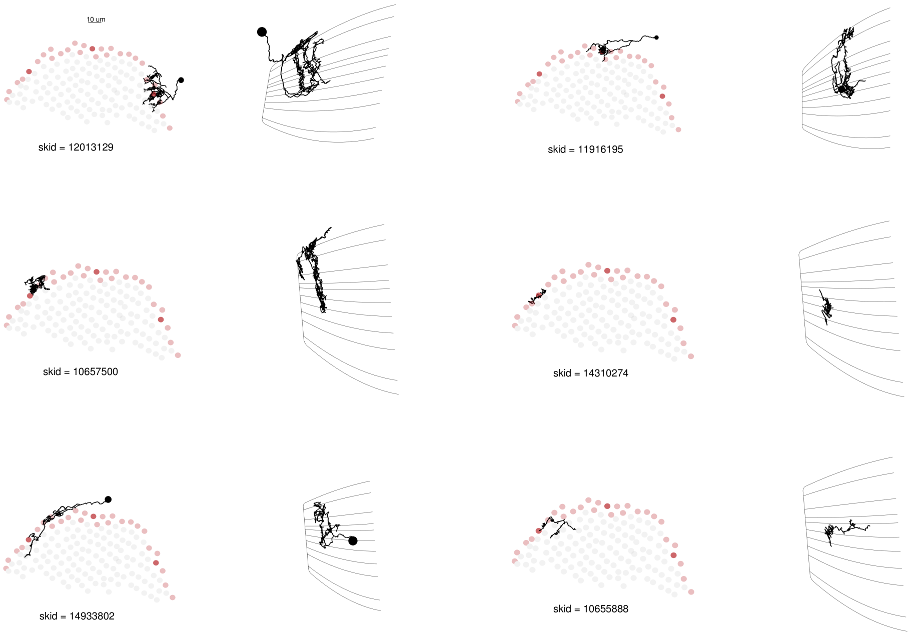

[1] 6 cells

### DRA seed column MeTu-DRA

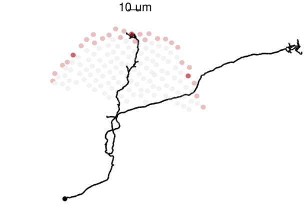

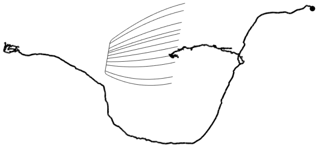

skid = 11995073

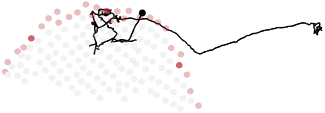

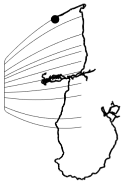

skid = 11994448

[1] 30 cells

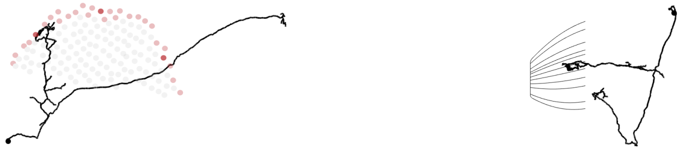

skid = 11908710

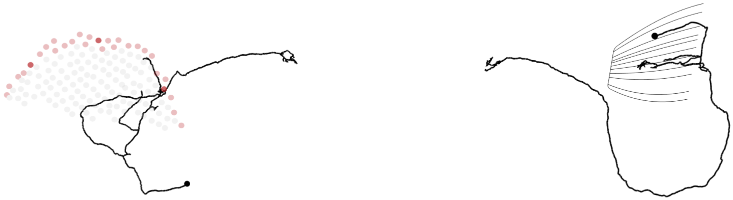

skid = 11695293

[1] 30 cells

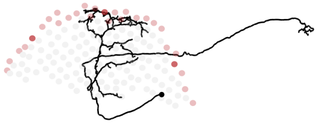

skid = 15749047

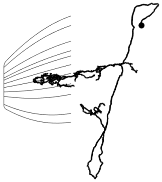

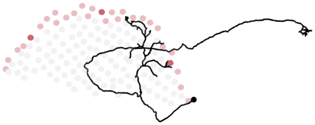

skid = 12106383

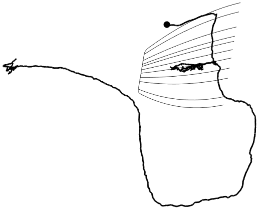

[1] 30 cells

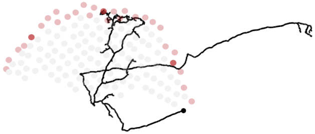

skid = 11994563

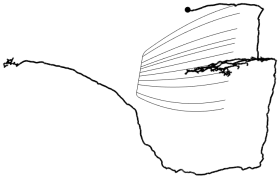

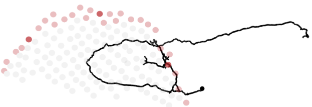

skid = 10438339

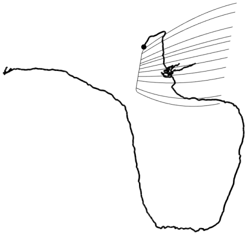

[1] 30 cells

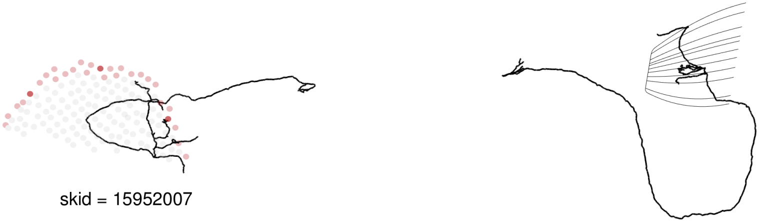

skid = 15952007

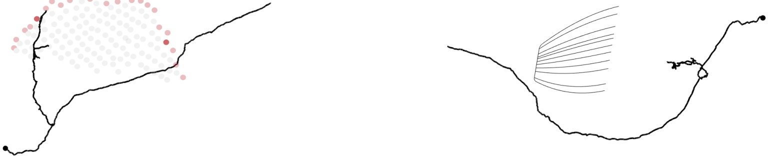

skid = 13136156

[1] 30 cells

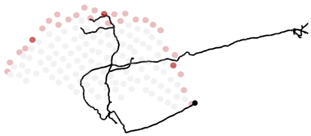

skid = 11995102

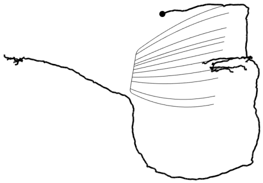

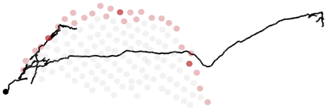

skid = 15616477

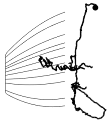

[1] 30 cells

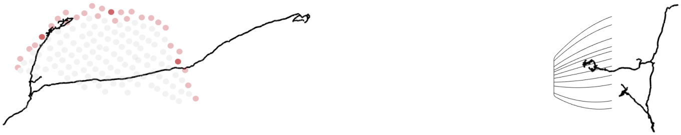

skid = 15600898

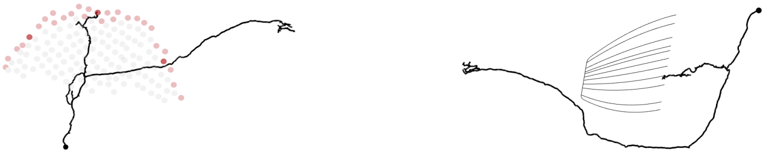

skid = 11993030

[1] 30 cells

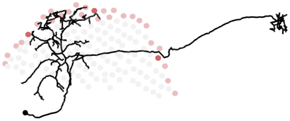

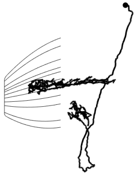

skid = 15698953

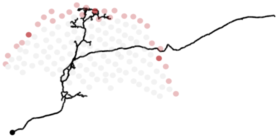

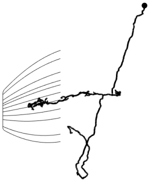

skid = 12127369

[1] 30 cells

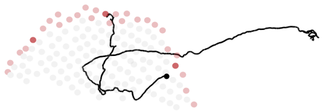

skid = 11993382

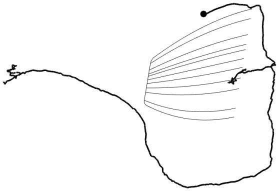

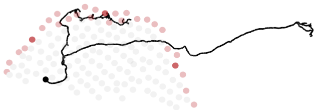

skid = 11992932

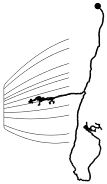

[1] 30 cells

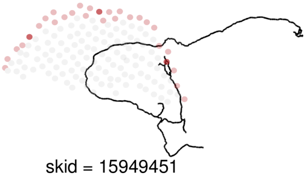

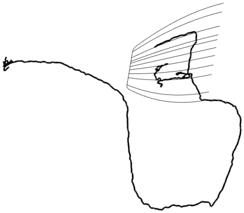

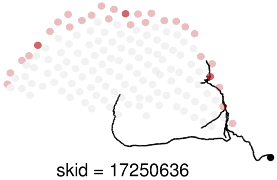

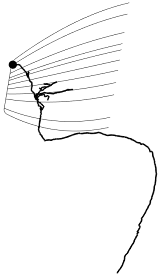

[1] 30 cells

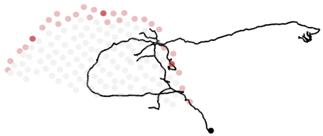

skid = 14889231

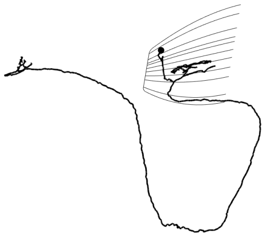

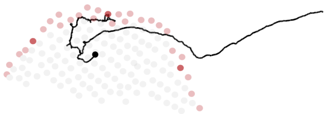

skid = 11995358

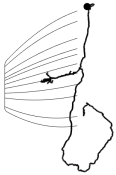

[1] 30 cells

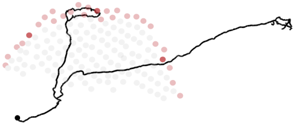

skid = 11993352

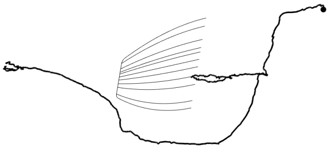

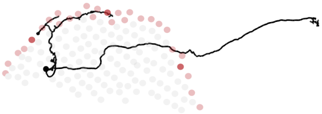

skid = 11993314

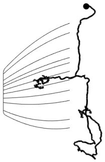

[1] 30 cells

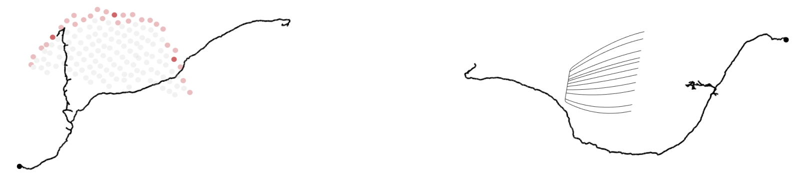

skid = 11908743

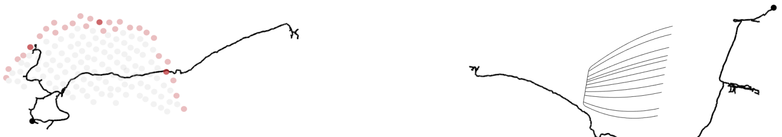

skid = 14864504

[1] 30 cells

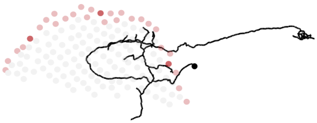

skid = 14936634

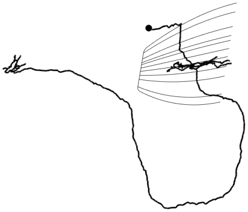

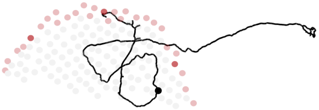

skid = 11993064

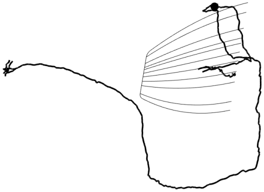

[1] 30 cells

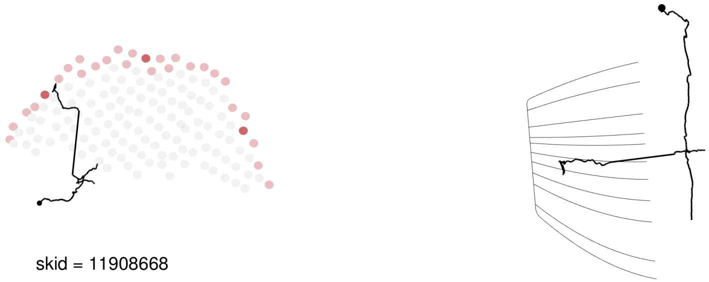

skid = 11908668

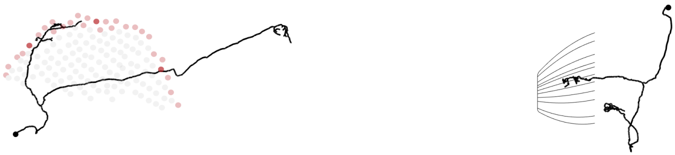

skid = 11903992

[1] 30 cells

### DRA seed column R7-DRA

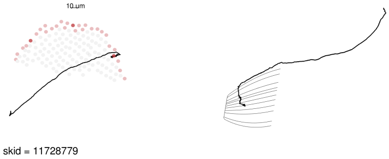

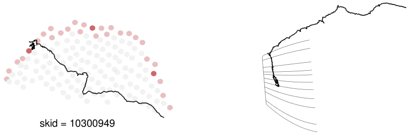

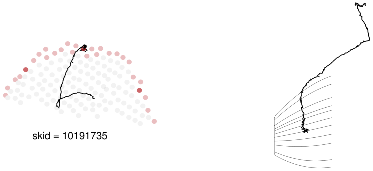

[1] 3 cells

### DRA seed column Dm-DRA2

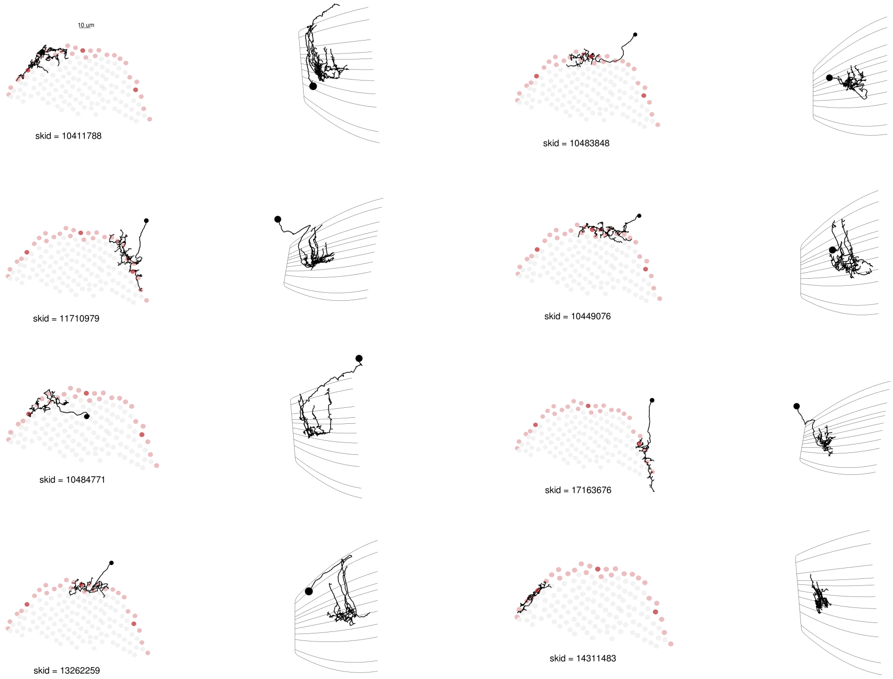

[1] 9 cells

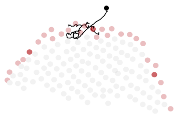

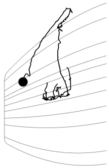

[1] 9 cells

### DRA seed column Dm2

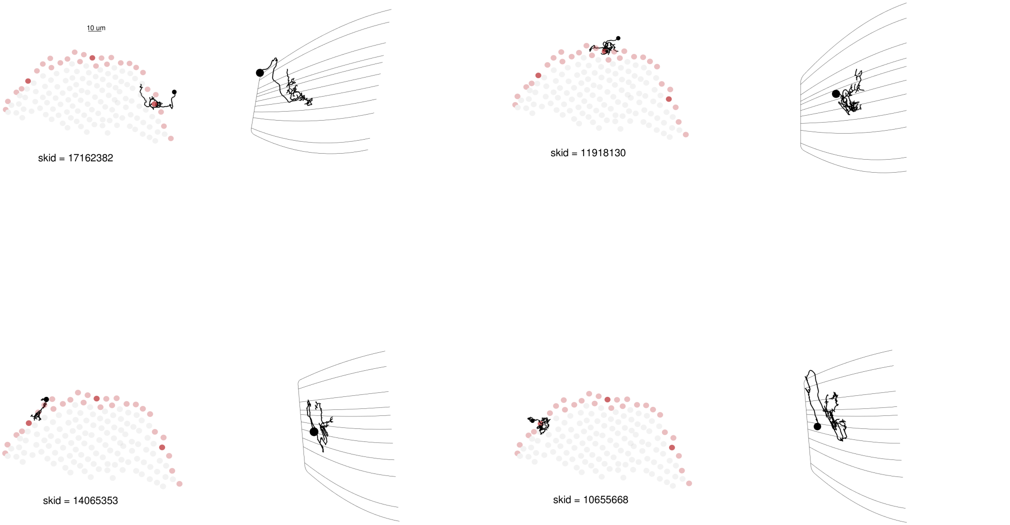

[1] 4 cells

### DRA seed column R8-DRA

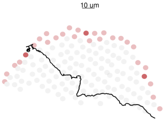

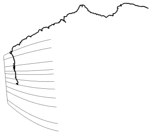

skid = 10300963

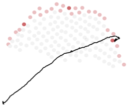
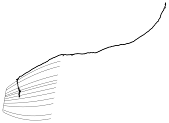

skid = 11728827

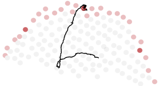
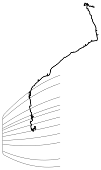

skid = 10190508

[1] 3 cells

### DRA seed column Mi15

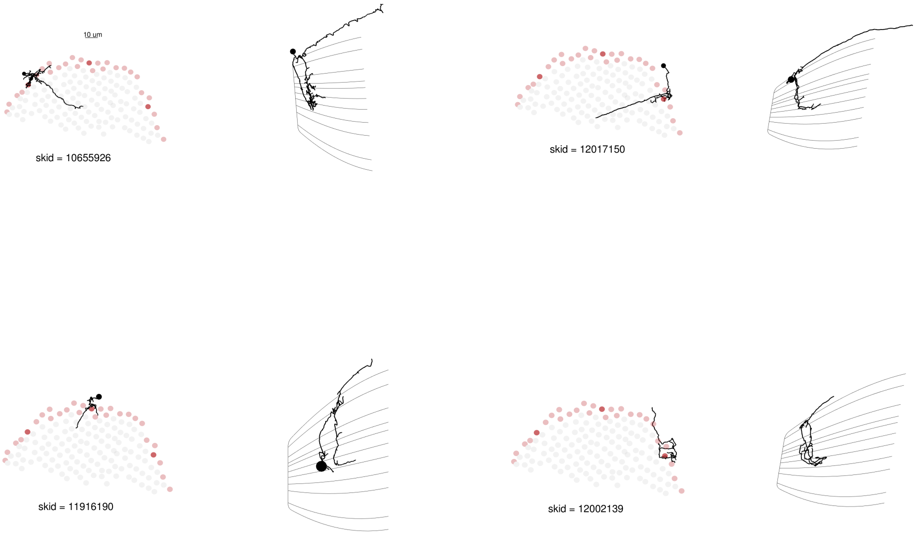

[1] 4 cells

### DRA seed column Mti-DRA-1

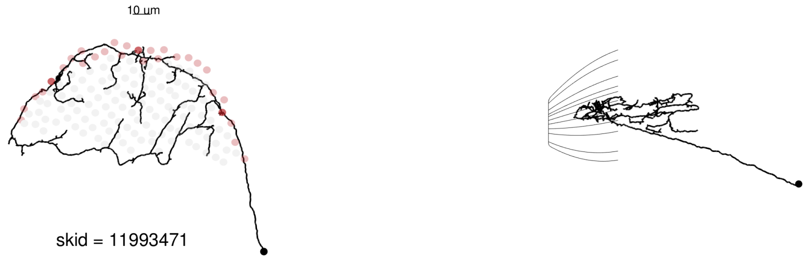

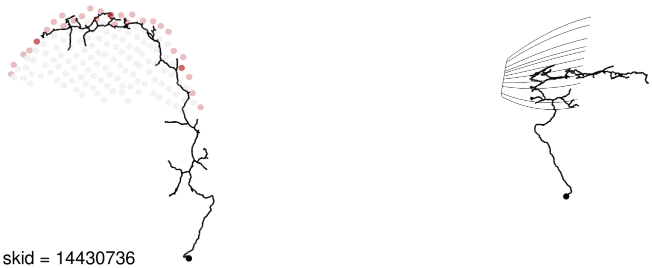

[1] 6 cells

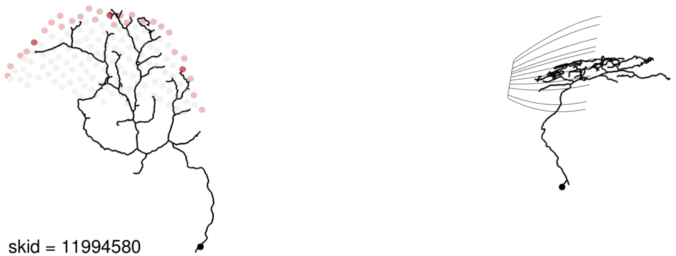

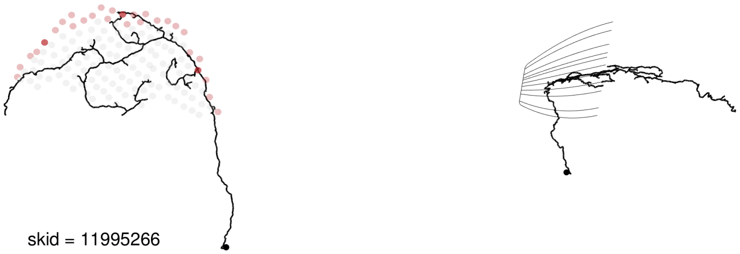

[1] 6 cells

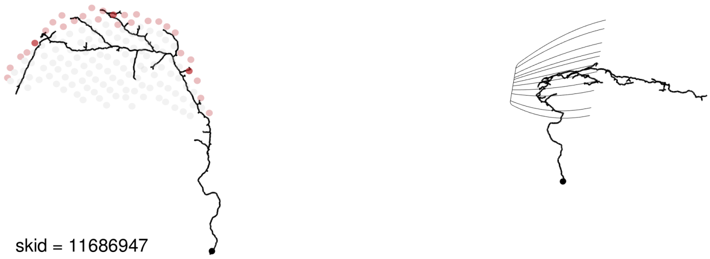

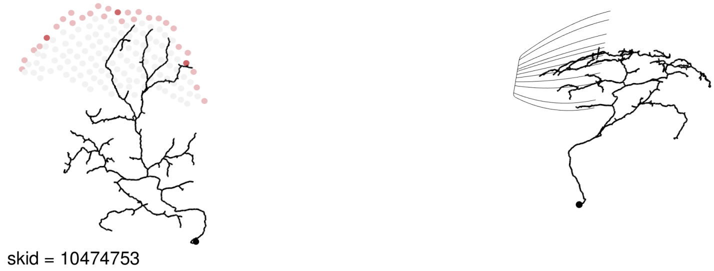

[1] 6 cells

### DRA seed column MeMe-DRA

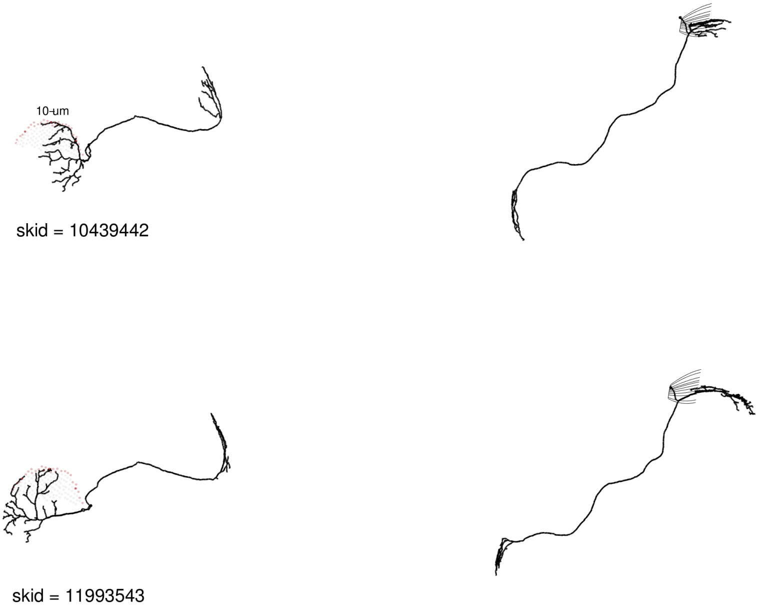

[1] 2 cells

### DRA seed column L3

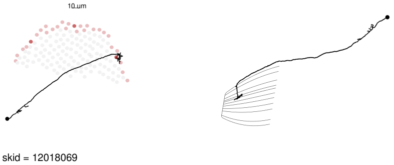

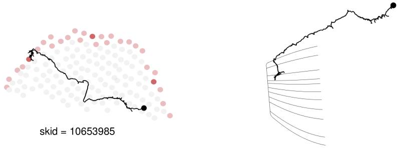

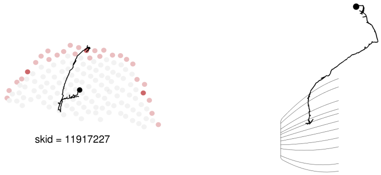

[1] 3 cells

### DRA seed column VPN-DRA

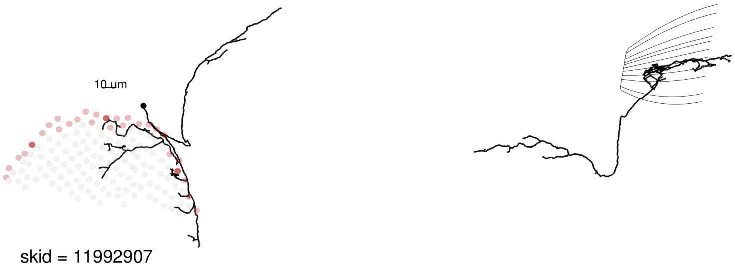

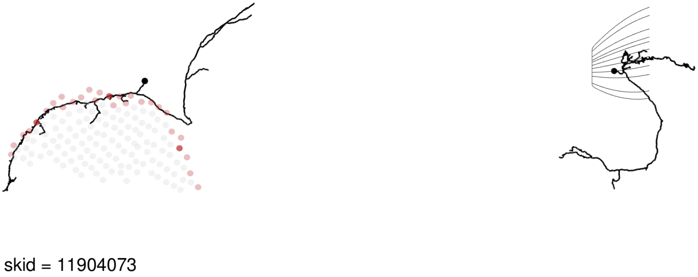

[1] 6 cells

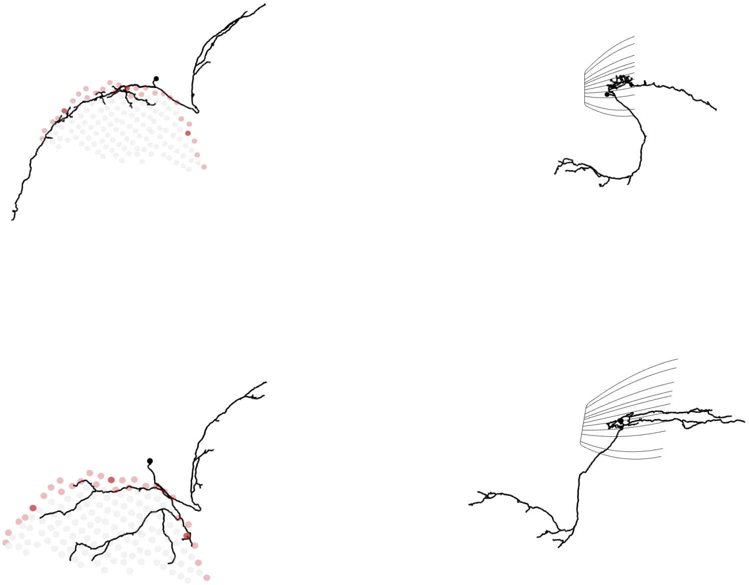

skid = 17165715

[1] 6 cells

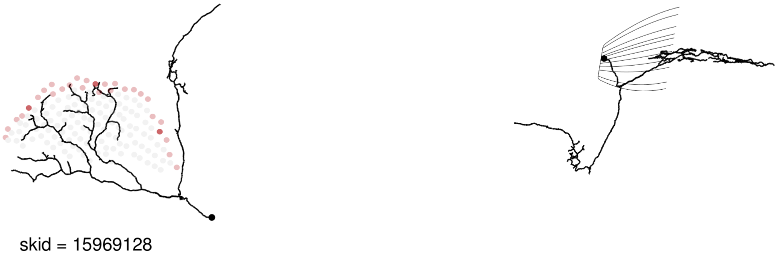

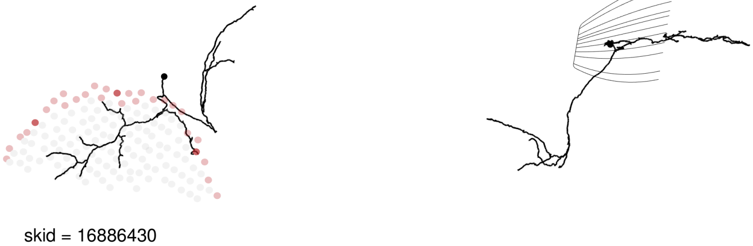

[1] 6 cells

### DRA seed column L1

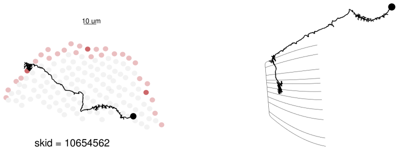

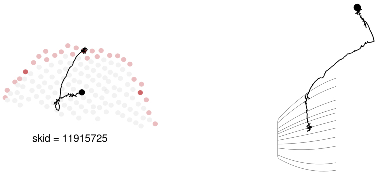

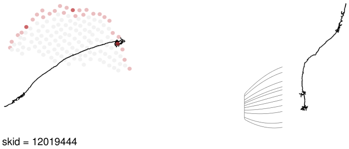

[1] 3 cells

### DRA seed column Tm20

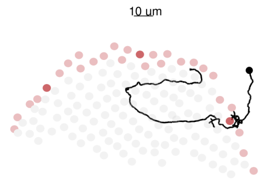

skid = 12018769

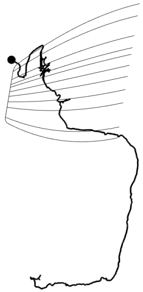

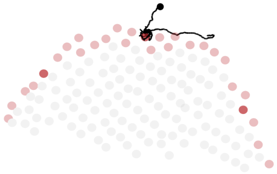

skid = 11918619

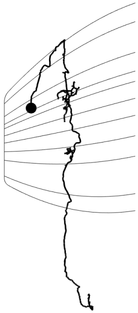

[1] 3 cells

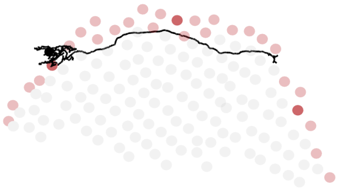

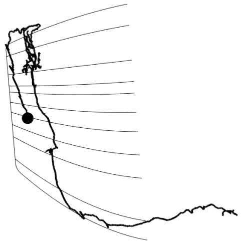

[1] 3 cells

### DRA seed column Mti-DRA-2

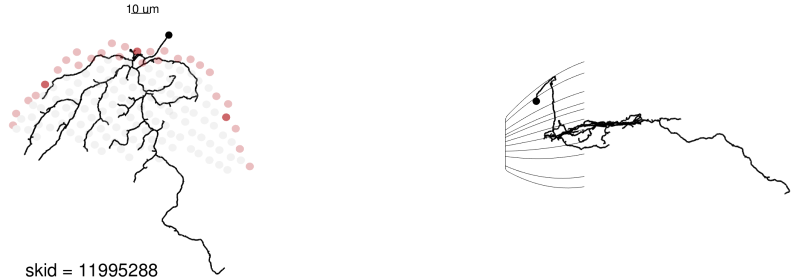

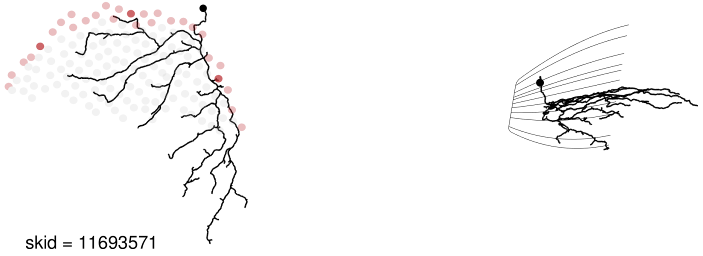

[1] 4 cells

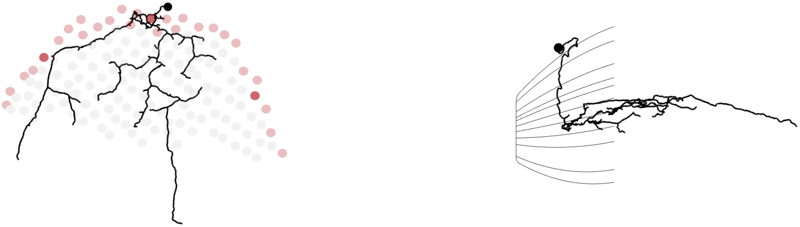

skid = 11993444

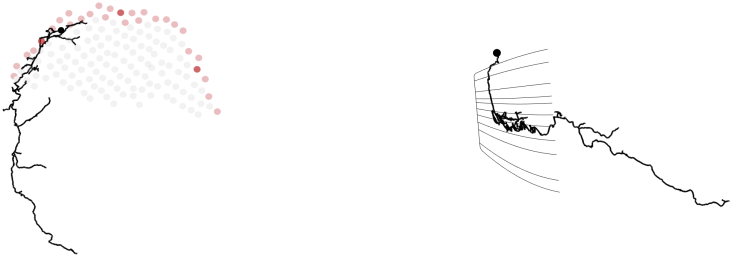

skid = 11903802

[1] 4 cells

### DRA seed column Mi1

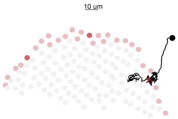

skid = 11829850

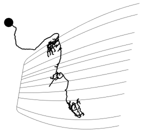

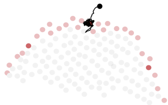

skid = 13294451

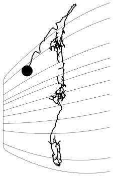

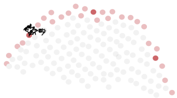

skid = 14811021

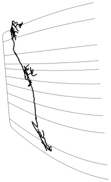

[1] 3 cells

### DRA seed column MeTu

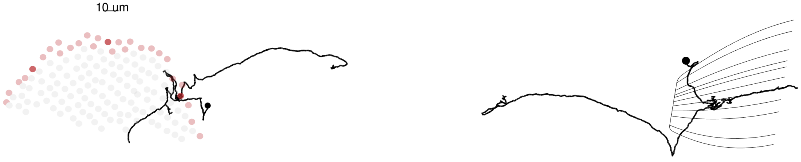

skid = 16730682

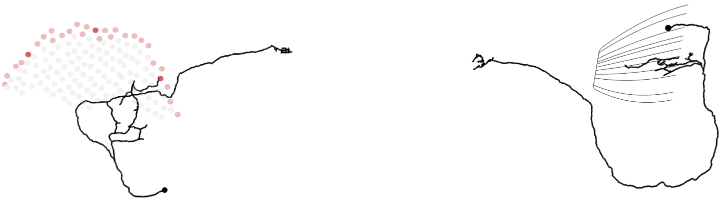

skid = 14888320

[1] 2 cells

### DRA seed column Tm5-like

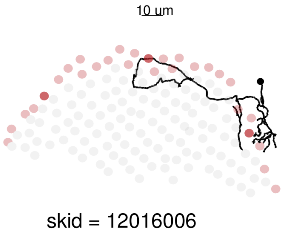

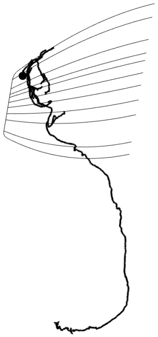

[1] 1 cells

### DRA seed column Mi9

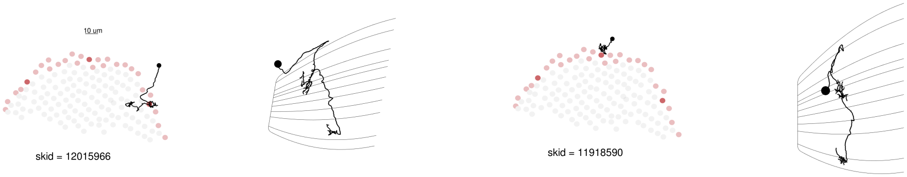

[1] 2 cells

### DRA seed column Dm11

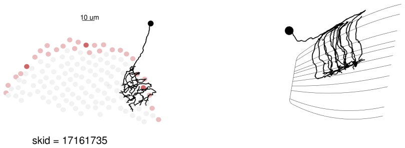

[1] 1 cells

### DRA seed column aMe12

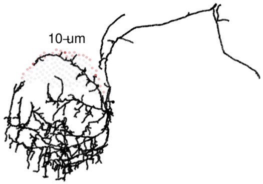

skid = 164544

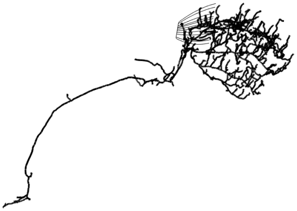

[1] 1 cells

### DRA seed column TmY

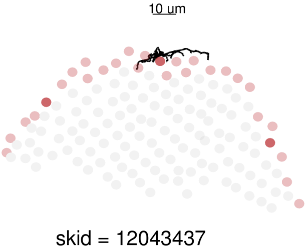

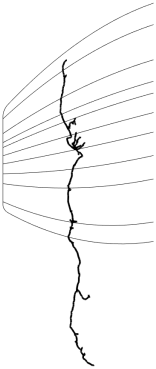

[1] 1 cells

### DRA seed column ML-VPN2

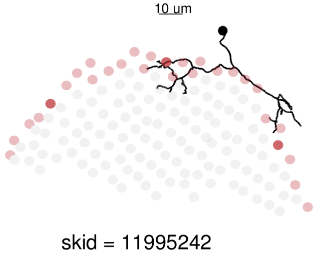

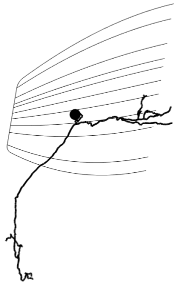

[1] 1 cells

#### DRA seed column C2

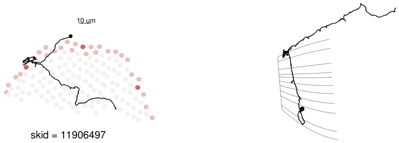

[1] 1 cells

### DRA seed column Identified-<3

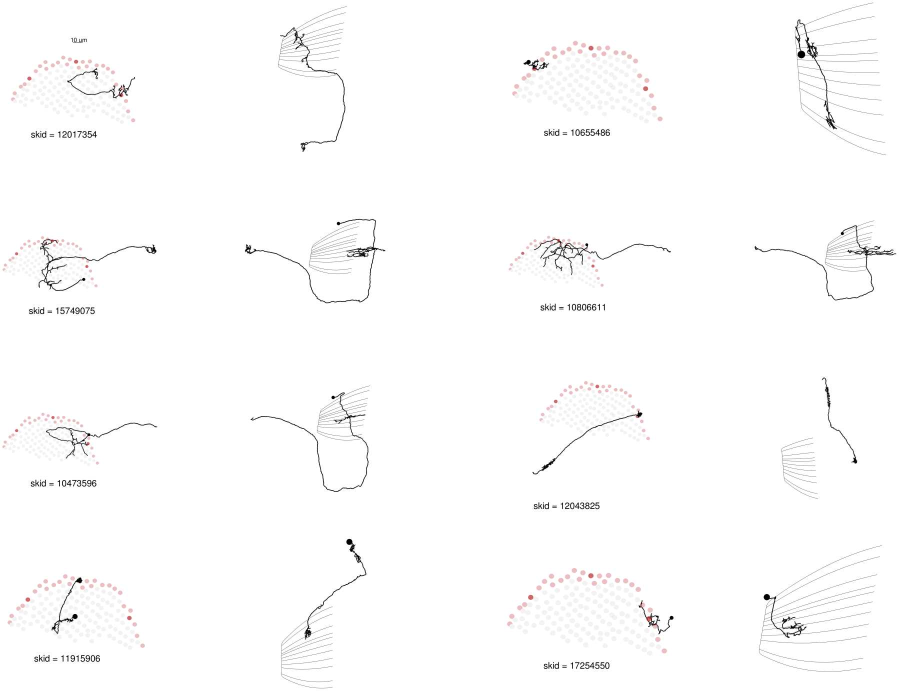

[1] 33 cells

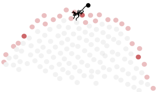
skid = 14305318
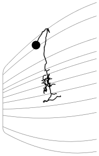
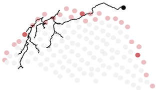
skid = 16110049
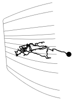
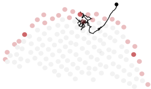
skid = 11993803
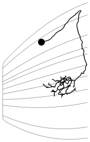
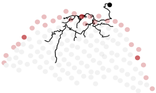
skid = 10479097
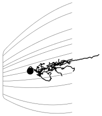
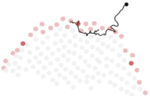
skid = 11981475
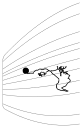
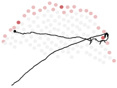
skid = 12044212
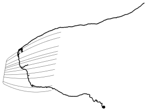
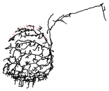
skid = 28841
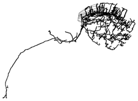
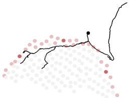
skid = 15997281
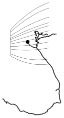

[1] 33 cells

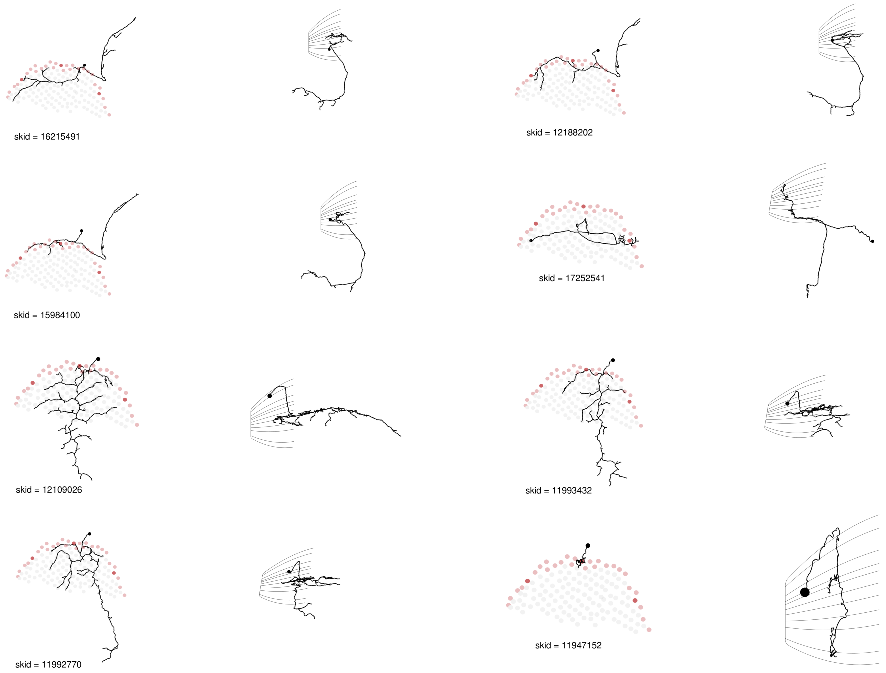
skid = 16215491
skid = 12188202
skid = 15984100
skid = 17252541
skid = 12109026
skid = 11993432
skid = 11992770
skid = 11947152

[1] 33 cells

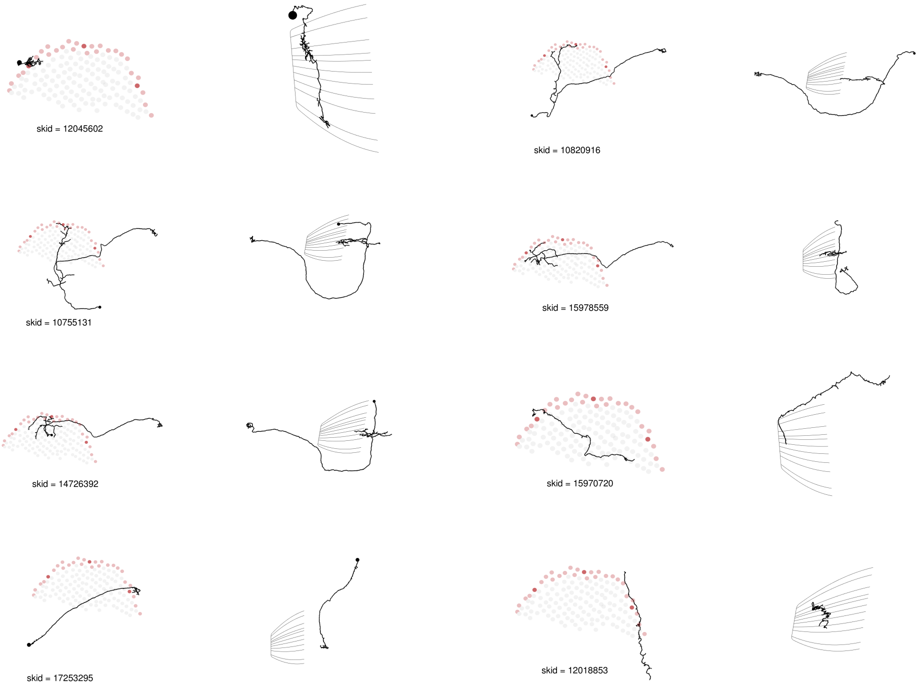
skid = 12045602
skid = 10820916
skid = 10755131
skid = 15978559
skid = 14726392
skid = 15970720
skid = 17253295
skid = 12018853

[1] 33 cells

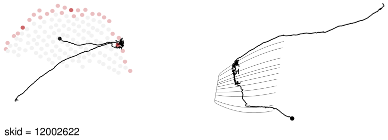

[1] 33 cells

### DRA seed column Unidentified->=3

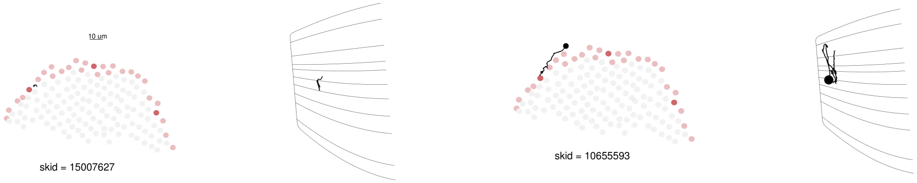

[1] 2 cells

### DRA seed column Unidentified-<3

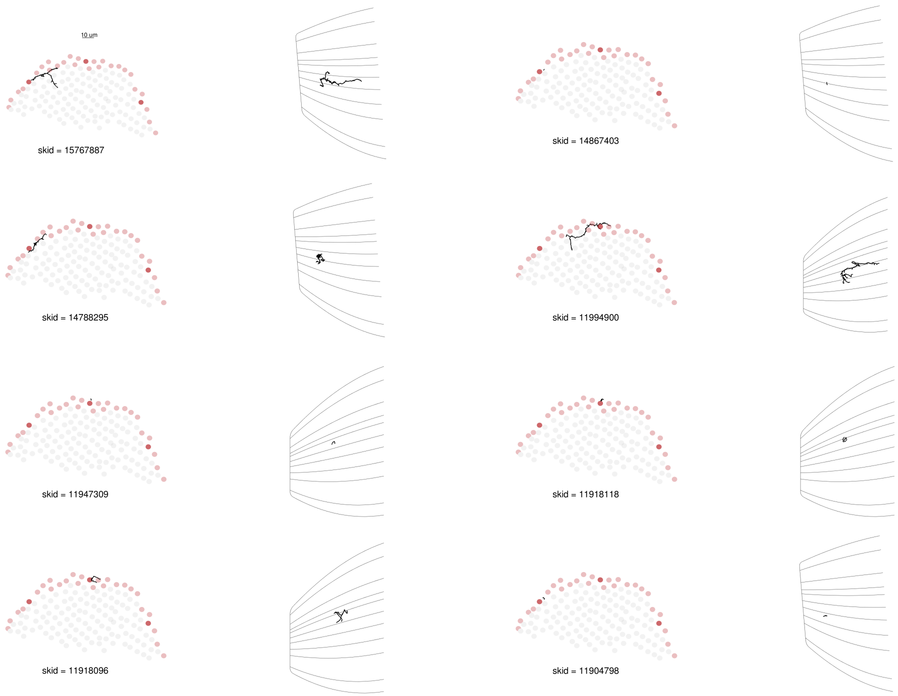

[1] 57 cells

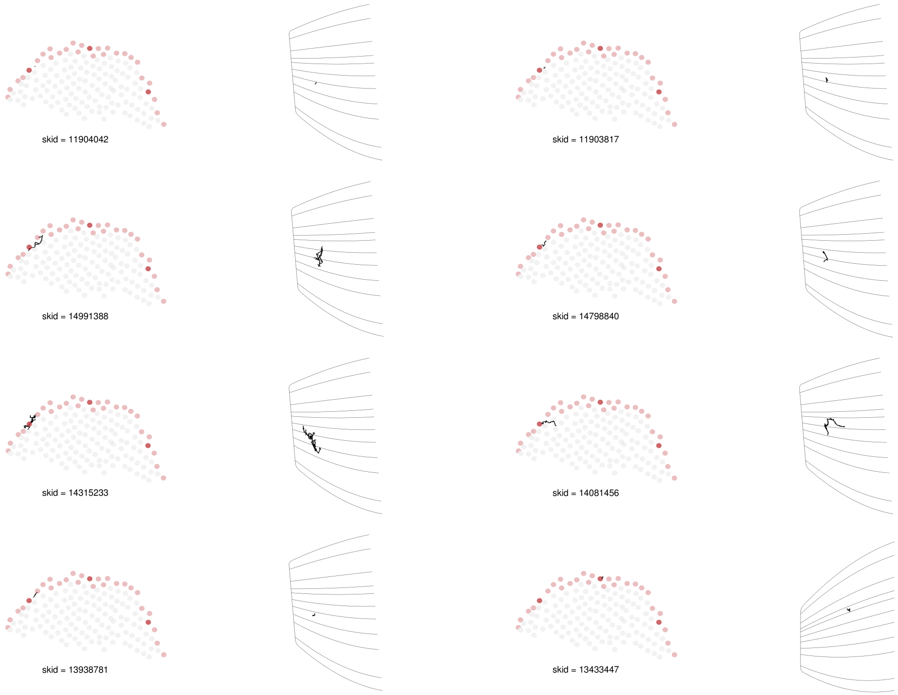
skid = 11904042
skid = 11903817
skid = 14991388
skid = 14798840
skid = 14315233
skid = 14081456
skid = 13938781
skid = 13433447

[1] 57 cells

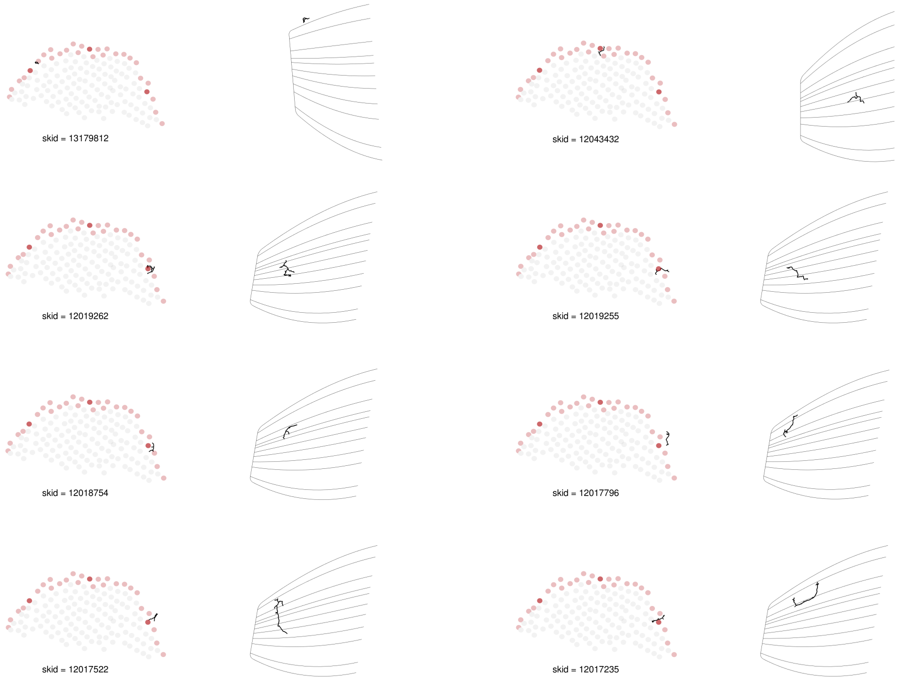
skid = 13179812
skid = 12043432
skid = 12019262
skid = 12019255
skid = 12018754
skid = 12017796
skid = 12017522
skid = 12017235

[1] 57 cells

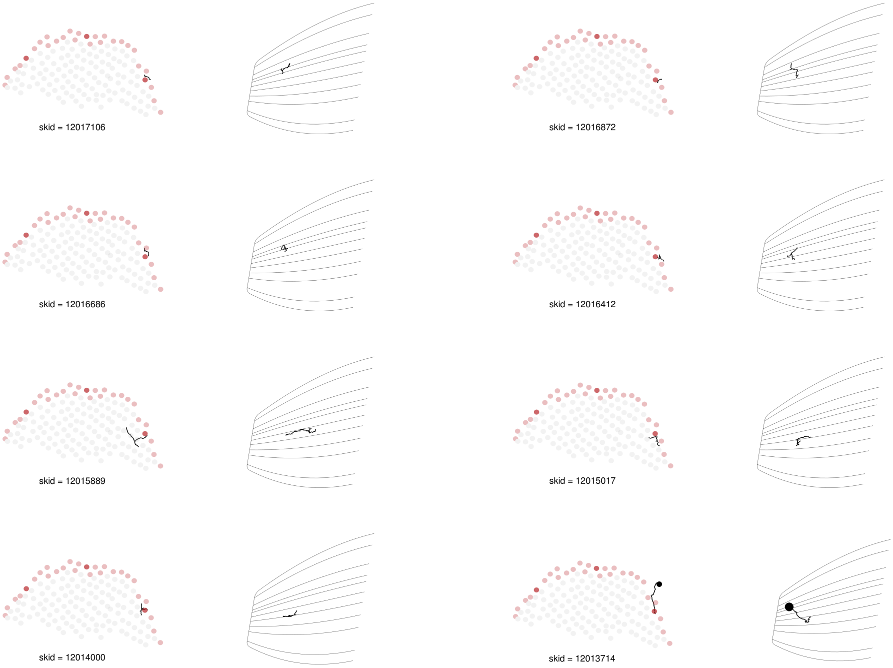
skid = 12017106
skid = 12016872
skid = 12016686
skid = 12016412
skid = 12015889
skid = 12015017
skid = 12014000
skid = 12013714

[1] 57 cells

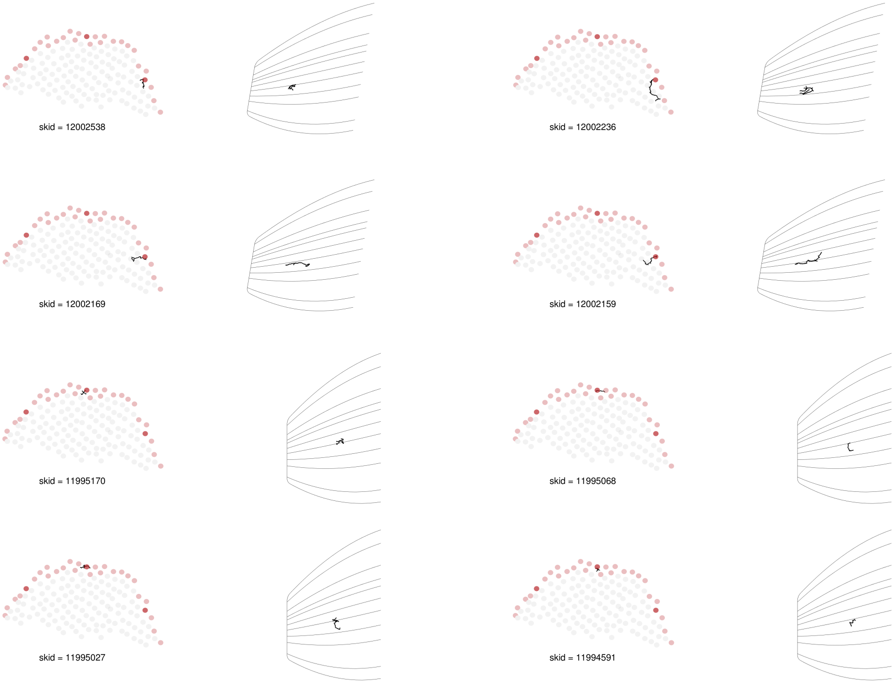
skid = 12002538
skid = 12002236
skid = 12002169
skid = 12002159
skid = 11995170
skid = 11995068
skid = 11995027
skid = 11994591

[1] 57 cells

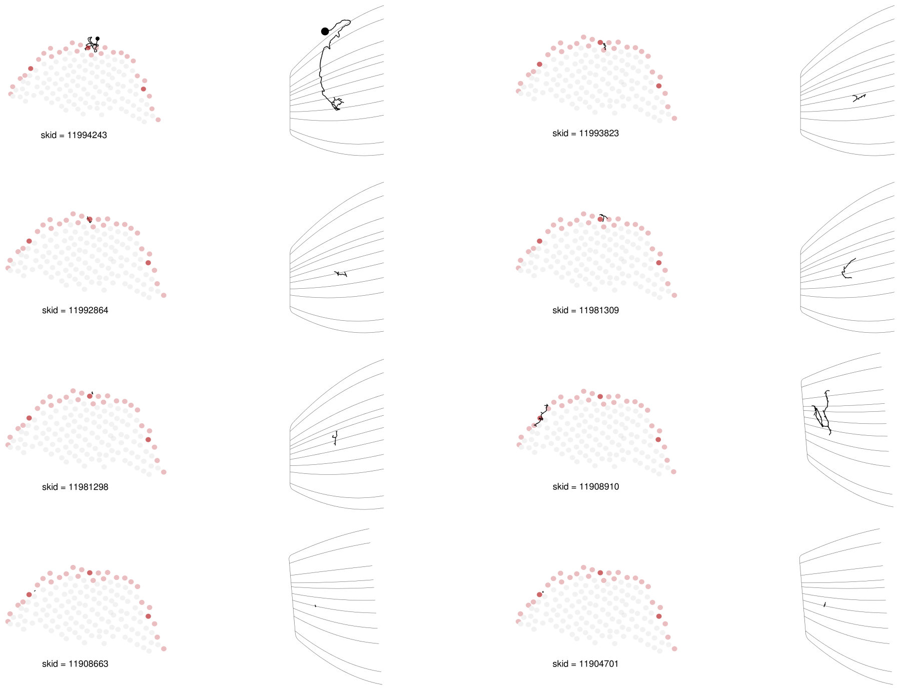
skid = 11994243
skid = 11993823
skid = 11992864
skid = 11981309
skid = 11981298
skid = 11908910
skid = 11908663
skid = 11904701

[1] 57 cells

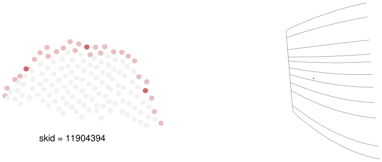
skid = 11904394

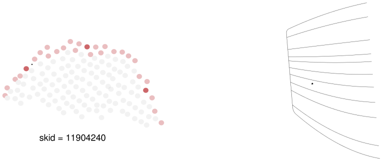
skid = 11904240

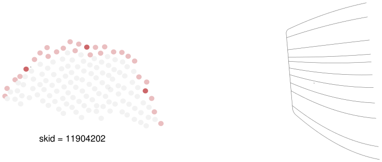
skid = 11904202

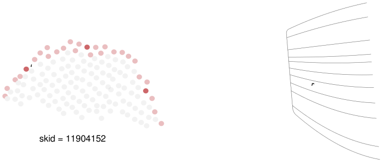
skid = 11904152

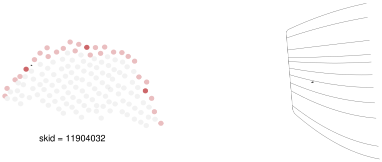
skid = 11904032

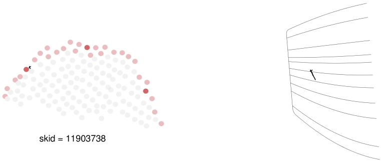
skid = 11903738

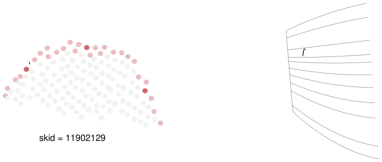
skid = 11902129

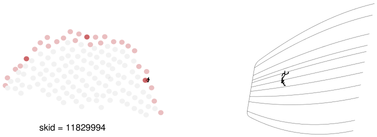
skid = 11829994

[1] 57 cells
