## Supplementary File 3 for "Synaptic targets of photoreceptors specialized to detect color and skylight polarization in *Drosophila*"

### Supplementary File 3: Tables of seed column synapses outside the medulla

#### R7 and R8 outgoing

| Type | No. | pR7 | yR7 | pR8 | yR8 | Sum |
|---|---|---|---|---|---|---|
| Dm9 | 6 | 13 | 9 | 6 | 6 | 34 |
| Dm8 | 15 | 0 | 0 | 0 | 0 | 0 |
| MeTu | 7 | 0 | 0 | 0 | 0 | 0 |
| R7 | 5 | 0 | 0 | 16 | 25 | 41 |
| Tm5c | 6 | 0 | 5 | 2 | 9 | 16 |
| Tm20 | 4 | 0 | 1 | 0 | 1 | 2 |
| Mi15 | 4 | 5 | 5 | 15 | 35 | 60 |
| Mi4 | 4 | 0 | 0 | 0 | 4 | 4 |
| ML1 | 4 | 0 | 0 | 0 | 0 | 0 |
| Dm2 | 4 | 0 | 0 | 0 | 0 | 0 |
| Dm11 | 2 | 16 | 23 | 1 | 4 | 44 |
| L3 | 4 | 15 | 17 | 7 | 10 | 49 |
| Mi1 | 4 | 0 | 0 | 1 | 1 | 2 |
| R8 | 4 | 22 | 24 | 0 | 0 | 46 |
| Tm5a | 2 | 0 | 0 | 0 | 0 | 0 |
| Tm5b | 2 | 0 | 0 | 0 | 0 | 0 |
| Tm | 9 | 0 | 0 | 0 | 0 | 0 |
| Tm5b-like | 3 | 0 | 0 | 0 | 0 | 0 |
| Mi9 | 4 | 0 | 0 | 0 | 0 | 0 |
| L1 | 4 | 2 | 1 | 10 | 7 | 20 |
| aMe12 | 2 | 0 | 0 | 0 | 0 | 0 |
| Dm | 5 | 0 | 0 | 0 | 0 | 0 |
| ML-VPN1 | 3 | 0 | 0 | 0 | 0 | 0 |
| C2 | 2 | 1 | 0 | 4 | 0 | 5 |
| Mt-VPN | 4 | 0 | 1 | 0 | 7 | 8 |
| Mti | 3 | 0 | 0 | 0 | 0 | 0 |
| Tm5a-like | 1 | 0 | 0 | 0 | 0 | 0 |
| TmY10 | 1 | 0 | 0 | 0 | 0 | 0 |
| Mi10 | 1 | 0 | 0 | 0 | 0 | 0 |
| Mi | 1 | 0 | 0 | 0 | 0 | 0 |
| C3 | 1 | 0 | 0 | 0 | 0 | 0 |
| Identified <3 | 26 | 0 | 0 | 2 | 0 | 2 |
| Unidentified >=3 | 2 | 0 | 0 | 0 | 2 | 2 |
| Unidentified <3 | 62 | 1 | 0 | 0 | 3 | 4 |
| Total | 211 | 75 | 86 | 64 | 114 | 339 |

#### R7 and R8 incoming

| Type | No. | pR7 | yR7 | pR8 | yR8 | Sum |
|---|---|---|---|---|---|---|
| Dm9 | 6 | 8 | 4 | 5 | 1 | 18 |
| R8 | 4 | 16 | 25 | 0 | 0 | 41 |
| R7 | 4 | 0 | 0 | 22 | 24 | 46 |
| Mt-VPN | 1 | 0 | 2 | 0 | 3 | 5 |
| C2 | 1 | 0 | 0 | 0 | 0 | 0 |
| L3 | 1 | 0 | 0 | 0 | 0 | 0 |
| Identified <3 | 11 | 1 | 0 | 2 | 5 | 8 |
| Unidentified >=3 | 0 | 0 | 0 | 0 | 0 | 0 |
| Unidentified <3 | 4 | 0 | 2 | 0 | 0 | 2 |
| Total | 32 | 25 | 33 | 29 | 33 | 120 |

#### R7-DRA and R8-DRA outgoing

| Type | No. | R7-DRA | R8-DRA | Sum |
|---|---|---|---|---|
| Dm-DRA1 | 20 | 0 | 0 | 0 |
| Dm9 | 6 | 1 | 1 | 2 |
| MeTu-DRA | 30 | 0 | 0 | 0 |
| R7-DRA | 3 | 0 | 16 | 16 |
| Dm-DRA2 | 9 | 0 | 6 | 6 |
| Dm2 | 4 | 0 | 0 | 0 |
| R8-DRA | 3 | 19 | 0 | 19 |
| Mi15 | 4 | 12 | 8 | 20 |
| Mti-DRA-1 | 6 | 0 | 0 | 0 |
| MeMe-DRA | 2 | 0 | 0 | 0 |
| L3 | 3 | 10 | 8 | 18 |
| VPN-DRA | 6 | 0 | 0 | 0 |
| L1 | 3 | 4 | 3 | 7 |
| Tm20 | 3 | 0 | 0 | 0 |
| Mti-DRA-2 | 4 | 0 | 0 | 0 |
| Mi1 | 3 | 0 | 0 | 0 |
| MeTu | 2 | 0 | 0 | 0 |
| Tm5-like | 1 | 0 | 0 | 0 |
| Mi9 | 2 | 0 | 0 | 0 |
| Dm11 | 1 | 0 | 0 | 0 |
| aMe12 | 1 | 0 | 0 | 0 |
| TmY | 1 | 0 | 0 | 0 |
| ML-VPN2 | 1 | 0 | 0 | 0 |
| C2 | 1 | 2 | 1 | 3 |
| Identified <3 | 33 | 3 | 2 | 5 |
| Unidentified >=3 | 2 | 0 | 0 | 0 |
| Unidentified <3 | 57 | 0 | 0 | 0 |
| Total | 211 | 51 | 45 | 96 |

#### R7-DRA and R8-DRA outgoing

| Type | No. | R7-DRA | R8-DRA | Sum |
|---|---|---|---|---|
| Dm9 | 4 | 0 | 0 | 0 |
| R8-DRA | 3 | 16 | 0 | 16 |
| R7-DRA | 3 | 0 | 19 | 19 |
| C2 | 2 | 3 | 3 | 6 |
| Mi15 | 1 | 2 | 2 | 4 |
| Identified <3 | 9 | 0 | 0 | 0 |
| Unidentified >=3 | 0 | 0 | 0 | 0 |
| Unidentified <3 | 2 | 1 | 1 | 2 |
| Total | 24 | 22 | 25 | 47 |
