## Supplementary File 4 for "Synaptic targets of photoreceptors specialized to detect color and skylight polarization in *Drosophila*"

| Figure(s) | Cross or Genotype | Additional information |
| --- | --- | --- |
| Figure1-Di | pJFRC19-13XLexAop2-IVS-myr::GFP in su(Hw)attP8, pJFRC21-10XUAS-IVS-mCD8::RFP in attP18/ w1118; R21B12-p65ADZp in attP40/ Rh6-nlsLexAGADfl in attP40; R20E07-ZpGDBD in attP2/+ | Split-GAL4 is SS01015<br>anti-dsRed, anti-GFP<br>SlowFade mounting<br>40x objective |
| Figure1-Dii | pJFRC19-13XLexAop2-IVS-myr::GFP in su(Hw)attP8, pJFRC21-10XUAS-IVS-mCD8::RFP in attP18/ w1118; R21B12-p65ADZpattP40/ Rh3-nlsLexAGADfl in attP40 in attP40; R20E07-ZpGDBD_attP2/+ | Split-GAL4 is SS01015<br>anti-dsRed, anti-GFP<br>SlowFade mounting<br>40x objective |
| Figure2-Diii | S003159 (VT043162-p65ADZp in attP40; R20E07-ZpGDBD in attP2) crossed to MCFO-1<br>MCFO stock names are as in Nern et al (2015) Table S2.) | 63x objective<br>DPX mounting<br>anti-Brp (mAb Nc82) reference |
| Figure3-Aii,Aiv | R41C05 crossed to MCFO-7 | 63x objective<br>anti-Brp (mAb Nc82) reference<br>DPX mounting |
| Figure3-Bii,Biv | VT014206 crossed to MCFO-7 | 63x objective<br>anti-Brp (mAb Nc82) reference<br>DPX mounting |
| Figure3-Ciii | SS02425 (R26H07-p65ADZp in attP40; VT010253-ZpGDBD in attP2) crossed to MCFO-1 | 63x objective<br>anti-Brp (mAb Nc82) reference<br>DPX mounting |
| Figure4-Dv | SS02978(R42E06-p65ADZp in attP40; VT064564-ZpGdbd-ZpGDBD in attP2) crossed to MCFO-1 | 63x objective<br>anti-Brp (mAb Nc82) reference<br>DPX mounting |
| Figure4-Dvi | SS02978 crossed to pJFRC51-3XUAS-IVS-Syt::smHA in su(Hw)attP1, pJFRC225-5XUAS-IVS-myr::smFLAG in VK00005 | 63x objective<br>anti-Brp (mAb Nc82) reference<br>DPX mounting |
| Figure5-Ci,Cii | SS28175 (R51E06-p65ADZp in attP40; R15D05-ZpGDBD in attP2) crossed to pJFRC51-3XUAS-IVS-Syt::smHA in su(Hw)attP1, pJFRC225-5XUAS-IVS-myr::smFLAG in VK00005 | overlay of a registered image with the template used for registration<br>63x objective<br>DPX mounting |

| | | |
|---|---|---|
| Figure5-Ciii | SS28175 crossed to MCFO-1 | overlay of a registered image with the template used for registration<br>63x objective<br>DPX mounting |
| Figure5-Civ | SS28175 crossed to MCFO-1 | overlay of a registered image with the template used for registration (the MCFO labeled cells are from the same optic lobe)<br>63x objective<br>DPX mounting |
| Figure5-Cv | ML-VPN1: SS28175 crossed to MCFO-1<br>and L2 neuron terminals in the medulla:<br>SS00801 (R53G02-p65ADZp in attP40; R29G11-ZpGDBD in attP2) crossed to pJFRC51-3XUAS-IVS-Syt::smHA in su(Hw)attP1, pJFRC225-5XUAS-IVS-myr::smFLAG in VK00005 (only synaptotagmin-HA pattern shown) | overlay of registered images showing a single ML-VPN1 cell and L2 terminals in the medulla<br><br>63x objectives<br>DPX mounting |
| Figure5-Cvi | pJFRC19-13XLexAop2-IVS-myr::GFP in su(Hw)attP8, pJFRC21-10XUAS-IVS-mCD8::RFP in attP18/ w1118; R51E06-p65ADZpattP40/ Rh6-nlsLexAGADfl in attP40; R15D05-ZpGDBD_attP2/+ | Split-GAL4 is S28175<br>anti-dsRed, anti-chaoptin (mAb 24B10), native GFP for Rh6 marker<br>40x objective<br>SlowFade mounting |
| Figure5-figure supplement 1 | pJFRC19-13XLexAop2-IVS-myr::GFP in su(Hw)attP8, pJFRC21-10XUAS-IVS-mCD8::RFP in attP18/ w1118; R51E06-p65ADZpattP40/ Rh5-nlsLexAGADfl in attP40; R15D05-ZpGDBD_attP2/+ | Split-GAL4 is S28175<br>anti-dsRed, anti-chaoptin (mAb 24B10), native GFP for Rh5 marker<br>40x objective<br>SlowFade mounting |
| Figure7-Aiv | ortC2b<br>-Gal4;ortC2b<br>-Gal4 crossed to MCFO-1 | 63x objective<br>VECTASHIELD mounting |
| Figure7-Civ | ortC2b<br>-Gal4;ortC2b<br>-Gal4 crossed to MCFO-1 | 63x objective<br>VECTASHIELD mounting |
| Figure8-Aii | R56F07 crossed to MCFO-1 | anti-cadherin (mAb DN-Ex #8) reference<br>63x objective<br>VECTASHIELD mounting |
| Figure8-Avi | R56F07 crossed to MCFO-1 | Photoreceptor neurons labeled with anti-chaoptin (mAb 24B10)<br>anti-cadherin (mAb DN-Ex #8) |

| | | |
|---|---|---|
| | | 63x objective<br>VECTASHIELD mounting |
| Figure8-Bii | OL0007B ((R35D04-p65ADZp in attP40; R65B05-ZpGDBD in attP2) crossed to MCFO-1<br><br>The image stack used is from Wu et al 2016. | 20x objective<br>anti-Brp (mAb Nc82) reference<br>DPX mounting |
| Figure8-Bvi | OL0007B crossed to 20XUAS-CsChrimson-mVenus in attP18 | Photoreceptor neurons labeled with anti-chaoptin (mAb 24B10)<br>40x objective<br>SlowFade mounting |
| Figure8-figure supplement 1 | 20XUAS-CsChrimson-mVenus in attP18/+; R24F06-p65ADZp in attP40/+; VT000770<br>-ZpGDBD in attP2/+ | Single image of this AD/DBD combination from split-GAL4 screening<br>anti-Brp (mAb Nc82) reference<br>20x objective<br>DPX mounting |
| Figure9 -Av | R70E04 crossed to MCFO-7 | anti-Brp (mAb Nc82) reference<br>20x objective<br>DPX mounting |
| Figure9 -Av inset | VT047171 crossed to MCFO-7 | anti-Brp (mAb Nc82) reference<br>63x objective<br>DPX mounting |
| Figure9-Bv | R71A09 crossed to MCFO-7 | anti-Brp (mAb Nc82) reference<br>20x objective<br>DPX mounting |
| Figure9-Cv | R72C08 crossed to MCFO-7 | anti-Brp (mAb Nc82) reference<br>20x objective<br>DPX mounting |
